# Connexinplexity: The spatial and temporal expression of *connexin* genes during vertebrate organogenesis

**DOI:** 10.1101/2021.11.19.469275

**Authors:** Rachel M. Lukowicz-Bedford, Dylan R. Farnsworth, Adam C. Miller

## Abstract

Animal development requires coordinated communication between cells. The Connexin family of proteins is a major contributor to intercellular communication in vertebrates by forming gap junction channels that facilitate the movement of ions, small molecules, and metabolites between cells. Additionally, individual hemichannels can provide a conduit to the extracellular space for paracrine and autocrine signaling. Connexin-mediated communication is well appreciated in epithelial, neural, and vascular development and homeostasis, and most tissues likely use this form of communication. In fact, Connexin disruptions are of major clinical significance contributing to disorders developing from all major germ layers. Despite the fact that Connexins serve as an essential mode of cellular communication, the temporal and cell-type specific expression patterns of *connexin* genes remain unknown in vertebrates. A major challenge is the large and complex *connexin* gene family. To overcome this barrier, we probed the expression of all *connexins* in zebrafish using single-cell RNA-sequencing of entire animals across several stages of organogenesis. Our analysis of expression patterns has revealed that few *connexins* are broadly expressed, but rather, most are expressed in tissue- or cell-type-specific patterns. Additionally, most tissues possess a unique combinatorial signature of *connexin* expression with dynamic temporal changes across the organism, tissue, and cell. Our analysis has identified new patterns for well-known *connexins* and assigned spatial and temporal expression to genes with no-existing information. We provide a field guide relating zebrafish and human *connexin* genes as a critical step towards understanding how Connexins contribute to cellular communication and development throughout vertebrate organogenesis.

## Introduction

Animal development and homeostasis requires coordinated cellular communication. One method mediating communication are gap junction (GJ) channels. GJs are intercellular channels that provide a direct path of low resistance for ionic and small molecule exchange between cells^1^. These channels are formed by the coupling of two apposed hemi-channels each contributed by adjacent communicating cells^1–3^. Additionally, hemi-channels can work independently within a single cell’s membrane, where they can release small molecules such as ATP and glutamate into the extracellular space for paracrine and autocrine signaling^2,3^. The proteins that create GJ channels are evolutionarily unrelated in vertebrates and invertebrates^4^. Yet despite little sequence similarity^5^, the vertebrate Connexins and the invertebrate Innexin proteins have similar structure, with both classes creating four-pass, transmembrane-domain proteins that oligomerize to form each hemichannel within the plasma membrane^4^. Moreover, the hemi-channels and intercellular GJs created by Connexins and Innexins have similar structure and function^4^. Outside of these traditional roles, Connexins can also modulate the formation of tunneling nanotubes, that connect non-adjacent cells to facilitate longer distance communication^6–8^. These varied functions in cellular communication are likely utilized individually and in combination in all animal tissues^9^, yet are best studied in epithelial^10^, neural^11^, and vascular^12^ systems. In these systems, mutations in human Connexin-encoding genes have been linked to defects in the development, regulation, and function including skin disorders^13–16^, cataracts^17,18^, deafness^19–21^, cardiovascular disease^22–24^, and gastrointestinal diseases^25–27^. While Connexin channels serve as an essential form of cellular communication, the temporal and cell-type specific expression patterns of *connexin* genes largely remain unknown.

A major challenge in characterizing *connexin* expression is the complexity of the gene family. In humans, there are 20 distinct *connexin* genes, and in other vertebrate lineages the number of Connexin-encoding genes is similarly large and varies widely^28–30^. Cell culture and *in vitro* work suggests that *connexin* complexity provides functional diversity governed by four general principles: first, hemichannels are created by hexamers of individual Connexin proteins^31^; second, single or multiple Connexin proteins can contribute to hemichannel formation (homo- or heteromeric hemichannels, respectively)^32–34^; third, gap junctions form intercellular channels via hemichannel docking at cell-cell junctions; fourth, each contributed hemichannel can contain the same or different Connexin proteins (homo- or heterotypic channels, respectively)^32–34^. The combinatorial possibilities of the gene family are restrained by molecular engagement rules that limit which Connexins are compatible to form mixed channels^34–37^. These diverse possibilities culminate in each hemichannel having its own unique permeability properties, dependent upon the pore-lining amino acids and channel gating properties of the individual Connexins^37,38^. These rules suggest animals might take advantage of Connexin-based complexity *in vivo* to generate unique functional outcomes, but given the large number of genes, we know little about how vertebrates deploy this gene family.

Most of our knowledge of *connexin* expression *in vivo* comes from only a handful of well-characterized genes. These examples support the idea that *connexins* can be expressed in distinct tissues, such as in mouse where *gap junction a1*/Connexin 43 (*Gja1*/CX43) is expressed extensively in non-neuronal cells, including epithelia^39^, heart^24,40^, and glia^41^. By contrast, *Gjd2*/CX36 is found almost exclusively in neurons^42^. Within the same tissue, *connexin* expression can have distinct temporal patterns, such as *Gjb2*/CX26 and *Gjb1*/CX32 that are both found in the developing mouse neocortex at distinct developmental time points^43^. Within the group of well-studied Connexins, there are also a few enticing examples that suggest the rules of Connexin functional complexity found *in vitro* are relevant to *in vivo* function. For example, heteromeric channels formed by *Gjb1*/CX32 and *Gjb2*/CX26 are found in the mammary gland and the composition of channels changed during development^44^. Heterotypic gap junctions composed of *gjd2a*/Cx35.5 and *gjd1a*/Cx34.1 are found at electrical synapses of zebrafish Mauthner cells where each Connexin was required for the localization of the other in the adjacent cell and both were necessary for synaptic transmission^45,46^. Finally, replacing the coding region of *Gja1*/CX43 with either *Gja5*/CX40 or *Gjb1*/CX32 results in sterility, cardiac malformations and arrhythmias, and mothers unable to nourish their pups, suggesting that each Connexin has unique properties that contribute to cellular homeostasis that cannot be simply interchangeable with other Connexins ^47^. While these examples provide a glimpse of functional complexity, understanding the expression of this gene family through vertebrate development remains as the critical first step to decoding the complexity of *connexin* usage *in vivo*.

Here we set out to examine the expression of all *connexins* in a vertebrate model system, the developing zebrafish, using single-cell RNA-sequencing (scRNA-seq) of cells derived from the entire animal during organogenesis (1-5 days post-fertilization (dpf))^48^. Our analysis of *connexin* expression patterns revealed several trends, including that few *connexins* are broadly expressed, but rather, most *connexins* are spatially restricted to tissue- or cell-type-specific expression patterns. Most cells contain combinatorial signatures of *connexins* with unique profiles within distinct tissues. Finally, *connexin* expression is dynamic with temporal changes across the organism, tissue, and cell type. Our results reveal the complexity of spatiotemporal *connexin* control, highlighting novel aspects of well-studied *connexins* and revealing patterns for *connexin* genes with no prior expression information. We provide a field guide to relate zebrafish and human *connexins* genes, based on evolutionary homologies and expression similarities. Collectively, this represents an important step towards understanding *connexin* gene contributions in cellular communication throughout organogenesis and provides a foundation for comparative analysis in vertebrates.

## Results

### Zebrafish have 41 *connexin* genes

To understand *connexin* expression throughout organogenesis we first set out to ensure the entire *connexin* gene family in zebrafish was identified. Previous efforts^28,49^ and a recent phylogenetic approach to identify the full teleost *connexin* family^30^ captured 40 individual *connexin* genes. Through reciprocal BLAST analysis between the zebrafish genome and (1) human and (2) other teleost Connexin sequences, coupled with phylogenetic analysis, we identified the 40 previously noted *connexins* and one previously unreported *connexin, gjz1*, which is conserved in mammals but forms an outgroup with the rest of the Connexin proteins (Supp Fig. 1, 2).

Across the family of *connexin* genes there are seven human *connexins* for which zebrafish only has a single homolog, eight human *connexins* for which zebrafish has two homologs, two zebrafish *connexins* that have no direct homolog but share sequence similarity to human *connexins*, and sixteen zebrafish *connexins* that are not present in humans but are conserved in other teleost and mammalian lineages^30^. We summarize these relationships in Table 1, listing zebrafish *connexin* genes and their closest relationship with their human counterparts, providing known human and zebrafish expression patterns and phenotypes for comparison. For clarity, we denote *connexins* by their Greek name and by their predicted molecular weight, a naming structure consistent with HUGO^50^ and ZFIN standards^51^ (Table 1, Supp. Table 1). The table is organized to emphasize Connexin similarities based on evolutionary homology, protein sequence, and expression, in alphabetical order of zebrafish *connexin* genes and denotes human similarity across merged rows. There are a limited number of rows where the zebrafish *connexin* gene resembles its human counterpart(s), but the genes are not direct homologs. For example, the human *GJB2/GJB6* genes are duplicated in the human lineage while having only a single similar gene in zebrafish called *gjb8*^30^. Despite not being direct homologs, expression and mutant analyses have found that zebrafish *gjb8* and human *GJB2/GJB6* genes are all involved in inner-ear support cell function and loss of these genes in their respective systems causes deafness^20,52,53^. The comprehensive list of 41 zebrafish *connexin* genes provided a basis to examine the expression patterns of this gene family.

**Table 1.**
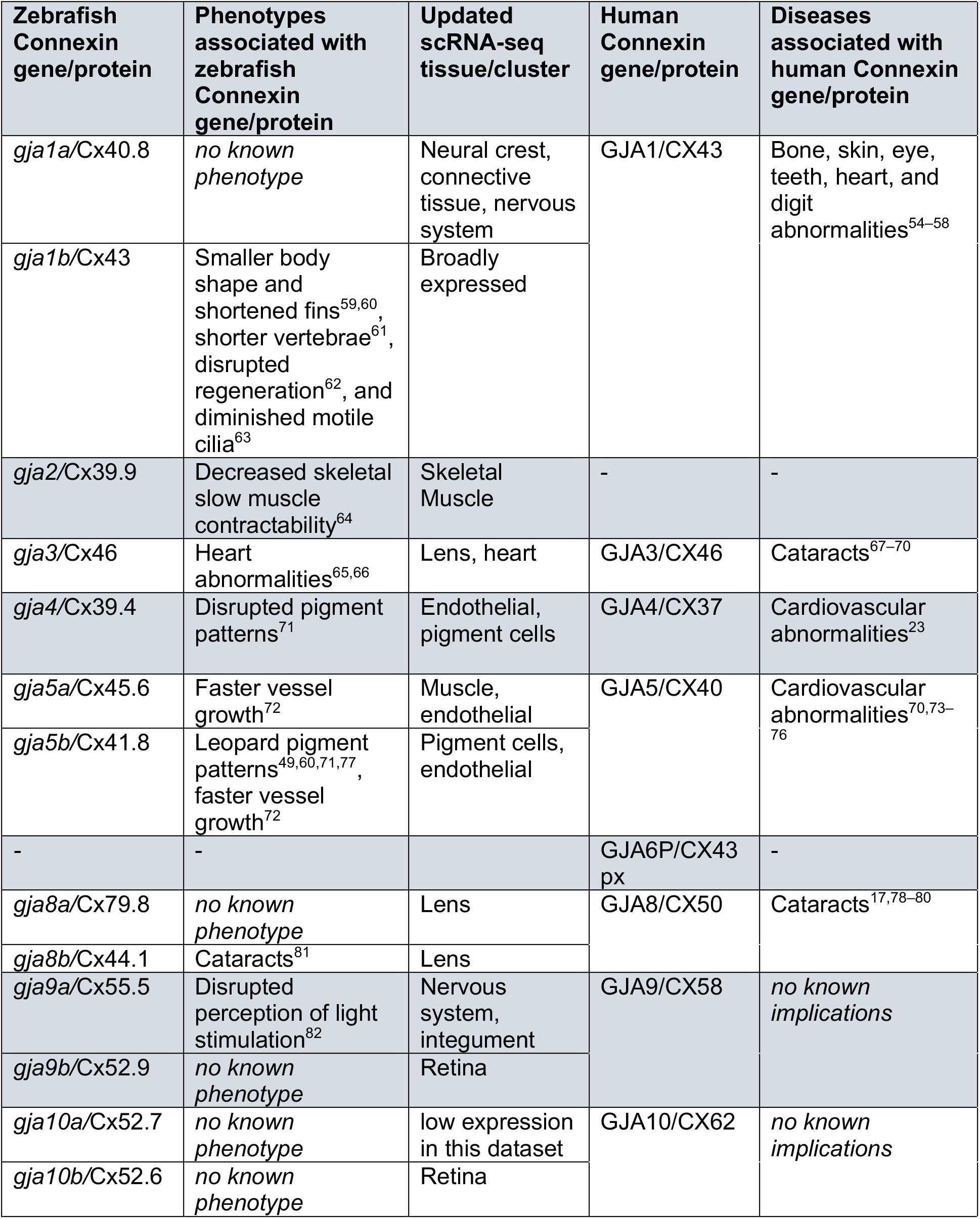
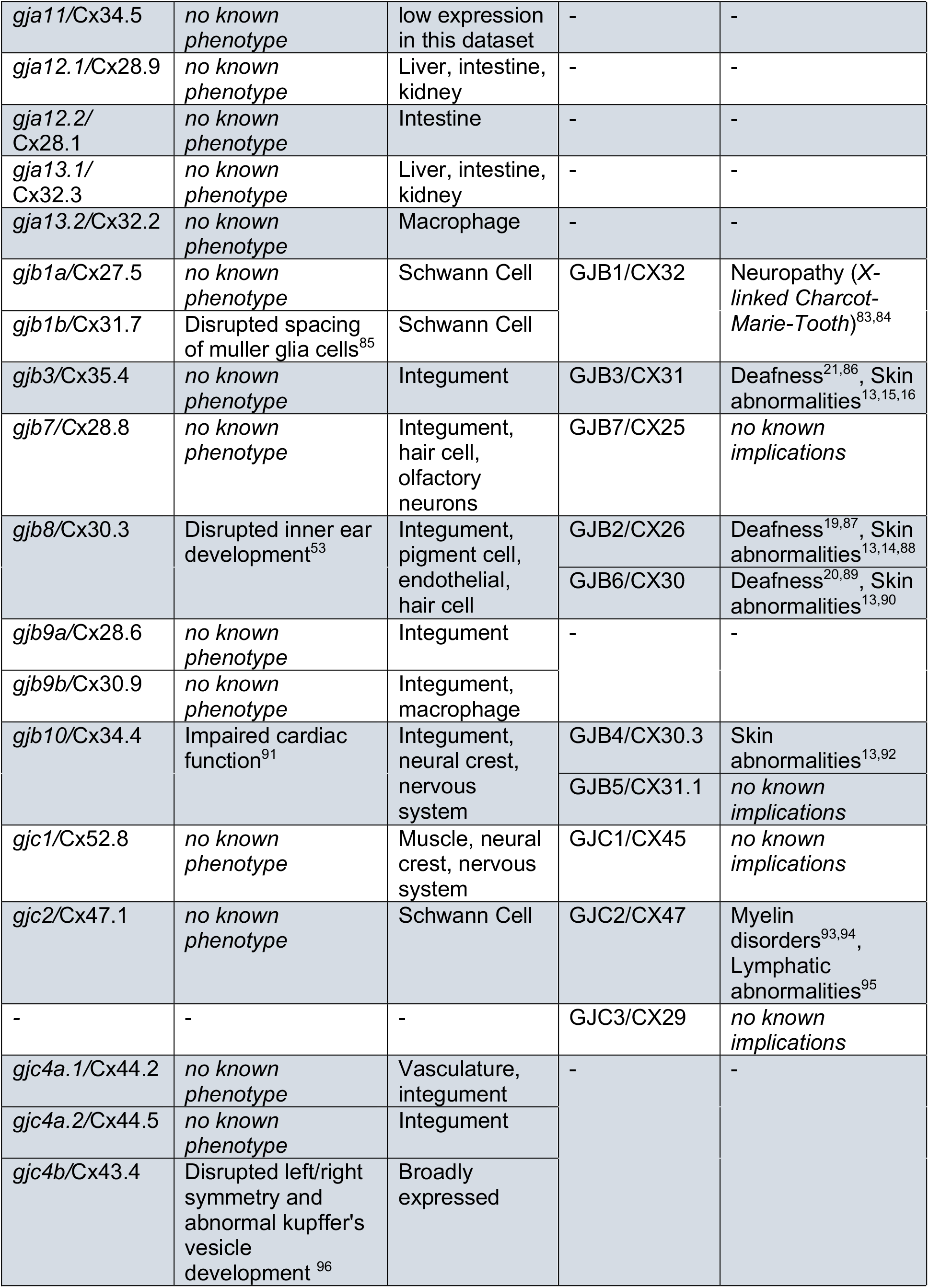
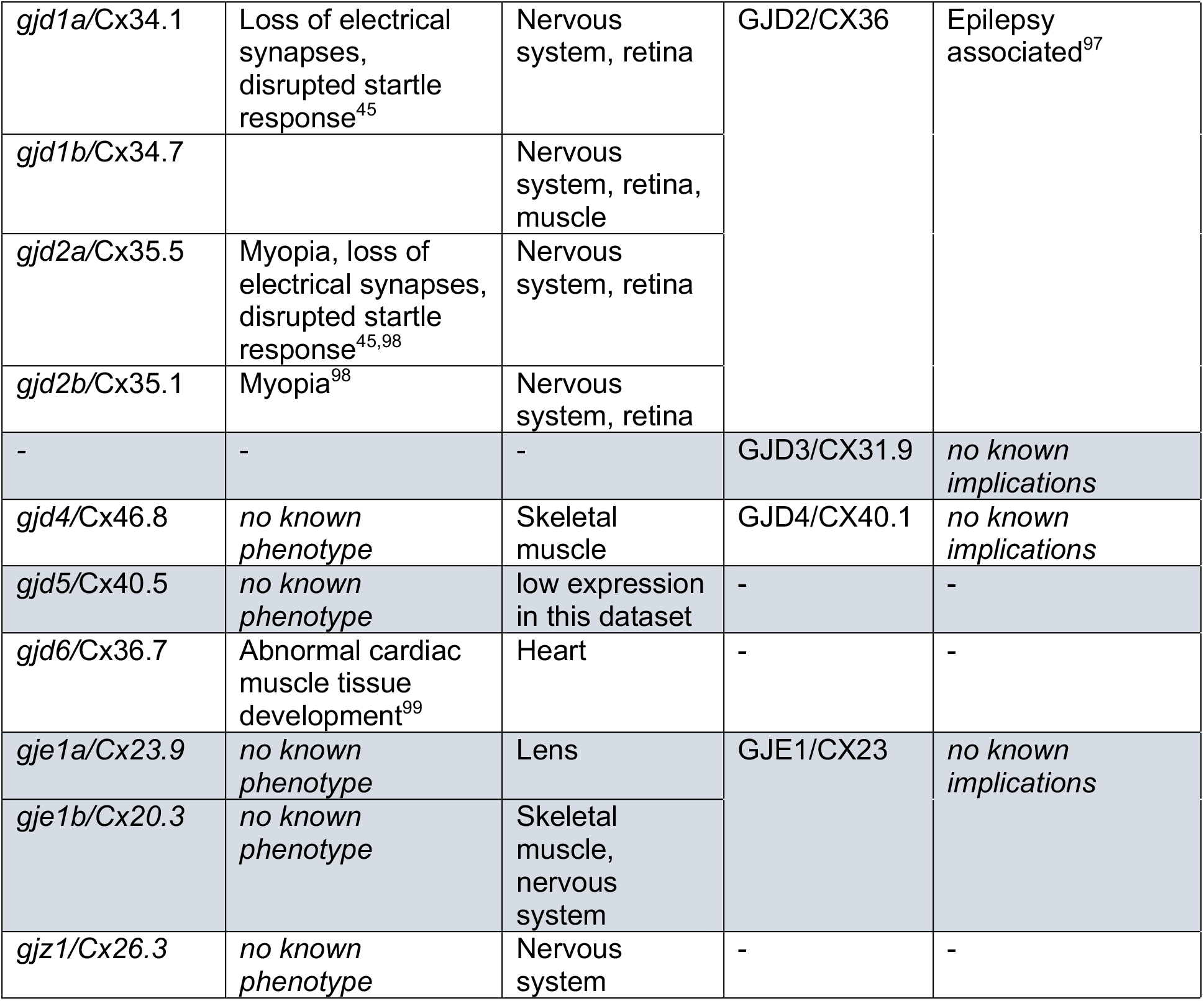
A field guide to zebrafish *connexins*.

| **Zebrafish Connexin gene/protein** | **Phenotypes associated with zebrafish Connexin gene/protein** | **Updated scRNA-seq tissue/cluster** | **Human Connexin gene/protein** | **Diseases associated with human Connexin gene/protein** |
| --- | --- | --- | --- | --- |
| *gja1a*/Cx40.8 | *no known phenotype* | Neural crest, connective tissue, nervous system | GJA1/CX43 | Bone, skin, eye, teeth, heart, and digit abnormalities<sup>54–58</sup> |
| *gja1b*/Cx43 | Smaller body shape and shortened fins<sup>59,60</sup>, shorter vertebrae<sup>61</sup>, disrupted regeneration<sup>62</sup>, and diminished motile cilia<sup>63</sup> | Broadly expressed |  |  |
| *gja2*/Cx39.9 | Decreased skeletal slow muscle contractability<sup>64</sup> | Skeletal Muscle | - | - |
| *gja3*/Cx46 | Heart abnormalities<sup>65,66</sup> | Lens, heart | GJA3/CX46 | Cataracts<sup>67–70</sup> |
| *gja4*/Cx39.4 | Disrupted pigment patterns<sup>71</sup> | Endothelial, pigment cells | GJA4/CX37 | Cardiovascular abnormalities<sup>23</sup> |
| *gja5a*/Cx45.6 | Faster vessel growth<sup>72</sup> | Muscle, endothelial | GJA5/CX40 | Cardiovascular abnormalities<sup>70,73–76</sup> |
| *gja5b*/Cx41.8 | Leopard pigment patterns<sup>49,60,71,77</sup>, faster vessel growth<sup>72</sup> | Pigment cells, endothelial |  |  |
| - | - |  | GJA6P/CX43 px | - |
| *gja8a*/Cx79.8 | *no known phenotype* | Lens | GJA8/CX50 | Cataracts<sup>17,78–80</sup> |
| *gja8b*/Cx44.1 | Cataracts<sup>81</sup> | Lens |  |  |
| *gja9a*/Cx55.5 | Disrupted perception of light stimulation<sup>82</sup> | Nervous system, integument | GJA9/CX58 | *no known implications* |
| *gja9b*/Cx52.9 | *no known phenotype* | Retina |  |  |
| *gja10a*/Cx52.7 | *no known phenotype* | low expression in this dataset | GJA10/CX62 | *no known implications* |
| *gja10b*/Cx52.6 | *no known phenotype* | Retina |  |  |
| *gja11/Cx34.5* | *no known phenotype* | low expression in this dataset | - | - |
| *gja12.1/Cx28.9* | *no known phenotype* | Liver, intestine, kidney | - | - |
| *gja12.2/Cx28.1* | *no known phenotype* | Intestine | - | - |
| *gja13.1/Cx32.3* | *no known phenotype* | Liver, intestine, kidney | - | - |
| *gja13.2/Cx32.2* | *no known phenotype* | Macrophage | - | - |
| *gjb1a/Cx27.5* | *no known phenotype* | Schwann Cell | GJB1/CX32 | Neuropathy (X-linked Charcot-Marie-Tooth)<sup>83,84</sup> |
| *gjb1b/Cx31.7* | Disrupted spacing of muller glia cells<sup>85</sup> | Schwann Cell |  |  |
| *gjb3/Cx35.4* | *no known phenotype* | Integument | GJB3/CX31 | Deafness<sup>21,86</sup>, Skin abnormalities<sup>13,15,16</sup> |
| *gjb7/Cx28.8* | *no known phenotype* | Integument, hair cell, olfactory neurons | GJB7/CX25 | *no known implications* |
| *gjb8/Cx30.3* | Disrupted inner ear development<sup>53</sup> | Integument, pigment cell, endothelial, hair cell | GJB2/CX26 | Deafness<sup>19,87</sup>, Skin abnormalities<sup>13,14,88</sup> |
|  |  |  | GJB6/CX30 | Deafness<sup>20,89</sup>, Skin abnormalities<sup>13,90</sup> |
| *gjb9a/Cx28.6* | *no known phenotype* | Integument | - | - |
| *gjb9b/Cx30.9* | *no known phenotype* | Integument, macrophage |  |  |
| *gjb10/Cx34.4* | Impaired cardiac function<sup>91</sup> | Integument, neural crest, nervous system | GJB4/CX30.3 | Skin abnormalities<sup>13,92</sup> |
|  |  |  | GJB5/CX31.1 | *no known implications* |
| *gjc1/Cx52.8* | *no known phenotype* | Muscle, neural crest, nervous system | GJC1/CX45 | *no known implications* |
| *gjc2/Cx47.1* | *no known phenotype* | Schwann Cell | GJC2/CX47 | Myelin disorders<sup>93,94</sup>, Lymphatic abnormalities<sup>95</sup> |
| - | - | - | GJC3/CX29 | *no known implications* |
| *gjc4a.1/Cx44.2* | *no known phenotype* | Vasculature, integument | - | - |
| *gjc4a.2/Cx44.5* | *no known phenotype* | Integument |  |  |
| *gjc4b/Cx43.4* | Disrupted left/right symmetry and abnormal kupffer's vesicle development<sup>96</sup> | Broadly expressed |  |  |
| *gjd1a/Cx34.1* | Loss of electrical synapses, disrupted startle response<sup>45</sup> | Nervous system, retina | GJD2/CX36 | Epilepsy associated<sup>97</sup> |
| *gjd1b/Cx34.7* |  | Nervous system, retina, muscle |  |  |
| *gjd2a/Cx35.5* | Myopia, loss of electrical synapses, disrupted startle response<sup>45,98</sup> | Nervous system, retina |  |  |
| *gjd2b/Cx35.1* | Myopia<sup>98</sup> | Nervous system, retina |  |  |
| - | - | - | GJD3/CX31.9 | *no known implications* |
| *gjd4/Cx46.8* | *no known phenotype* | Skeletal muscle | GJD4/CX40.1 | *no known implications* |
| *gjd5/Cx40.5* | *no known phenotype* | low expression in this dataset | - | - |
| *gjd6/Cx36.7* | Abnormal cardiac muscle tissue development<sup>99</sup> | Heart | - | - |
| *gje1a/Cx23.9* | *no known phenotype* | Lens | GJE1/CX23 | *no known implications* |
| *gje1b/Cx20.3* | *no known phenotype* | Skeletal muscle, nervous system |  |  |
| *gjz1/Cx26.3* | *no known phenotype* | Nervous system | - | - |

### The *connexin* gene family is broadly expressed, but spatially distinct

Next, we examined the spatio-temporal expression patterns of the zebrafish *connexin* genes through organogenesis using scRNA-seq. We used our recent scRNA-seq atlas dataset in which cells were dissociated from whole embryos at 1, 2, and 5 days post fertilization (dpf), and resultant single-cell expression profiles were captured using the 10X platform^48^. In our initial analysis of the data, we found that many of the *connexin* genes lacked expression information. An examination of the *connexin* gene models generated by Ensembl (GRCz11_93) that were used for mapping single cell reads revealed that most annotations were truncated at or near the end of the protein coding sequence, with most lacking 3’UTRs leading to a failure in capturing the 3’-biased 10X sequencing information (Supp. Fig. 3). To amend this, we used a recently updated gene annotation file that extends gene models^100^, evaluated and updated each *connexin* gene model in reference to bulk RNA-seq data^101^, and imported the Greek gene names. Using this updated gene annotation file, we processed the scRNA-seq data using Cellranger^102^ and evaluated clustering and transcriptional profiles with Seurat^103^. Analysis of the updated scRNA-seq dataset captures transcriptional profiles that appear to represent all major tissues of the developing zebrafish (Fig. 1A*i*, A*ii*) and contains 49,367 cells and 238 clusters. This is 5,355 more cells and 18 more clusters than the original analysis^48^, as expected due to the richer transcriptional information captured from the updated gene model^100^. In our original analysis, we extensively annotated each cluster, assigning the most likely anatomical annotation based on comparing the differentially expressed genes for each cluster to RNA *in situ* patterns^48^. We transferred these previous annotations to our updated analysis by identifying cell-specific barcodes from the original dataset, identifying them in the updated clusters, and transferring the cluster annotations (Fig. 1A*i*, A*ii*; Supp. Table 1,2). As a result, we identified all 220 original clusters^48^ and annotated the remaining clusters by analyzing RNA *in situ* expression information for the most differentially expressed genes (Supp. Table 3). The updated scRNA-seq dataset greatly improves the capture of *connexin* expression throughout the atlas (Supp. Fig. 3), allowing us to examine their spatio-temporal expression pattern during zebrafish organogenesis.

**Figure 1:**
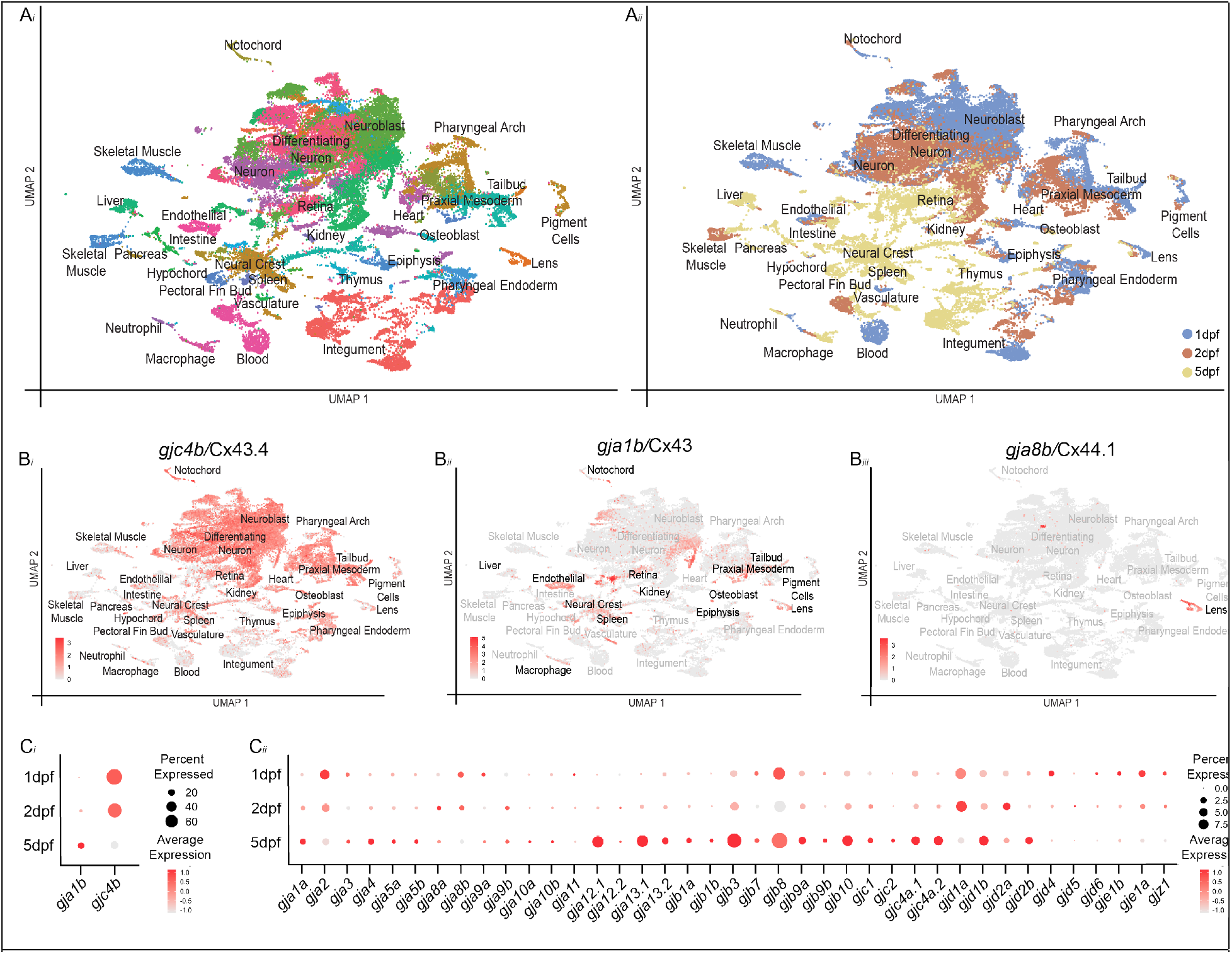
scRNA-seq dataset of zebrafish organogenesis and *connexin* expression. (A*i*) Clustered cell types, where each dot represents a single cell and each color represents a set of transcriptionally related cells. (A*ii*) The age of animals from which cells were dissociated denoted by color – 1 days post fertilization (dpf) cells are blue, 2dpf cells are orange, and 5dpf cells yellow. (B*i*-B*iii*) Expression of well-studied *connexins* in the dataset, where grey represents low expression and red represents the highest level of expression. (B*i*) *gjc4b*/Cx43.4 is expressed broadly across the dataset. (B*ii*) *gja1b*/Cx43 is expressed in a large number of clusters, with notable patterns in liver, endothelial, macrophage, neural crest, spleen, retina, kidney, epiphysis, osteoblast, mesoderm, tailbud, pigment cells and lens clusters. (B*iii*) *gja8b*/Cx44.1 is expressed in lens clusters. (C*i*) Broadly expressed *connexins, gja1b*/Cx43 and *gjc4b*/Cx43.4 and (C*ii*) the remaining *connexin* family shown for each sampled time point. Here, all cells from the cooresponding age are pooled and the percent of cells expressing a given *connnexin* are represented through dot size while the relative expression level is denoted through color intensity.

Using the updated scRNA-seq organogenesis dataset, we examined the expression of each *connexin* related to its clusters, its correlation with marker gene expression, and with cluster annotations (Supp. Fig. 4A-OO, Supp. Table 4). Overall, *connexin* genes had a variety of expression patterns, varying from nearly ubiquitous to cluster-specific and showing a variety of temporal profiles, including constant expression over time or temporal specificity (Fig. 1B,C, Supp. Fig.4 A-OO). To begin to evaluate the dataset’s utility, we first turned our attention to several well-studied *connexin* genes. First, *gjc4b*/Cx43.4 displayed the broadest expression, with particularly high levels in the nervous system, and with diminishing expression from 1 to 5 dpf (Fig. 1B*i*, C*i*; Supp. Fig. 5; Supp. Table 4). This is similar to expression reports for *gjc4b*/Cx43.4 that used RNA *in situ* and transgenic methods^104–106^. *gja1b*/Cx43 is another well-described *connexin*, with broad expression in the cardiovascular system, non-neuronal cells of the retina and central nervous system, mesenchymal cells such as chondrocytes, and within the digestive system including the pancreas^62,107–110^. We find that expression of *gja1b*/Cx43 within the updated clusters largely matches these reported expression patterns (Fig.1 B*ii*; Supp. Fig.4B; Supp. Table 4). We also find expected patterns for *connexins* that have well-known, spatially-restricted expression. For example, *gja8b*/Cx44.1 is expressed almost exclusively in the early developing lens^111–114^, and in the scRNA-seq dataset we find expression of *gja8b*/Cx44.1 within clusters with transcriptional profiles consistent with lens cells (Fig. 1B*iii*; Supp. Figure 5; Supp. Table 4). Further, we find *gja2*/Cx39.9 expression in presumptive skeletal muscle cells, *gjd6*/Cx36.7 specifically in presumptive cardiac muscle, and both *gja9b*/Cx52.9 and *gja10b*/Cx52.6 in presumptive horizontal cells, all well-matching published reports on the expression of these genes^64,99,111,114,115^ (Supp. Figure 5; Supp. Table 4). Taken together, we conclude that the data represented in the updated dataset provides a useful resource for determining the spatio-temporal patterns of *connexin* expression during zebrafish organogenesis.

### *connexins* exhibit complex and combinatorial patterns of expression

To examine the relationship of *connexin* gene expression relative to one another, we organized the scRNA-seq clusters by their tissue annotations and plotted both expression levels and percentage of cells within each cluster (Fig. 2). When arranged in this fashion, the complexity of *connexin* expression within putative tissues and cell types is revealed. In particular, unique combinatorial patterns of *connexins* are observed within tissues developing from all germ layers. For example, within neural clusters (ectoderm), we find that there are four broadly expressed *connexins*, yet each displays bias to either the retina, *gjd1b*/Cx34.7 and *gjd2b*/Cx35.1, or central nervous system, *gjd1a*/Cx34.1 and *gjd2a*/Cx35.5 (Fig. 2; Supp. Figure 4 AF-AI; Supp. Table 4). Within the the skeletal muscle clusters (mesoderm), a unique set of *connexins* are expressed and display a nested hierarchy of expression, with *gja2*/Cx39.9 in all skeletal muscle clusters, *gja5a*/Cx45.6 and *gjd4*/Cx46.8 are restricted to slow muscle clusters, and *gje1b*/Cx20.3 is restricted to fast muscle clusters (Fig. 2; Supp Fig. 4C, JJ, F, NN). We also observed temporally complex patterns of expression. For example, within presumptive intestinal epithelial cells (endoderm), we find that *gjc4b*/Cx43.4 expression diminishes from 1 to 5 dpf, while *gja13.1*/Cx32.3 begins expression at 2 dpf and continues at 5 dpf and *gja12.1*/Cx28.9 becomes co-expressed at 5 dpf (Fig. 2; Supp. Fig. 4Q, O, Supp. Fig. 6). Finally, we observed that primordial germ cells (PGC) express several different *connexins*, including *gja9a*/Cx55.5, *gjb8*/Cx30.3, *gjc4b*/Cx43.4, and *gjd1b*/Cx34.7 (Fig. 2; Supp. Figure 4W, J, GG, EE, Supp. Fig. 7). These observations highlight aspects of the complexity of *connexin* spatial and temporal expression patterns within and across tissues and cell types during zebrafish organogenesis.

**Figure 2:**
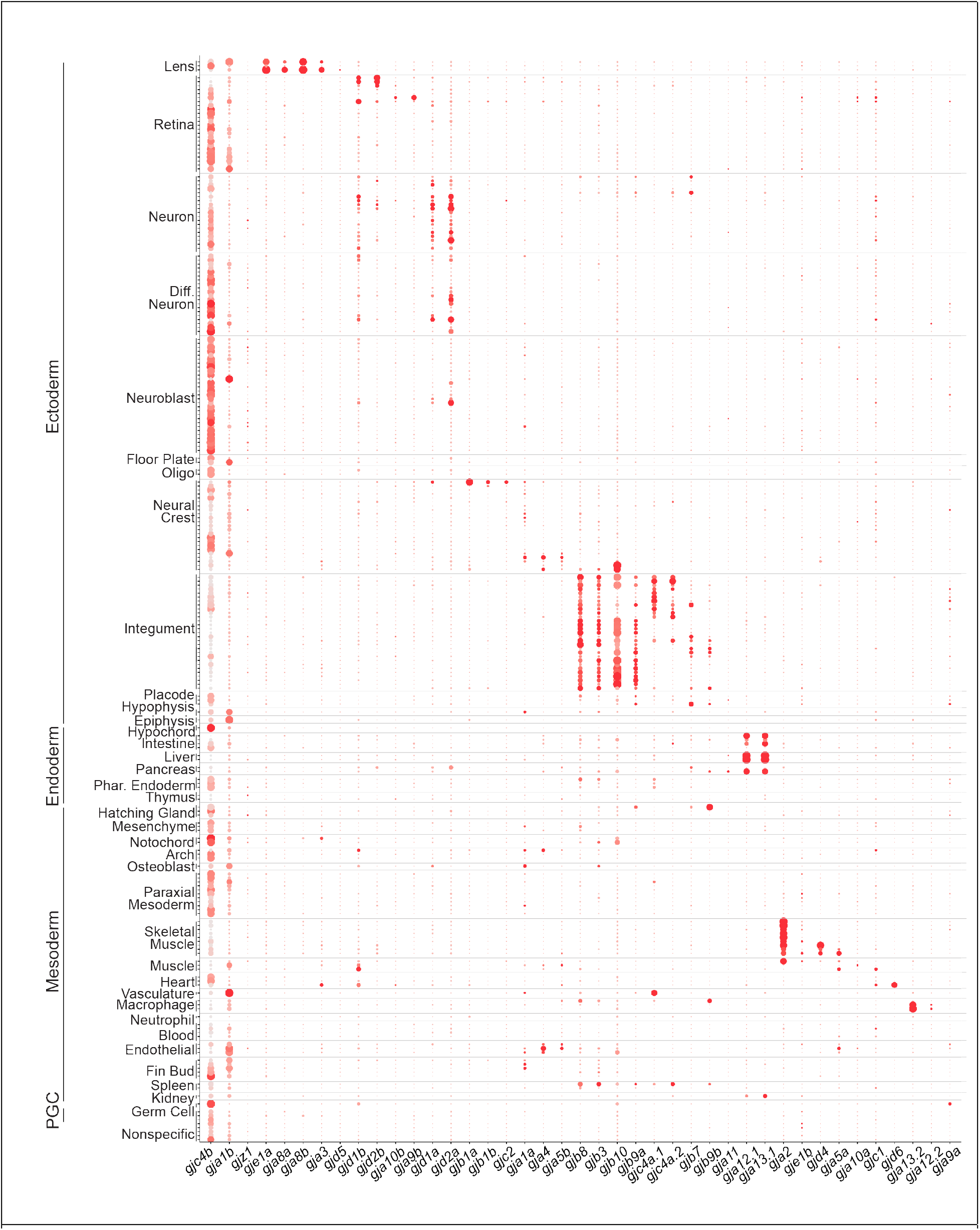
*connexin* expression during zebrafish organogenesis. Clusters are organized by annotations and grouped into tissues and germ layers, denoted on the y-axis. Along the x-axis, *connexins* are arranged based on spatial expression patterns. Each dot represents a single cluster. The percent of cells expressing a given *connnexin* are represented through dot size while the relative expression level is denoted through color intensity. Diff. Neuron = Differentiating Neuron, Oligo = Oligodendrocyte, Phar. Endoderm = Pharyngeal Endoderm, Arch = Pharyngeal Arch, PGC = Primordial Germ Cell

### Cell-type specific expression of *connexins* in the integument *in vivo*

To validate that the *connexin* expression identified in the updated atlas related to *in vivo* tissues and cell types, we examined the integument, or the embryonic skin, as it represented one of the most striking trends of combinatorial expression (Fig. 2). Throughout zebrafish organogenesis, the integument is composed of distinct cellular populations including the periderm (the outermost epidermal layer), the basal cells (a keratinocyte stem cell population), the ionocytes (epithelial cells that maintain osmotic homeostasis), and the pigment cells (neural crest derived cells that provide pigmentation)^116,117^. These individual cell populations are molecularly identifiable using distinct markers including *ppl* (periderm)^104,109^, *tp63* (basal cells)^118^, *foxi3a* (ionocytes)^119^, and *sox10* (pigment cells)^116,120^ (Fig. 3; Supp. Fig. 8). We used these canonical markers in conjunction with our annotations (Supp. Fig. 8; Supp. Table 2) to identify clusters that represent all four cell types of the integument (Fig. 3A*i*). We identified all *connexins* that are significantly expressed within these presumptive integument clusters (Fig. 3; Supp. Fig. 8). We found that *gjb3*/Cx35.4, *gjb8*/Cx30.3, *gjb10*/Cx34.4, and *gjc4b*/Cx43.4 are expressed broadly across these clusters (Fig. 3A*ii*). We then looked for *connexins* enriched in subsets of clusters and found unique and specific patterns of expression. Within periderm clusters, we discovered *gjb9a*/Cx28.6, which has not previously been documented in the skin (Fig. 3A*iii*). Within the presumptive neural crest derived pigment clusters, we found *gja4*/Cx39.4 and *gja5b*/Cx41.8, which are both known to contribute to adult zebrafish skin patterns^49,71^ (Fig. 3A*iv*). Within ionocyte clusters, we identified novel expression for two *connexins*, *gjb7*/Cx28.8 and *gjb9b*/Cx30.9 (Fig. 3A*v*). Finally, within presumptive basal cell clusters, we found novel expression for two *connexins, gjc4a.1*/Cx44.2 and *gjc4a.2*/Cx44.5 (Fig. 3A*vi*). These results suggest the integument uses a complex set of *connexins* throughout organogenesis.

**Figure 3:**
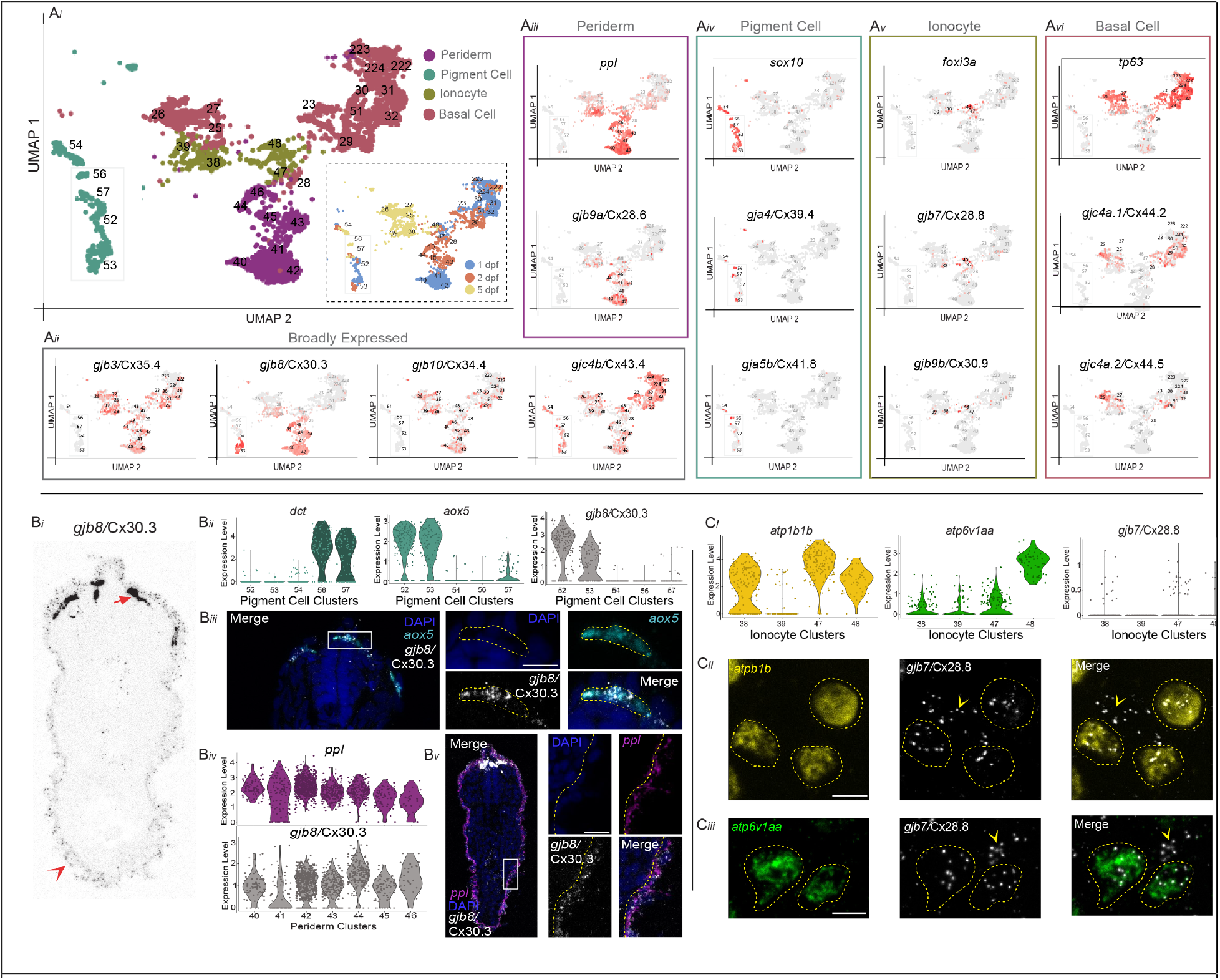
*connexin* expression in the zebrafish integument during organogenesis. (A*i*) The developing integument includes periderm, pigment cells, ionocytes and basal cells. Relevent integument clusters were subsetted from the scRNA-seq dataset. Inset shows the age of animals from which cells were dissociated. (A*ii*) Four *connexins* are broadly expressed in integument clusters, *gjb3*/Cx35.4, *gjb8*/Cx30.3, *gjb10*/Cx34.4, and *gjc4b*/Cx43.4. Grey represents low expression and red represents the highest level of expression. (A*iii*) Periderm marker *ppl* and *gjb9a*/Cx28.6 are expressed in clusters 40-46. (A*iv*) Neural crest derived pigment cell marker *sox10* and *gja4*/Cx39.4 are expressed in clusters 52-57, while *gja5b*/Cx41.8 is only expressed in clusters 54 and 56. (A*v*) Ionocyte marker *foxi3a* and *gjb7*/Cx28.8 are expressed in clusters 38, 39, 47, 48. (A*vi*) Basal cell marker *tp63* and *gjc4a.1*/Cx44.2 are expressed in clusters 23, 25-32, 51, 222-224, while *gjc4a.2*/Cx44.5 is only expressed in clusters 25-29. (B*i*) Fluorescent RNA *in situ* for *gjb8*/Cx30.3 in a transverse cross section of a 1 dpf zebrafish embryo, contrast is inverted for clarity. Dorsal is up, section is from the trunk. Strong expression of *gjb8*/Cx30.3 in neural crest cells is denoted with arrow and weaker, but distinct, periderm expression is denoted with arrowhead. (B*ii*) Within the pigment cell clusters the melanocyte marker *dct* is expressed in clusters 56 and 57, whereas xanthophore marker *aox5* is primarly expressed in clusters 52 and 53. *gjb8*/Cx30.3 is predominantly expressed in clusters 52 and 53. (B*iii*) Transverse cross section of a 1 dpf zebrafish embryo stained with DAPI (blue) and fluorescent RNA *in situ* against *aox5* (cyan) and *gjb8*/Cx30.3 (white), with white box denoting the zoomed panels at the right. Scale bar = 10 uM. (B*iv*) Expression of *ppl* and *gjb8*/Cx30.3 within the periderm clusters. (B*v*) Transverse cross section of a 1 dpf zebrafish embryo stained with DAPI (blue) and fluorescent RNA *in situ* against *ppl* (purple) and *gjb8*/Cx30.3 (white) with white box denoting the zoomed panels at the right. (C*i*) Within the ionocyte clusters the Na+,K+-ATPase-rich cell and H+-ATPase-rich cell markers *atp1b1b* and *atp6v1aa*, respectively, are expressed in conjunction with low expression of *gjb7*/Cx28.8. (C*ii*) Fluorescent RNA *in situ* in a 1dpf zebrafish embryo against *atp1b1b* (yellow), *gjb7*/Cx28.8 (white), with merged signal (right). *atp1b1b* expressing cells are outlined with a dashed yellow line, and *gjb7*/Cx28.8 signal outside of those cells are marked with yellow arowhead. Scale bar = 10 uM. (C*iii*) Fluorescent RNA *in situ* in a 1dpf zebrafish embryo against *atp6v1aa* (green), *gjb7*/Cx28.8 (white), with merged signal (right). *atp6v1aa* expressing cells are outlined with a dashed yellow line, and *gjb7*/Cx28.8 signal outside of those cells are marked with yellow arowhead. Scale bar = 10 uM.

We next examined a subset of the identified integument *connexins in vivo*. We first tested a broadly expressed *connexin, gjb8*/Cx30.3, to see if it was expressed in the pigment cells and periderm using fluorescent RNA *in situ* on 1 dpf embryos. Transverse cross sections through the trunk revealed prominent *gjb8*/Cx30.3 staining in dorsally located cells near the neural tube and additional dim staining was observed in a single layer of cells surrounding the entire embryo (Fig. 3B*i*). We first confirmed *gjb8*/Cx30.3’s expression in pigment cells by subsetting the five clusters that appear to represent pigment cells, including melanophores^121,122^ (*dct*+, clusters 56, 57, Fig. 3 B*ii*; Supp. Table 2) and xanthophores^123^ (*aox5*+, clusters 52, 53, Fig. 3 B*ii*; Supp. Table 2). We find that *gjb8*/Cx30.3 is highly expressed in only the presumptive xanthophore clusters (Fig. 3 B*ii*). We then performed fluorescent RNA *in situ* for *aox5* and *gjb8*/Cx30.3 in a 1 dpf embryo and found robust co-localization of these transcripts, confirming that *gjb8*/Cx30.3 is expressed in xanthophore cells (Fig. 3B*iii*). We then examined *gjb8*/Cx30.3’s expression in the periderm through sub-setting the seven presumptive periderm clusters (*ppl*+, clusters 40-46, Supp. Table 2) and find expression of *gjb8*/Cx30.3 in all clusters (Fig. 3 B*iv*). Indeed, fluorescent RNA *in situ* for *ppl* and *gjb8*/Cx30.3 reveal robust co-localization of these transcripts in the outermost epithelial layer (Fig. 3B*v*), confirming that *gjb8*/Cx30.3 is expressed in the developing periderm.

We next tested a *connexin* with more specific expression within the integument clusters, *gjb7*/Cx28.8, which has expression specific to the presumptive ionocytes (Fig. 3A*v*). Developing *foxi3a*+ ionocytes form Na+,K+-ATPase-rich (NaR) cells or H+-ATPase-rich (HR) cells, which are characterized by the expression of the specific ATPase genes *atp1b1b* and *atp6v1aa*, respectively^119^. First, we sub-setted all ionocyte clusters^119^ (*foxi3a*+, 38, 39, 47, 48) and found unique expression combinations of *atp1b1b* and *atp6v1aa* across clusters and low expression of *gjb7*/Cx28.8 in 3 of 4 clusters (Fig. 3C*i*). Fluorescent RNA *in situ* revealed co-localization of *gjb7*/Cx28.8 with both *atp1b1b* (Fig. 3C*ii*) and *atp6v1aa* (Fig. 3C*iii*), confirming that *gjb7*/Cx28.8 is expressed in ionocytes. Together, these data confirm the predictive power of the scRNA-seq dataset for *connexin* expression in the integument and support the utility of the dataset as a novel tool for discovery of investigating *connexin* complexity in vertebrate development.

## Discussion

Here we reveal the details of *connexin* gene-family expression during zebrafish organogenesis showing that *connexin* usage is widespread yet displays gene-specific variations across tissue, cell type, and developmental time. The large gene family of *connexins* in zebrafish (41 genes) is expressed in complex patterns ranging from nearly ubiquitous to celltype specific, with unique combinatorial and nested expression sets restricted to individual tissues. Temporally, *connexins* display sustained, increasing, and diminishing expression profiles across development, dependent upon gene and tissue. Together, these data reveal the complexity of expression of this critical gene family in a model vertebrate and demonstrate that this critical form of communication is likely to be used by all tissues during organogenesis. These data provide a critical framework facilitating analysis of how these genes contribute to cellular communication in tissues developing from all germ layers, providing a basis to understand *connexins* in development and in modeling human disease.

We find that all cells express *connexins*, but each tissue expresses a unique combinations of the gene family with the composition of the expressed set evolves over developmental time. This spatiotemporal complexity of *connexin* family usage likely contributes to both functional redundancy within tissues as well as functional diversity. The many *connexins* expressed might allow for a myriad of combinatorial interactions amongst Connexin proteins, which could contribute to heteromeric hemichannels and heterotypic GJs. Importantly, Connexins can only interact with potential partners if they are expressed in the same cell or between interacting cells, thus the work here constrains the combinatorial problem of complex usage by revealing the details of the expression patterns through organogenesis. For example, *gjd2a*/Cx35.5 and *gjd1a*/Cx34.1 have been shown to form heterotypic GJs (unique Connexins on each side of the GJ) at electrical synapses of the Mauthner cell neural circuit^45^. The data here shows extensive overlapping expression of these two *connexins* throughout the central nervous system, suggesting complex hemi-channels and GJs could be common throughout the brain. Given that each Connexin-mediated hemichannel has its own unique set of compatibilities and permeability properties, this dataset provides a platform for future research to explore whether *connexins* expressed within the same tissue or cell type form functional channels, and how the molecular identity of these channels influences function.

This dataset presents as a powerful resource for zebrafish and *connexin* biology. We establish *connexin* expression in cells previously unknown to express *connexins*, such as the ionocytes of the skin. Within our dataset there are numerous other cell types with striking *connexin* expression patterns that have under-appreciated *connexin* usage inviting exploration, including macrophages (*gja13.2*/Cx32.2) and primordial germ cells (*gja9a*/Cx55.5, *gjb8*/Cx30.3, *gjc4b*/Cx43.4 and *gjd1b*/Cx34.7). Another strength of this dataset is the exploration of expression across multiple cell types, tissues, and timepoints simultaneously. For example, *gja3*/Cx46 has only been examined in the heart ^65,66^, yet, in our dataset we find robust *gja3*/Cx46 expression in both heart and lens clusters, which suggests an enticing link to human GJA3/CX46, in which mutations are associated with cataracts^67–69^. Finally, this dataset provides putative expression to many *connexin* genes that had no previous expression information (22/41 genes). For example, *gjb1a*/Cx27.5 and *gjc2*/Cx47.1 are both highly expressed in the Schwann cell cluster. While neither of these genes had previously known expression information, mutations of their human orthologs GJB1/CX32 and GJC2/CX47 contribute to neuropathy and myelin disorders^86,93,94^. The identification of tissues and cell type expression patterns for the entire gene family creates a basis to explore *connexin* related diseases in zebrafish and provide comparisons to human biology. Through exploring the *connexin* family expression across diverse cell types and tissues, we can begin to envision a holistic view of Connexins utilization and usage in cellular communication throughout organogenesis.

## Methods

### scRNA-seq

#### Embryo dissociation and cDNA library prep

As described by Farnsworth et al., 2020^48^, larvae from the *Tg(olig2:GFP)vu12* and *Tg(elavl3:GCaMP6s)* backgrounds were pooled (n=15 per replicate), with 2 replicates at each sampled timepoint (1, 2, 5dpf). Cells from entire larvae were dissociated using standard protocols. Dissociated cells were then run on a 10X Chromium platform using 10x v.2 chemistry aiming for 10,000 cells per run.

#### Alignment

To ensure that the full transcripts of the Connexin-encoding genes were represented in the dataset, we used gene models with lengthened 3’ UTRs across the zebrafish genome generated and validated by the Lawson Lab^100^. We ensured that the *connexin* genes were annotated properly by comparing pooled deep-sequencing information and extended the 3’ UTR regions as needed. Using this updated GTF file, we aligned reads to the zebrafish genome, GRCz11_93, using the 10X Cellranger pipeline (version 3.1).

#### Computational Analysis

Cells were analyzed using the Seurat (V3.1.5) software package for R (V4.1.0) using standard quality control, normalization, and analysis steps. We performed PCA using 115 PCs based on a Jack Straw-determined significance of P < 0.01. UMAP analysis was performed on the resulting 49,367 cells with 115 PC dimensions and a resolution of 15.0, which produced 238 clusters.

#### Cluster annotation

The unique barcode assigned to each cell was extracted from the original Farnsworth dataset^48^ and identified in our updated dataset. For each updated cluster, we analyzed the percentage of cells contributing which were associated with the original Farnsworth’s clusters. Frequently, we found the updated dataset contained clusters with a significant proportion of cells (>80%) from a single Farnsworth cluster, and in such we transferred the annotation from the original cluster to the updated cluster. We also found instances of a single Farnsworth cluster breaking nearly evenly across two of the updated clusters – for example, the original dataset had a single ‘photoreceptor’ cluster (cluster 115), whereas the updated data had two clusters (clusters 13, 14) with cells from original photoreceptor cluster. Further analysis revealed that these two new clusters represented likely rods and cones. Finally, we also found updated clusters that did not have a clear previous annotation. In these instances, we analyzed the most differentially expressed genes from that cluster and compared them with canonical markers.

#### Fluorescent RNA in-situ

Custom RNAscope probes to target *connexin* genes were designed and ordered through ACD (https://acdbio.com/). For the fluorescent *in situs*, we used a modified RNAscope protocol^124^. Briefly, 1 dpf embryos were fixed for 2 hours at room temp in 4% PFA and then stored in 100% methanol at −20C. The tissue was then exposed to protease plus for 30 min, washed with PBS with 1% Triton X (PBSTx), and then hybridized with the 1x probe overnight at 40C. Standard RNAscope V2 multiplex reagents and Opal fluorophores were used, with the modification that PBSTx which was used for all wash steps. Stained tissue was either mounted (whole mount) or immediately cryo-sectioned and mounted with ProLong Gold Antifade (ThermoFisher).

#### Zebrafish Husbandry

Fish were maintained by the University of Oregon Zebrafish Facility using standard husbandry techniques^125^. Embryos were collected from natural matings, staged, and pooled. Animals used in the original Farnsworth data were: *Tg(olig2:GFP)vu12* and *Tg(elavl3:GCaMP6s)*^48^, and animals used for RNAscope *in situs* were *ABC-WT*. Animal use protocol AUP-18-35 was approved by the University of Oregon IACUC committee and animal work was overseen by Dr. Kathy Snell.

## Acknowledgments

We thank the entire Miller Lab ongoing support, comments, and discussions on this manuscript. We thank Clay Small for discussions and expertise in regards to data handling, statistics, and annotation transfer from the original to updated atlas. We thank the University of Oregon AqACS facility for superb animal care. We thank the ZFIN team, and Svein-Ole Mikalsen, for communication regarding the *connexin* gene names. This work was supported by the NIH National Institute of General Medical Sciences, Genetics Training Grant T32GM007413 to R.M.L., and the NIH Office of the Director R24OD026591 and the NIH National Institute of Neurological Disorders and Stroke R01NS105758 to A.C.M.

## Data availability

All data generated or analyzed during this study are included in the published article and its supplementary information files. Sequences used in this study were deposited to the NCBI SRA and can be found using the identifier PRJNA564810. Additional files, including the updated GTF and analysis, can be found at https://www.adammillerlab.com/.

## Supplemental Figure Legends

Supplemental Figure 1: Phylogeny of human (*hs*) and zebrafish (*dr*) Connexin proteins.

Supplemental Figure 2: Protein similarities and phylogeny of the novel *gjz1*/Cx26.3. (A) Zebrafish (*dr*.) Gjz1/Cx26.3 aligned with the most similar human (*hs*.) Connexin protein, GJB3. The predicted intercellular portions (coral), transmembrane domains (green) and extracellular loops (blue) for GJB3 are denoted above the sequence. (B) Ensembl-generated phylogenetic tree of *gjz1*/Cx26.3 (si:rp71-1c10.10 in red). Bony fishes are highlighted in pink, while lobed-fin lineages from coelacanth to mammals are in other colors. Protacanthopterygii and Euacanthomorphacea sub-trees are collapsed for visual purposes (grey triangle). Each related gene is represented on a node and colored based on Ensembl predications, including a gene node (white box), speciation nodes (dark blue box), duplication nodes (red box), and ambiguous nodes (teal box). Genomic sequence alignment similarity is denoted on the right with black boxes (representing 66-100% alignment), moderate sequence alignment is denoted on the right with green boxes (representing 33-66% sequence alignment) and gaps in alignments are denoted in white.

Supplemental Figure 3: Gene model updates to capture *connexin* expression. (A) Aligned sequencing view showing coverage pileups for scRNA-seq (1, 2, 5 dpf) and bulk RNA-seq, as well as bulk RNA-seq reads (grey boxes). The Ensembl GTF (light green) captures few reads associated with *gjd4*/Cx46.8. Extending the gene model (dark green) in the updated GTF captures these missed transcripts. (B) Comparisons between the Farnsworth et al., 2020 dataset and the updated dataset. (B*i*) Farnsworth cluster 57 has robust expression of slow muscle marker *smyhc2*, but poor expression of *gjd4*/Cx46.8. In the updated dataset, the related cluster 4 has similar expression of slow muscle marker *smyhc2* but a significant increase of *gjd4*/Cx46.8 representation (right). (B*ii*) Farnsworth cluster 205 has robust expression of cardiac muscle marker *myl17* and *gjd6*/Cx36.7. The corresponding cluster in the updated dataset did not significantly alter either of these expression patterns (right). (B*iii*) Farnsworth cluster 173 has low expression of Schwann Cell marker, *mbpa*, as well as *gjb1a*/Cx27.5. The corresponding cluster in the updated dataset captures robust expression for both *mbpa* and *gjb1a*/Cx27.5.

Supplemental Figure 4: *connexin* expression throughout the atlas. Note that this figure extends across 41 pages, one for each *connexin* gene, labeled A-OO. Expression of each *connexin* is plotted on the scRNAseq atlas and visualized through color intensity on UMAP plots and by violin plots for each gene and cluster.

Supplemental Figure 5: Tissue and cell type markers with corresponding *connexin* expression. (A*i*) Central nervous system clusters express *snap25a* and *pcna*, and *gjc4b*/Cx43.4. (A*ii*) Lens clusters (193, 112 and 237) express lens markers *crybb1* and *crybab2*, and *gja8b*/Cx44.1. (A*iii*) Skeletal muscle clusters express slow muscle marker *smyhc1* (4, 5, 234), fast muscle marker *myhz2* (7, 8, 9, 10, 11, 233), and *gja2*/Cx39.9. (A*iv*) Cardiac muscle cluster 233 expresses markers *nppa* and *myl17*, and *gjd6*/Cx36.7. (A*v*) Retinal horizonal neurons, cluster 203, express marker *mdka* and *gja9b*/Cx52.9 and *gja10b*/Cx52.6.

Supplemental Figure 6: Temporal expression patterns of *connexins* within the intestine. (A*i*) Intestinal clusters (3, 60, 67, 140) selected for expression of canonical markers *cldnc, fabp2*, and *foxa3*. (A*ii*) These clusters display an evolution of *connexin* expression at different developmental time points. *gjc4b*/Cx43.4 (left) is expressed at 1 and 2 dpf, *gja13*.1/Cx23.3 is expressed at 2 and 5 dpf, and *gja12*.1/Cx28.9 is expressed at 5 dpf.

Supplemental Figure 7: Primordial germ cells express several *connexins*. Putative primordial germ cell (PGC) cluster (128) expressing PGC markers like *ddx4* and *nanos3*, in addition to *connexins* including *gja9a*/Cx55.5, *gjb8*/Cx30.3, *gjc4b*/Cx43.4, and *gjd1b*/Cx34.7.

Supplemental Figure 8: *connexin* expression in the integument. Clusters classified into cell types are grouped and labelled accordingly. (A*i*) *gjb3*/Cx35.4, *gjb8*/Cx30.3, *gjb10*/Cx34.4, and *gjc4b*/Cx43.4 are all expressed broadly through the integument clusters. (A*ii*) *ppl*, *krt4*, and *evpla* are expressed highly in periderm clusters, with *gjb9a*/Cx28.6 being primarily expressed in periderm clusters. (A*iii*) *sox10* and *aox5* expression identifying pigment cell clusters with *gja4*/Cx39.4 and *gja5b*/Cx41.8. (A*iv*) *foxi3a* expression identifying ionocyte clusters with *gjb7*/Cx28.8 and *gjb9b*/Cx30.9. (A*v*) *tp63* expression identifying basal cell clusters with *gjc4a*.1/Cx44.2 and *gjc4a*.2/Cx44.5.

Supplemental Table 1: The zebrafish *connexin* family.

Supplemental Table 2: Transferring Farnsworth et. al, 2020 cluster annotations to the updated dataset. Updated cluster number (Column A) and the corresponding Farnsworth cluster (Column B). The count of cells from the Farnsworth cluster that ended in the corresponding updated cluster (Column C), and the proportion of cells from a given Farnsworth cluster that contribute to the updated cluster (Column D, E). Total cell counts are colored in blue and bolded. All previous Farnsworth annotations were transferred over (columns H-AE), and update cluster markers are included in orange. The most significant Farnsworth cluster contributor is denoted in black font.

Supplemental Table 3: List of differentially expressed genes for each cluster. (Sheet1) For each cluster (Column A), annotations at germ layer, tissue, cell type, and subtype (Columns B – E) level are listed. For each cluster, the top 16 most differentially expressed genes are listed (Column F - U). (Sheet 2) All differentially expressed genes for each updated cluster, generated using the FindAllMarkers command of Seurat, using the Wilcoxon rank sum test. Pct.1 (Column D) and Pct.2 (Column E) reflect the fraction of cells expressing each marker gene (Boolean) within each cluster and for all other cells, respectively.

Supplemental Table 4: The proportion of cells within each cluster that express each *connexin*.

Supplemental Table 5: All reagents used for fluorescent RNA in-situ and other immunohistochemistry in this study.

