## Supplemental Figures for "Connexinplexity: The spatial and temporal expression of *connexin* genes during vertebrate organogenesis"

Supplemental Figure 1

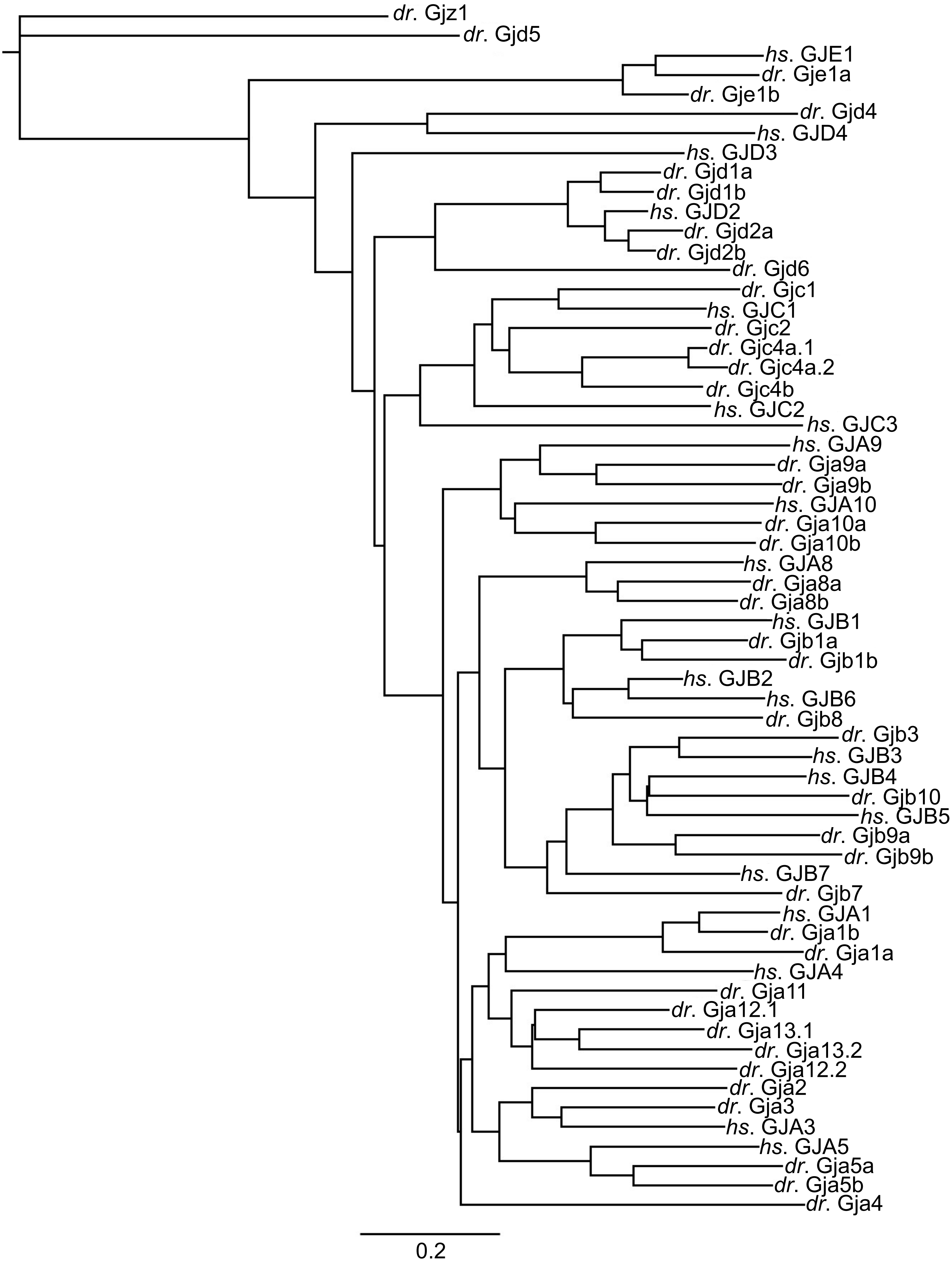

Supplemental Figure 2

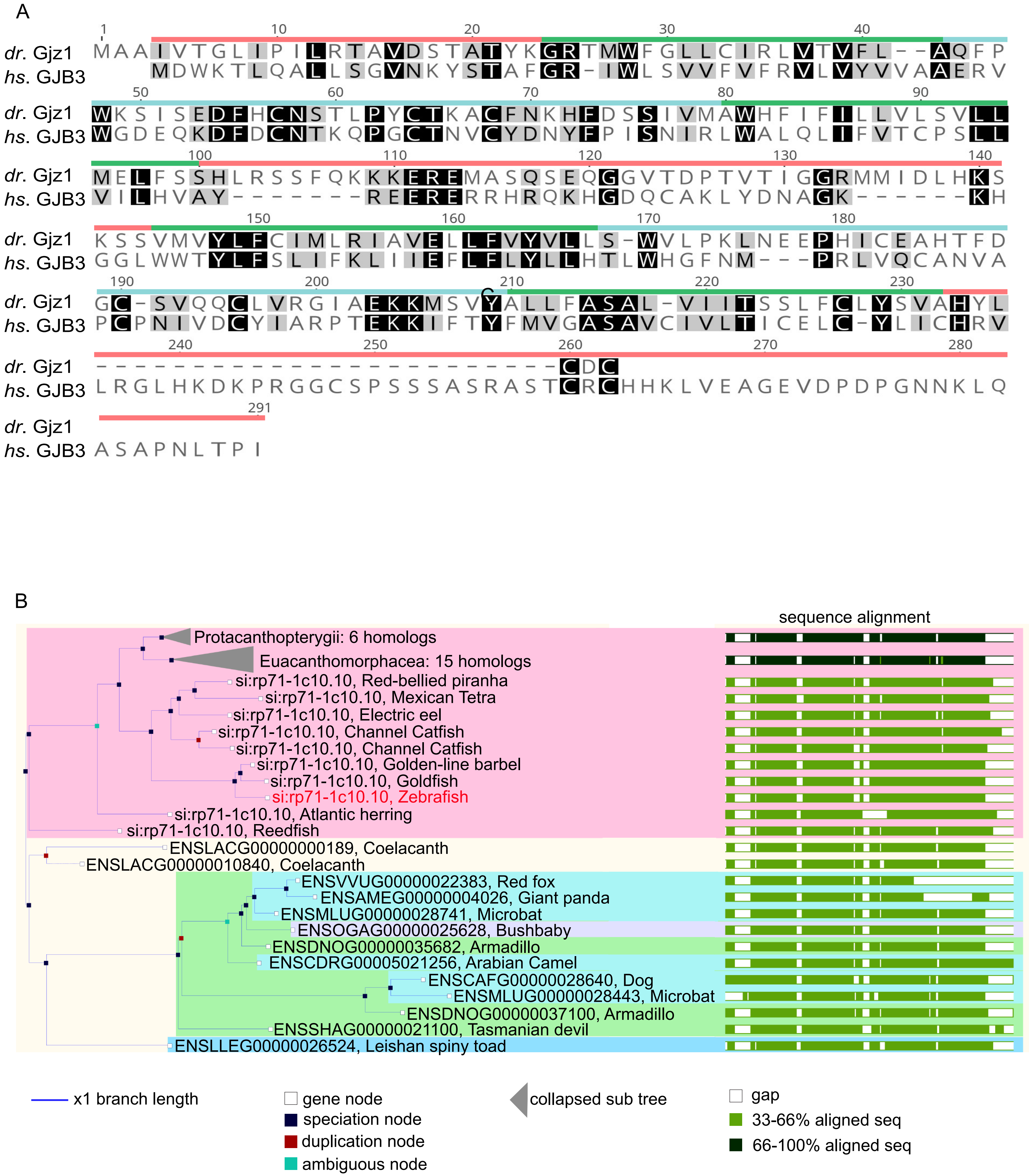

Supplemental Figure 3

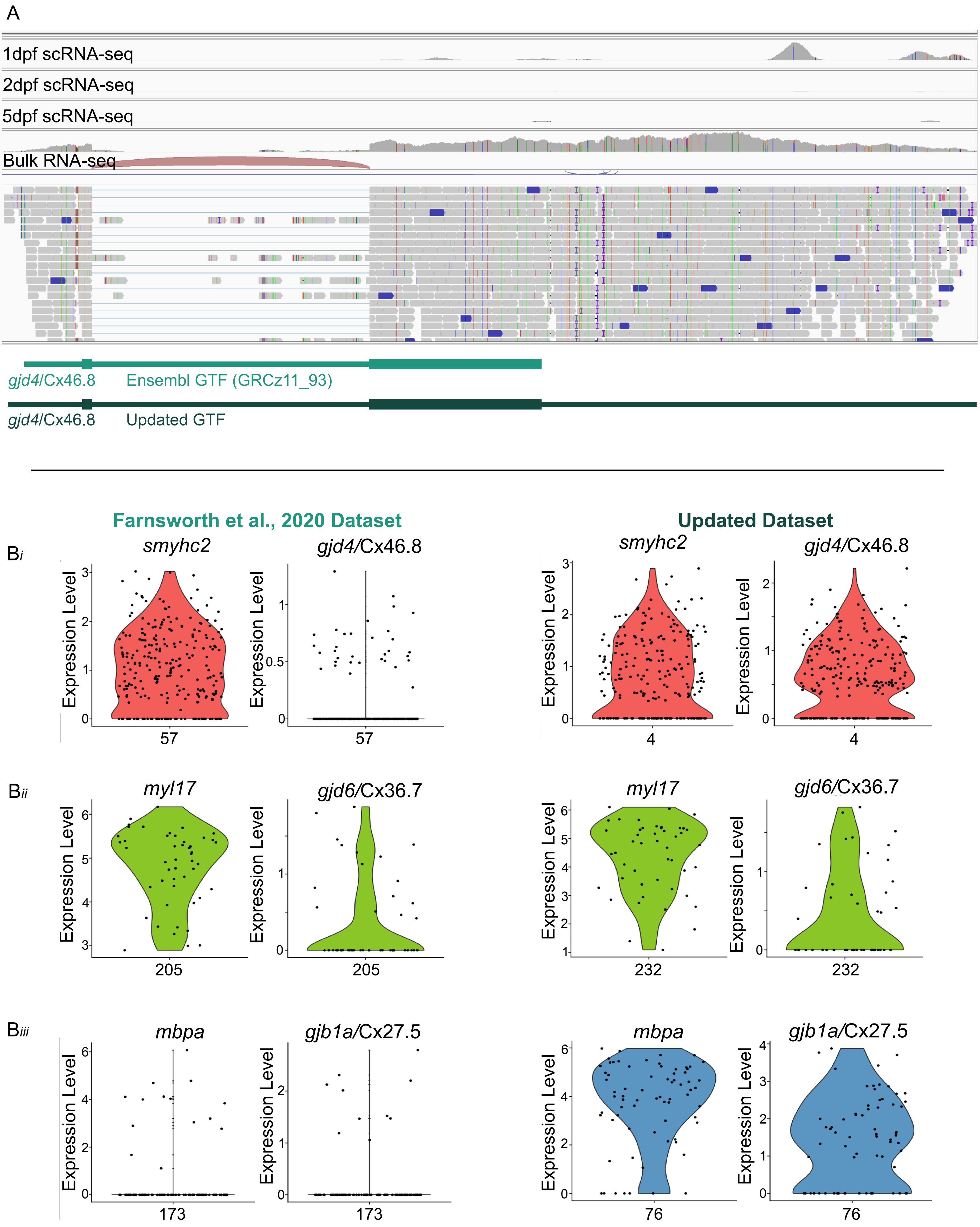

Supplemental Figure 4

A

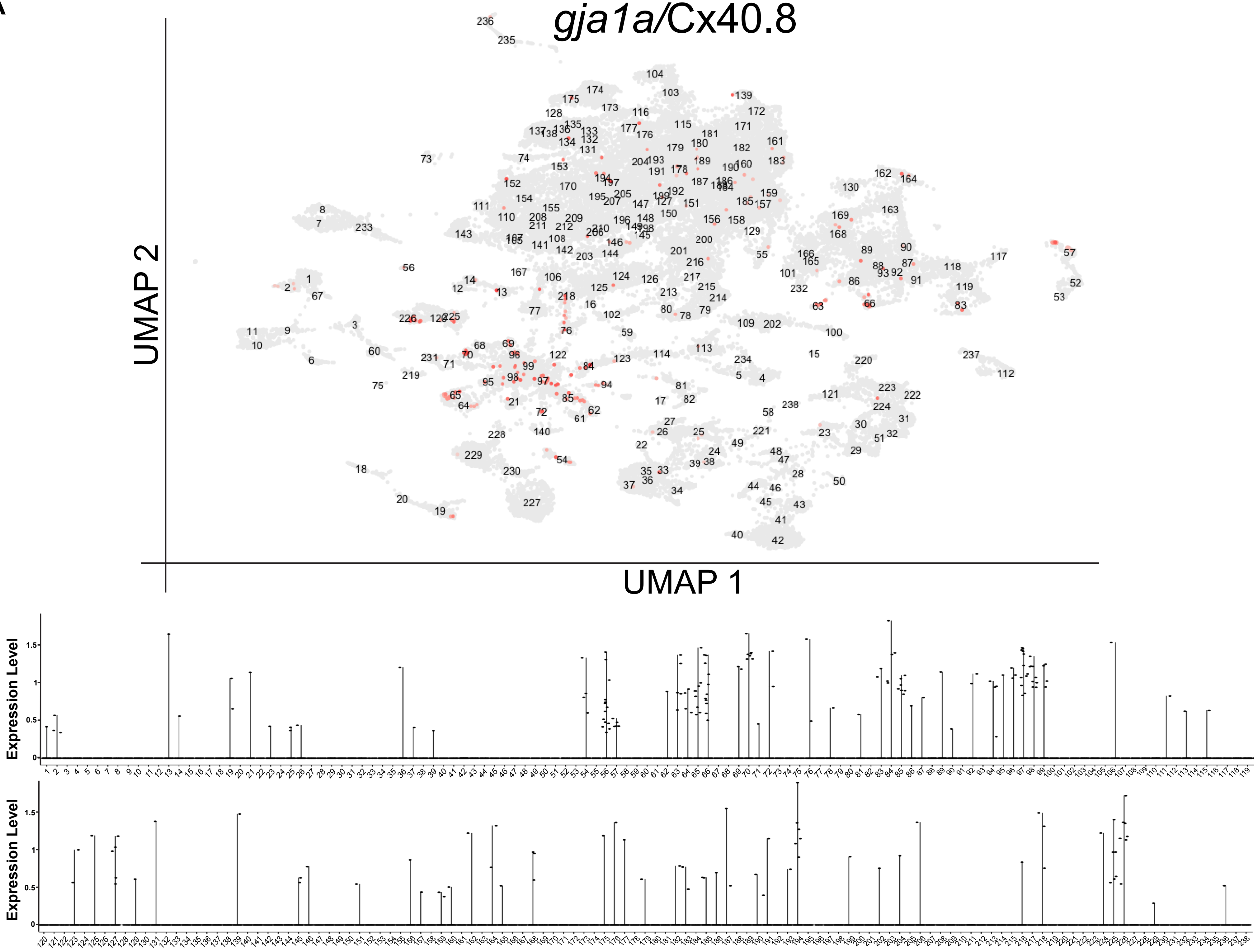

B

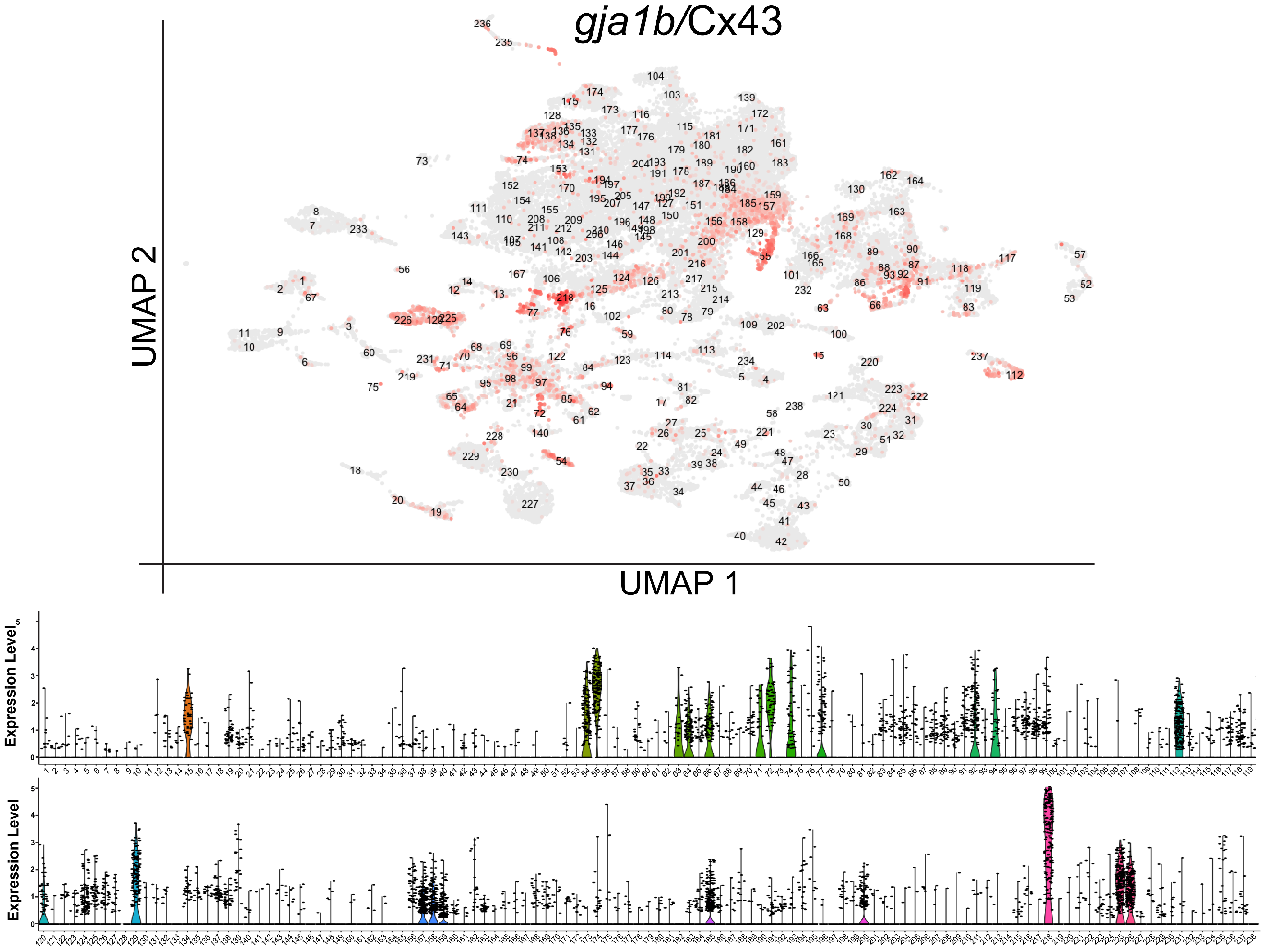

C

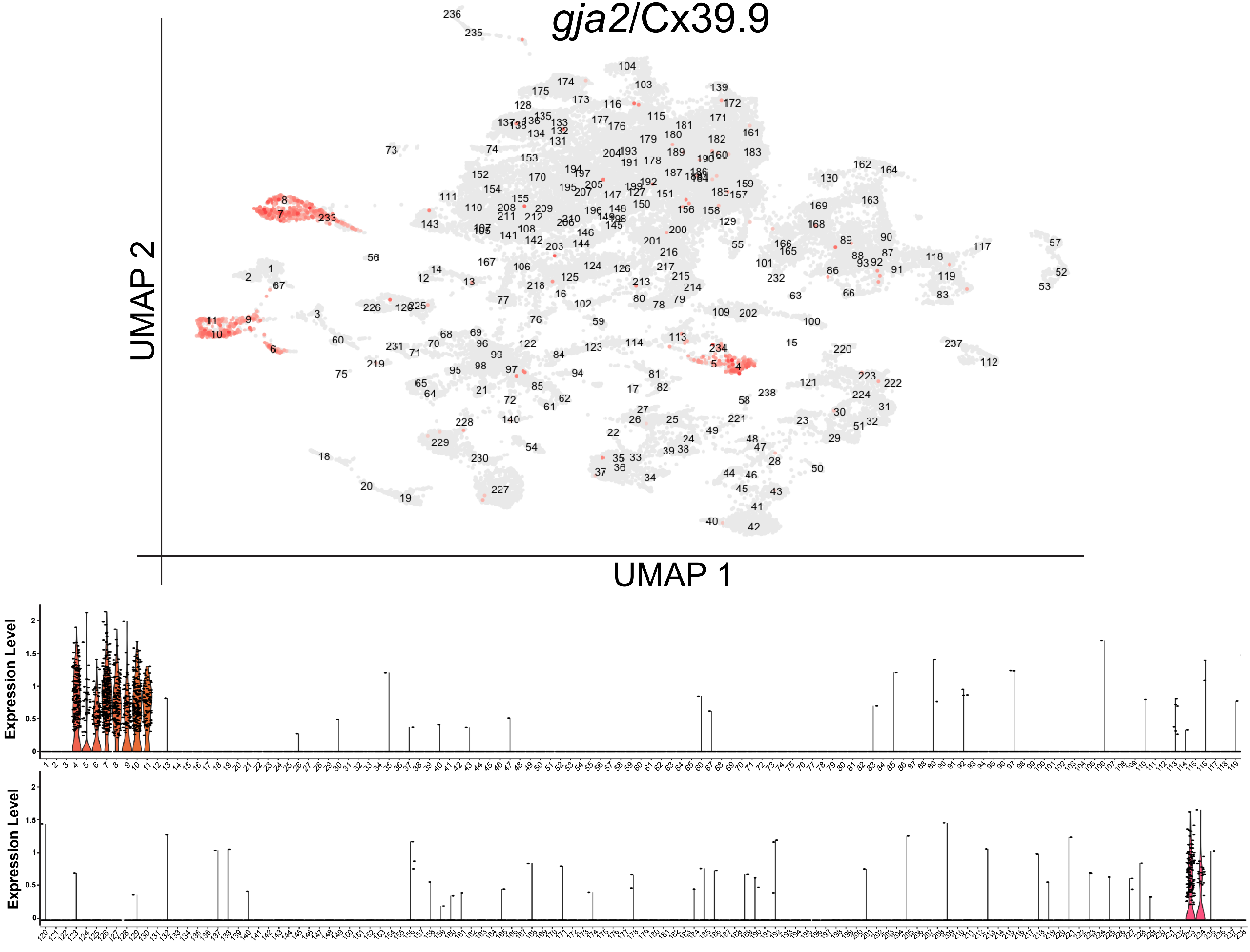

D

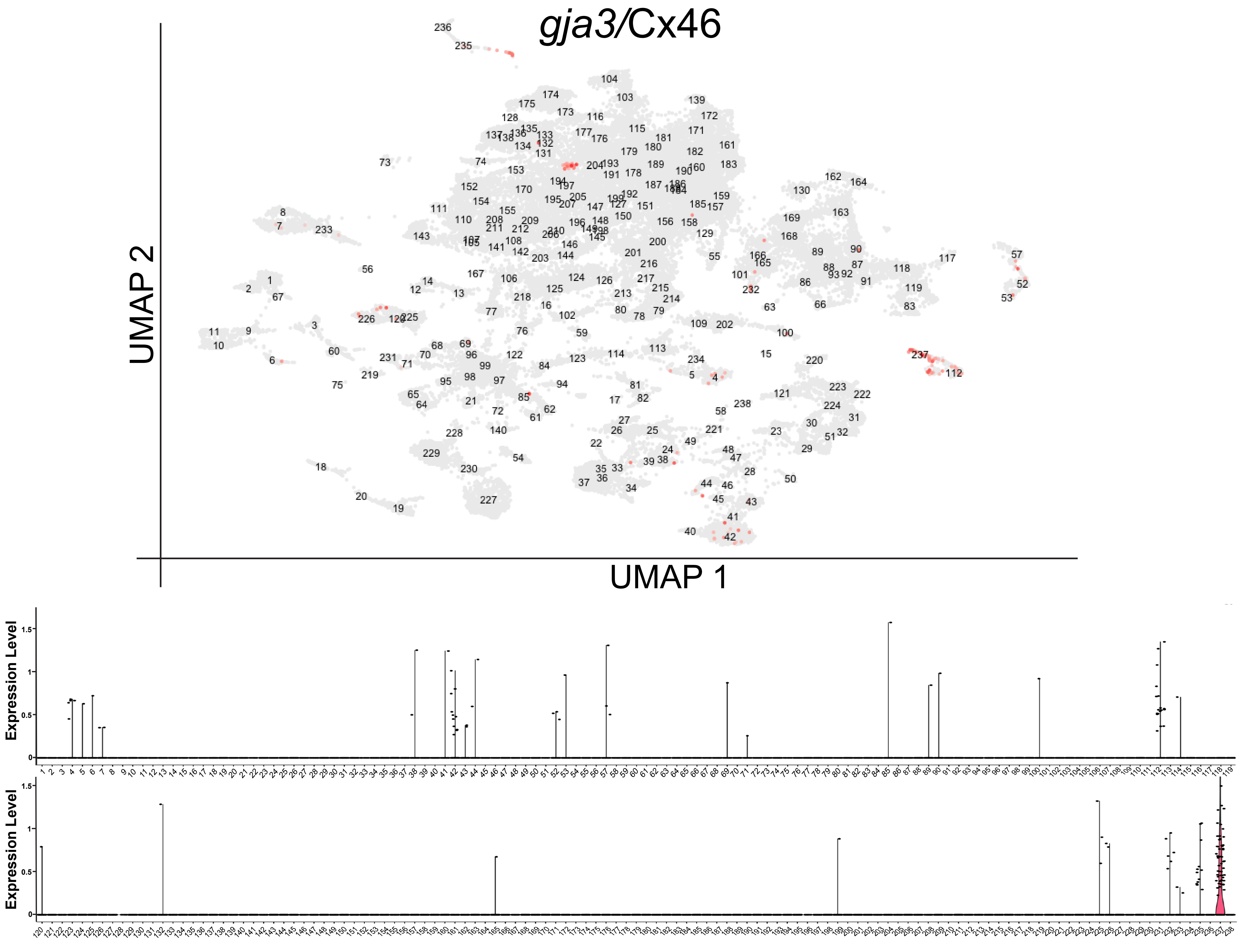

E

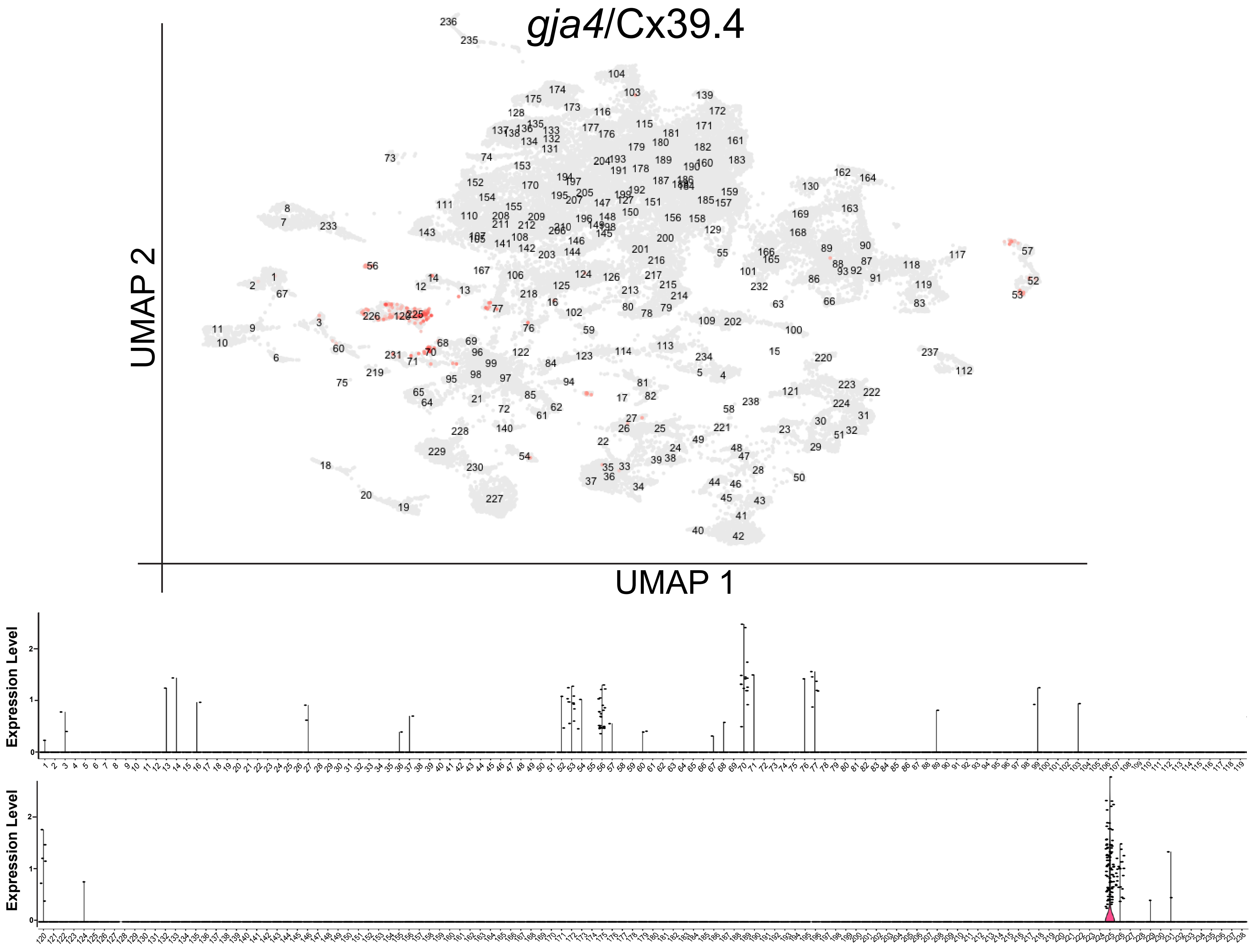

F

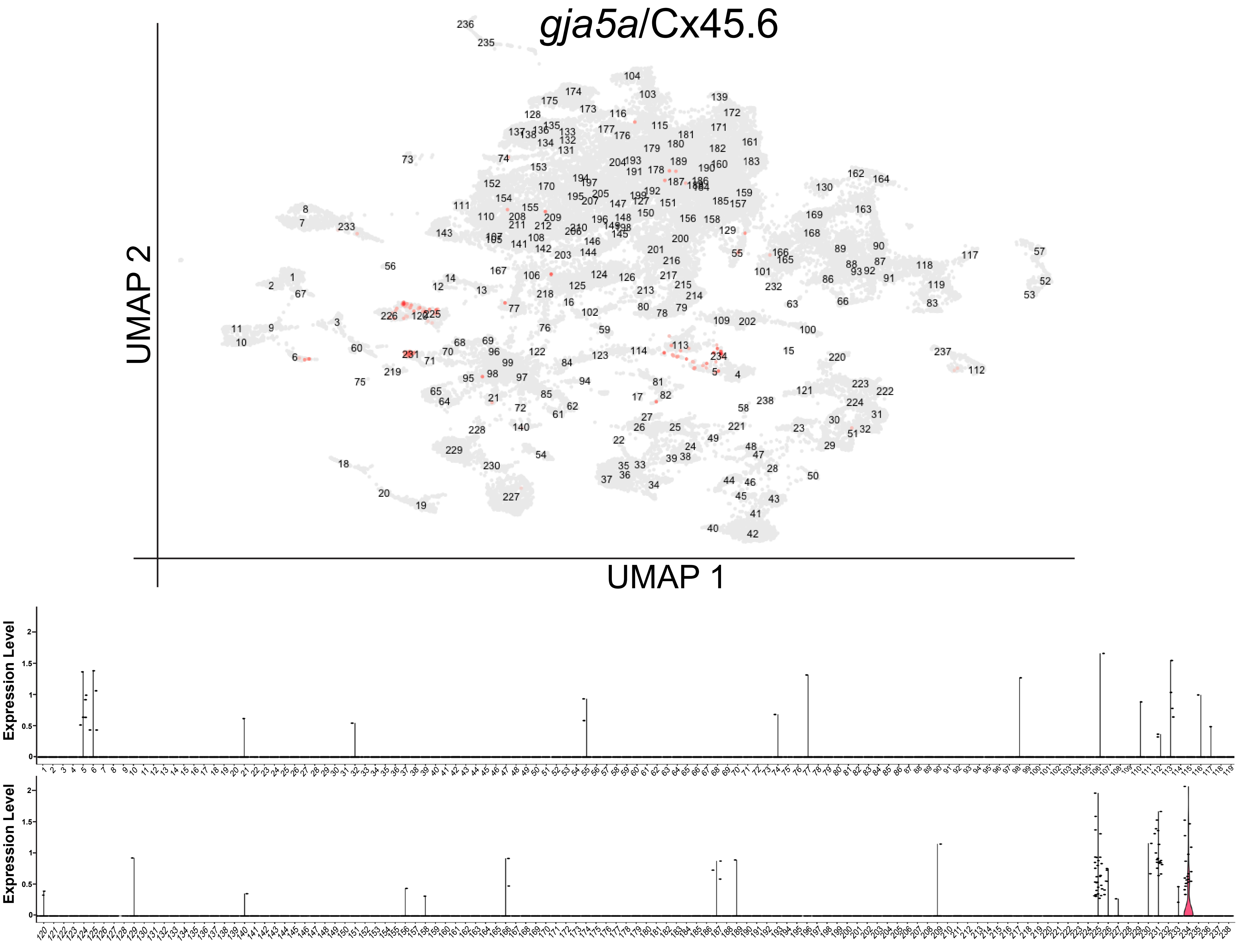

gja5a/Cx45.6
UMAP 2
UMAP 1
Expression Level
Expression Level

G

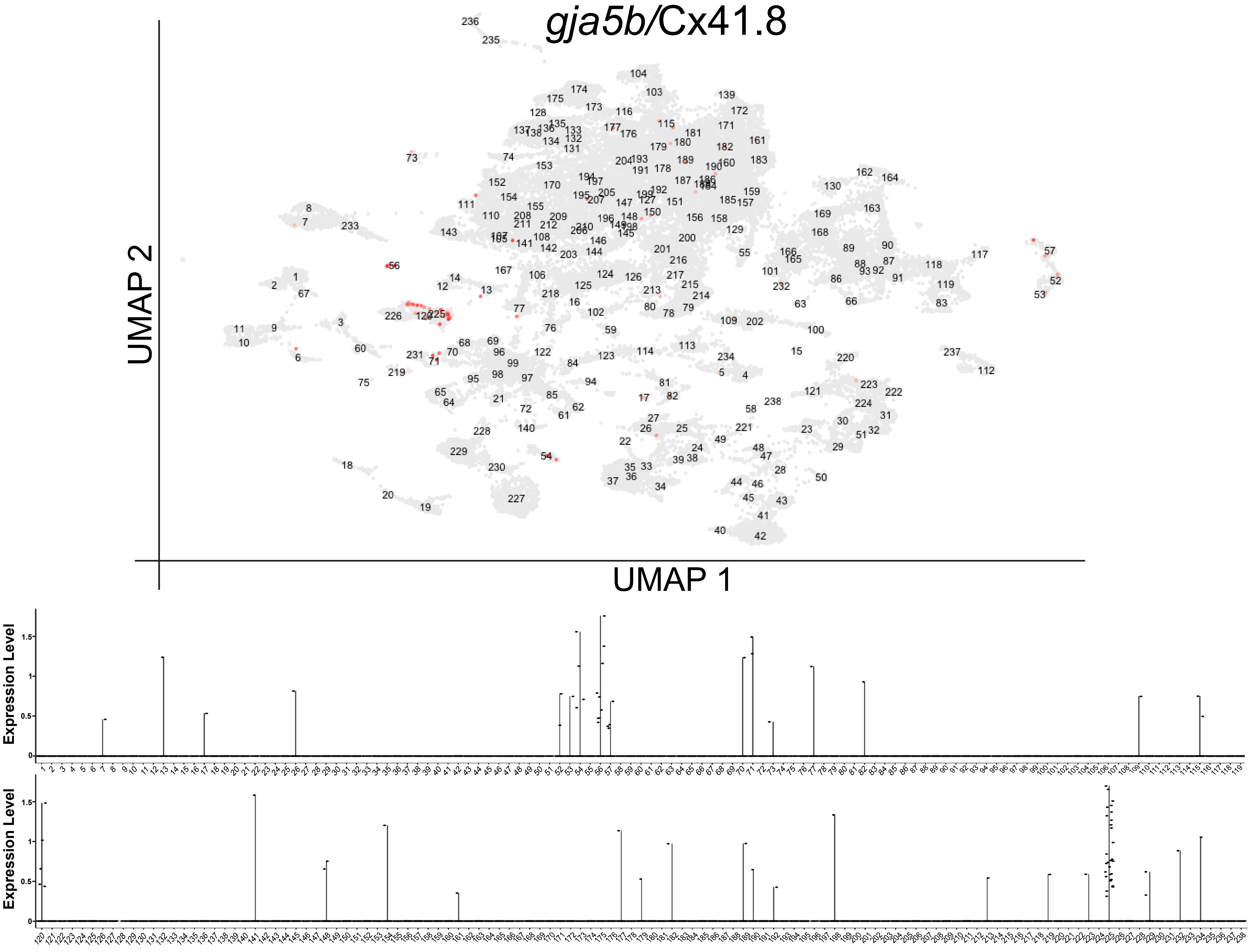

gja5b/Cx41.8
UMAP 2
UMAP 1
Expression Level
Expression Level

H

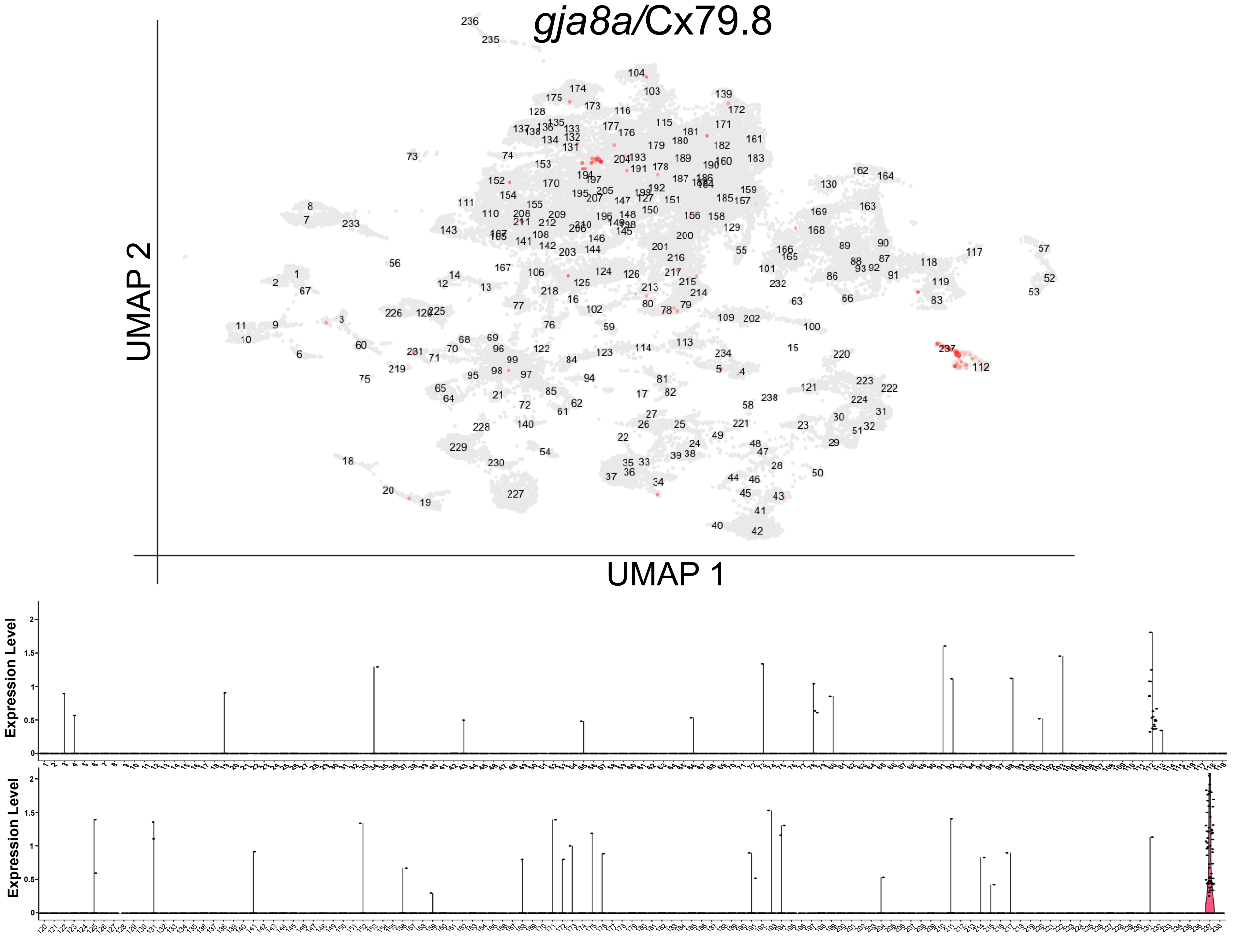

*gja8a/Cx79.8*
UMAP 2
UMAP 1
Expression Level
Expression Level

I

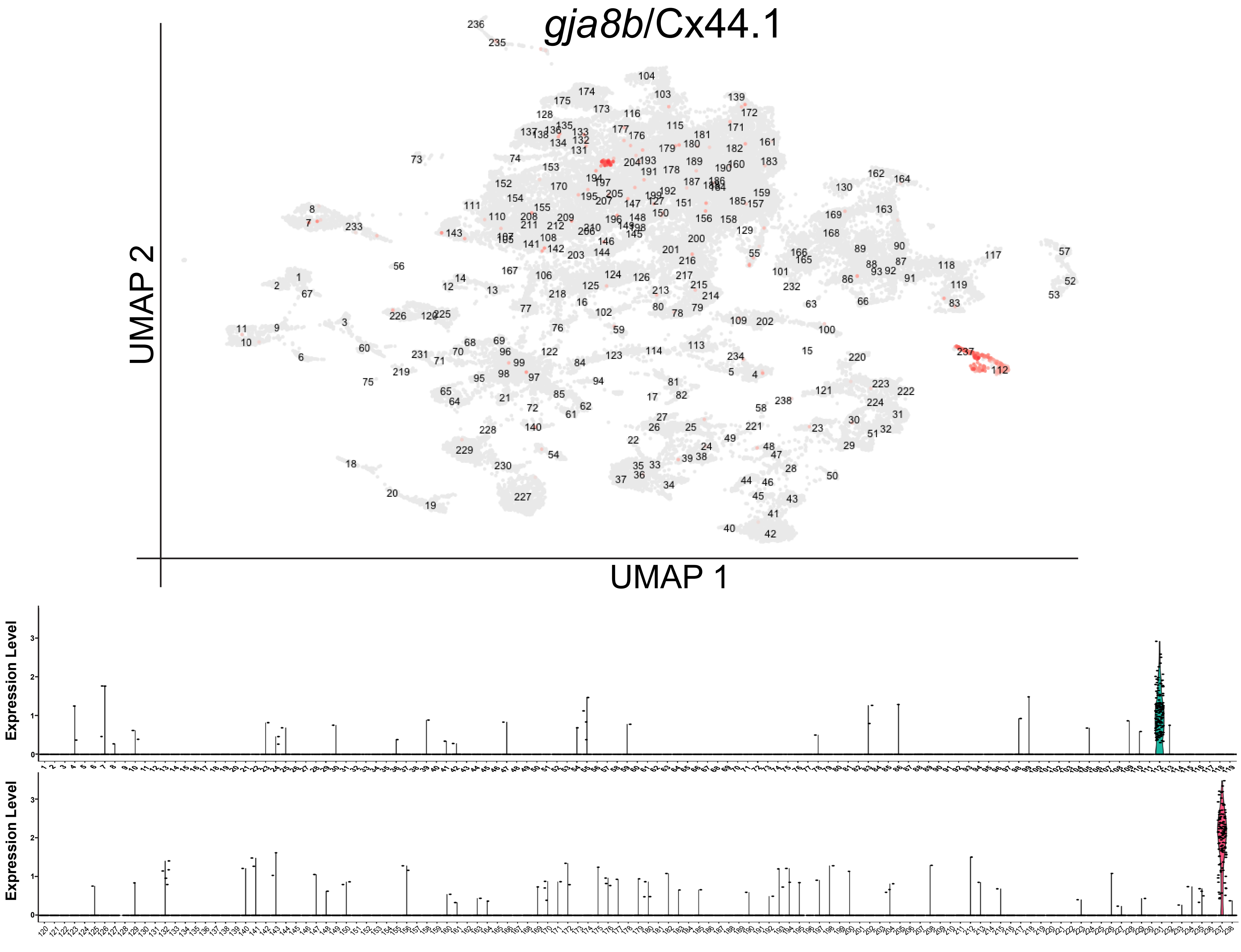

gja8b/Cx44.1
UMAP 2
UMAP 1
Expression Level
Expression Level

J

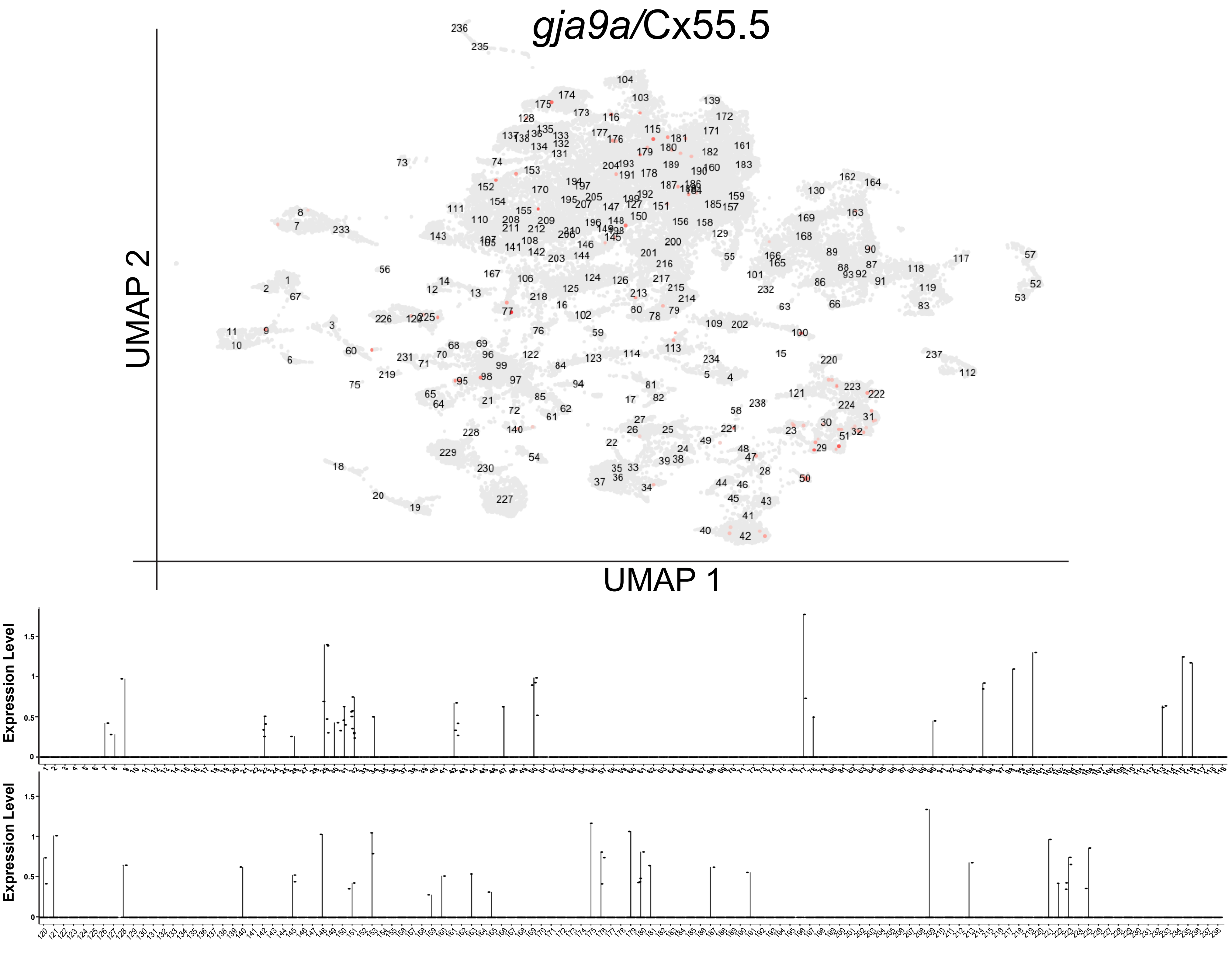

gja9a/Cx55.5
UMAP 2
UMAP 1
Expression Level
Expression Level

K

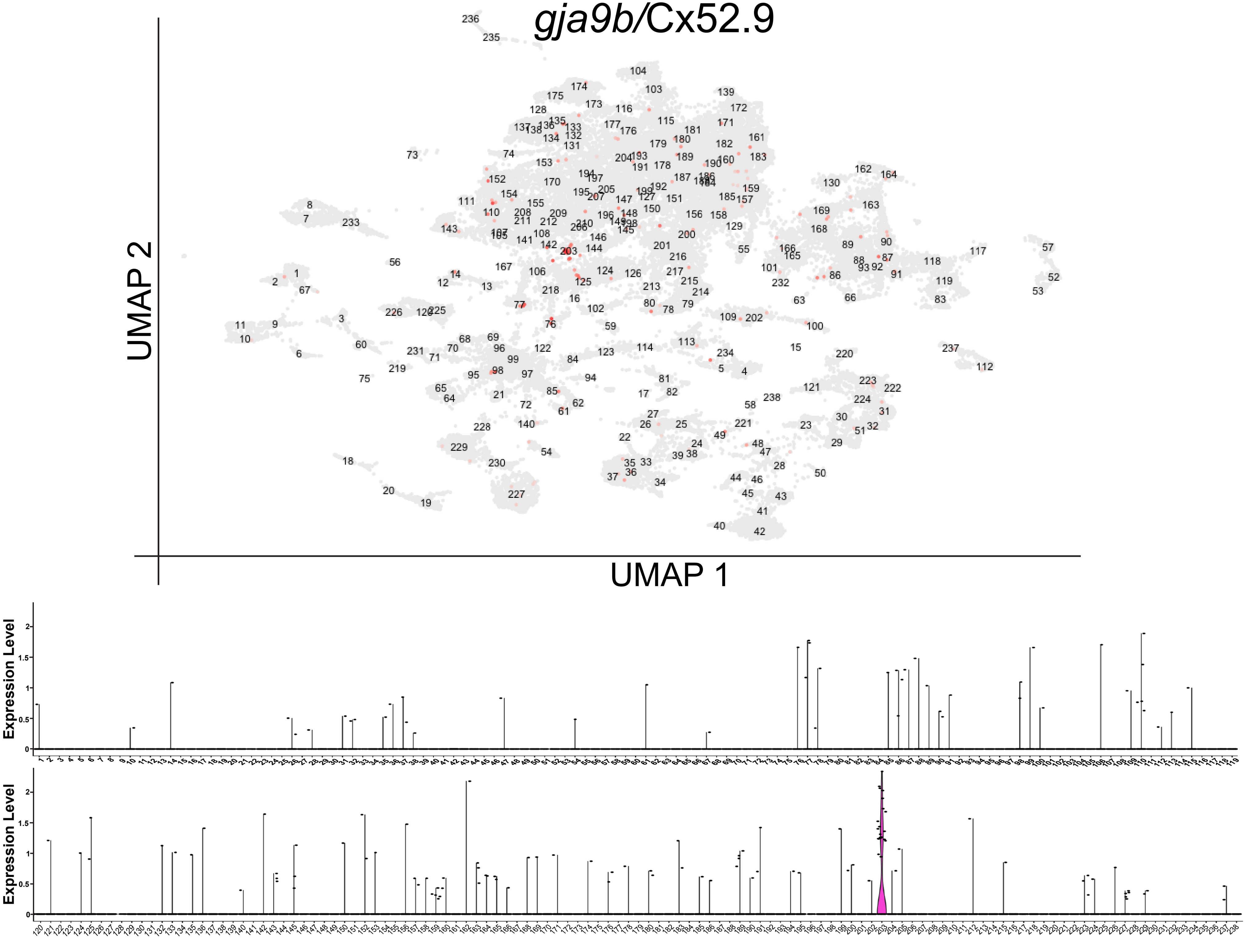
gja9b/Cx52.9
UMAP 2
UMAP 1
Expression Level
Expression Level

L

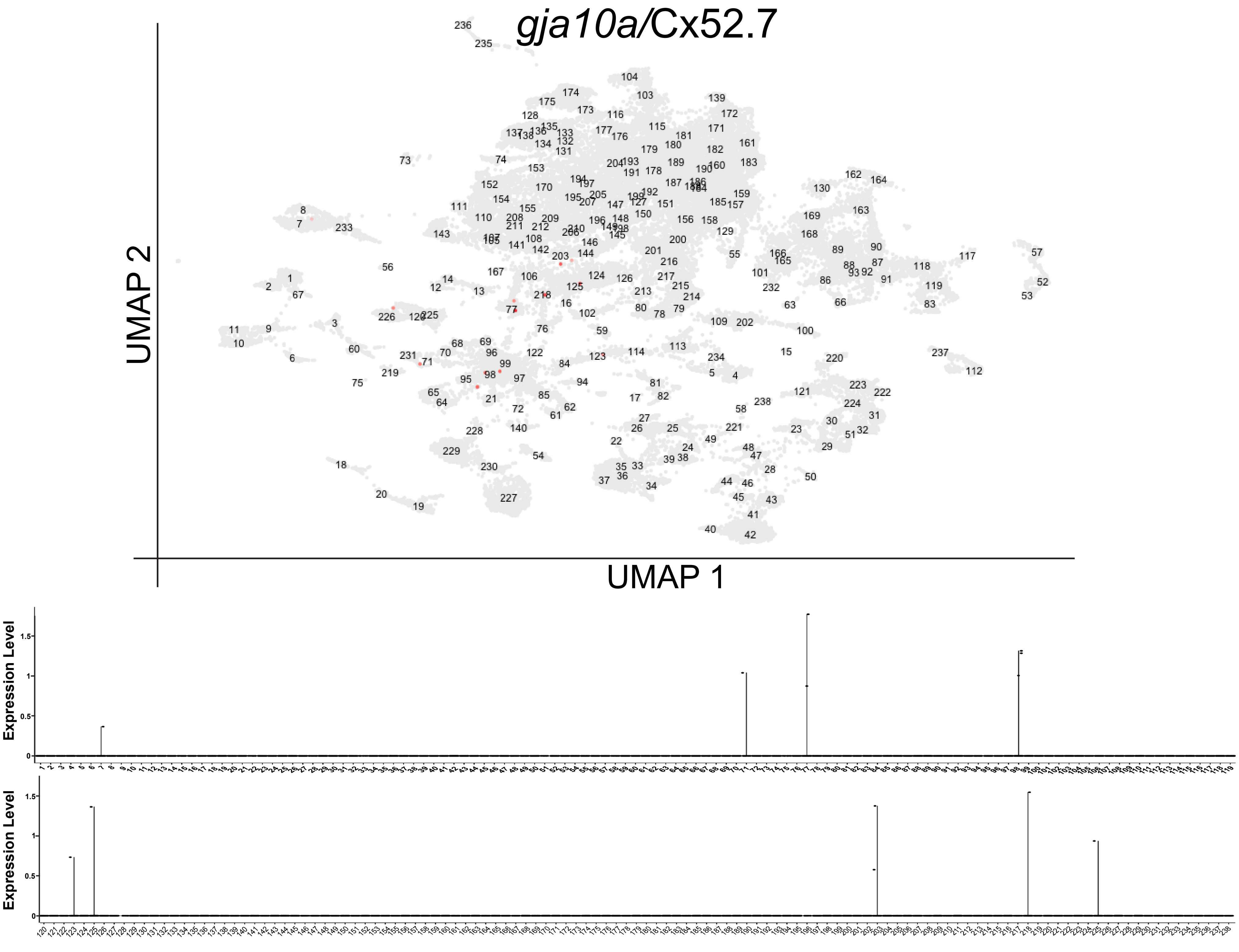

gja10a/Cx52.7
UMAP 2
UMAP 1
Expression Level
Expression Level

M

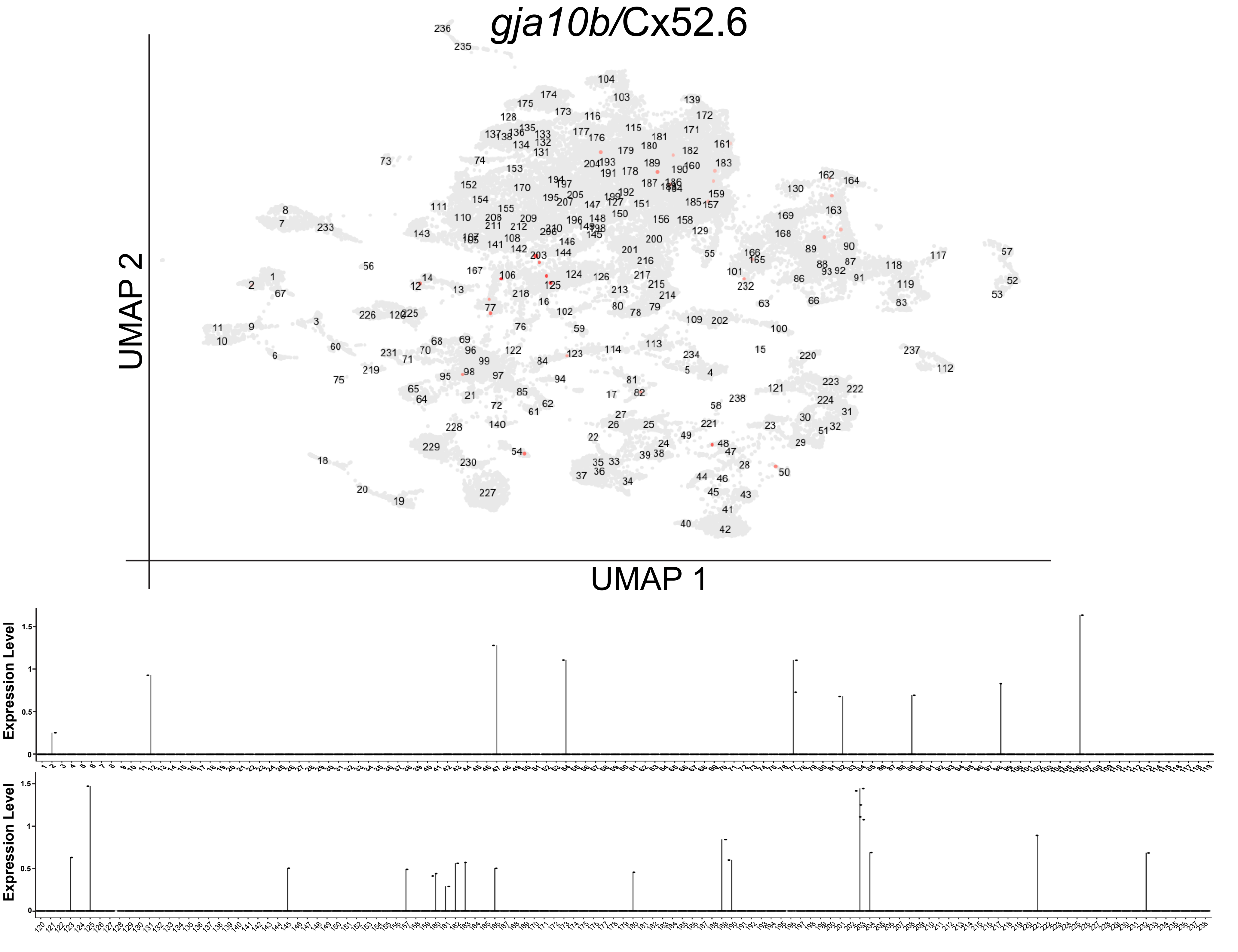
gja10b/Cx52.6
UMAP 2
UMAP 1
Expression Level
Expression Level

N

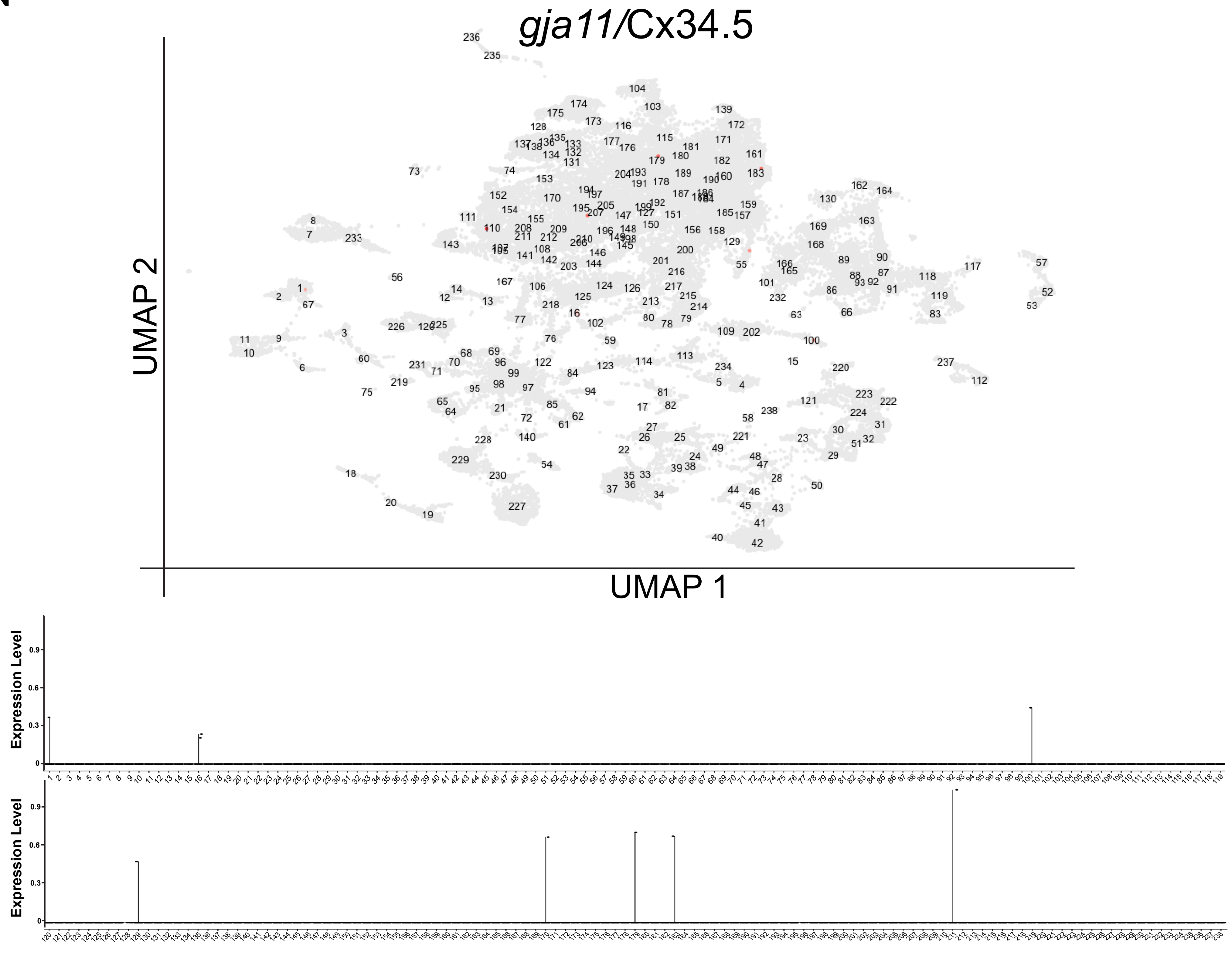

gja11/Cx34.5
UMAP 2
UMAP 1
Expression Level
Expression Level

O

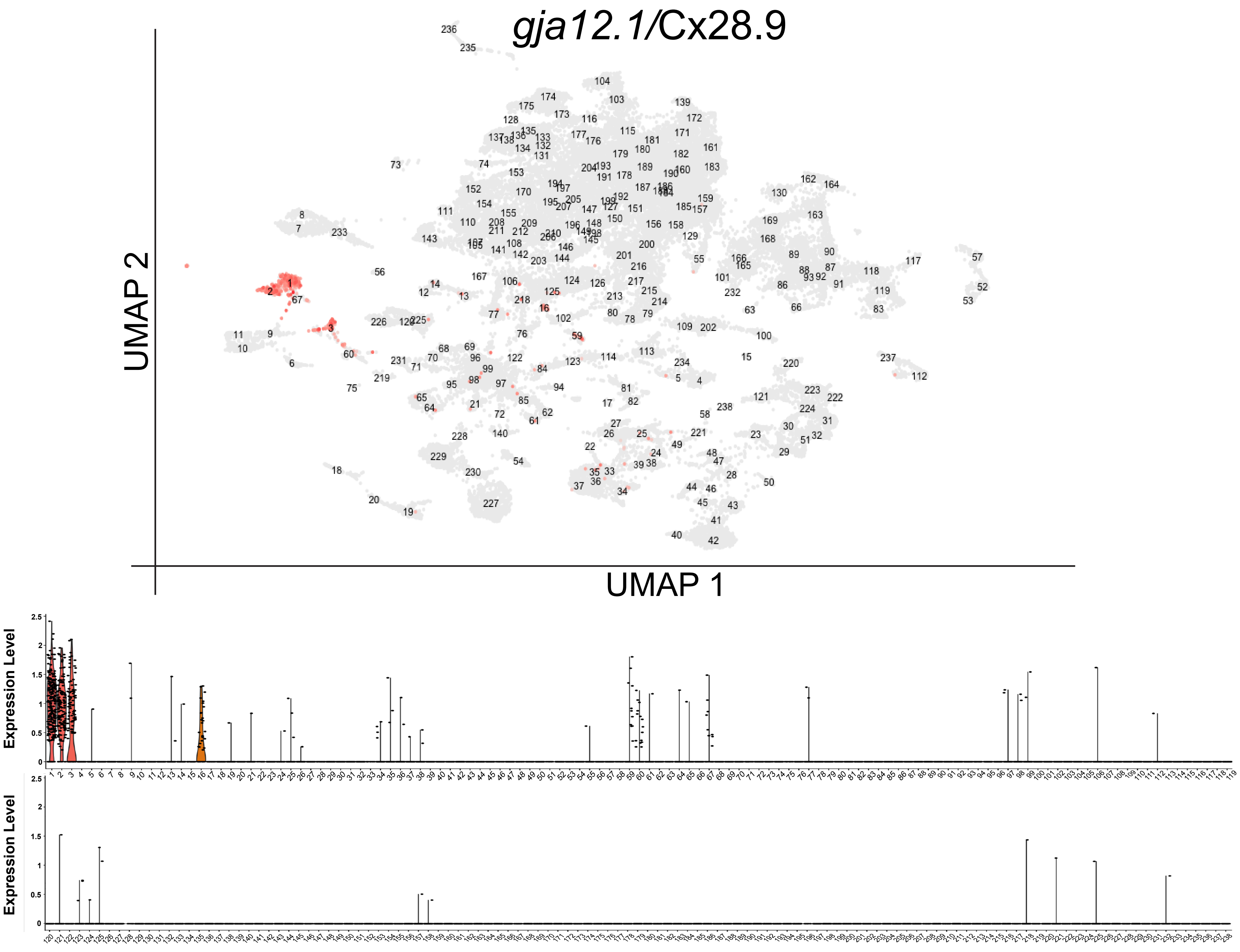

gja12.1/Cx28.9
UMAP 2
UMAP 1
Expression Level
Expression Level

P

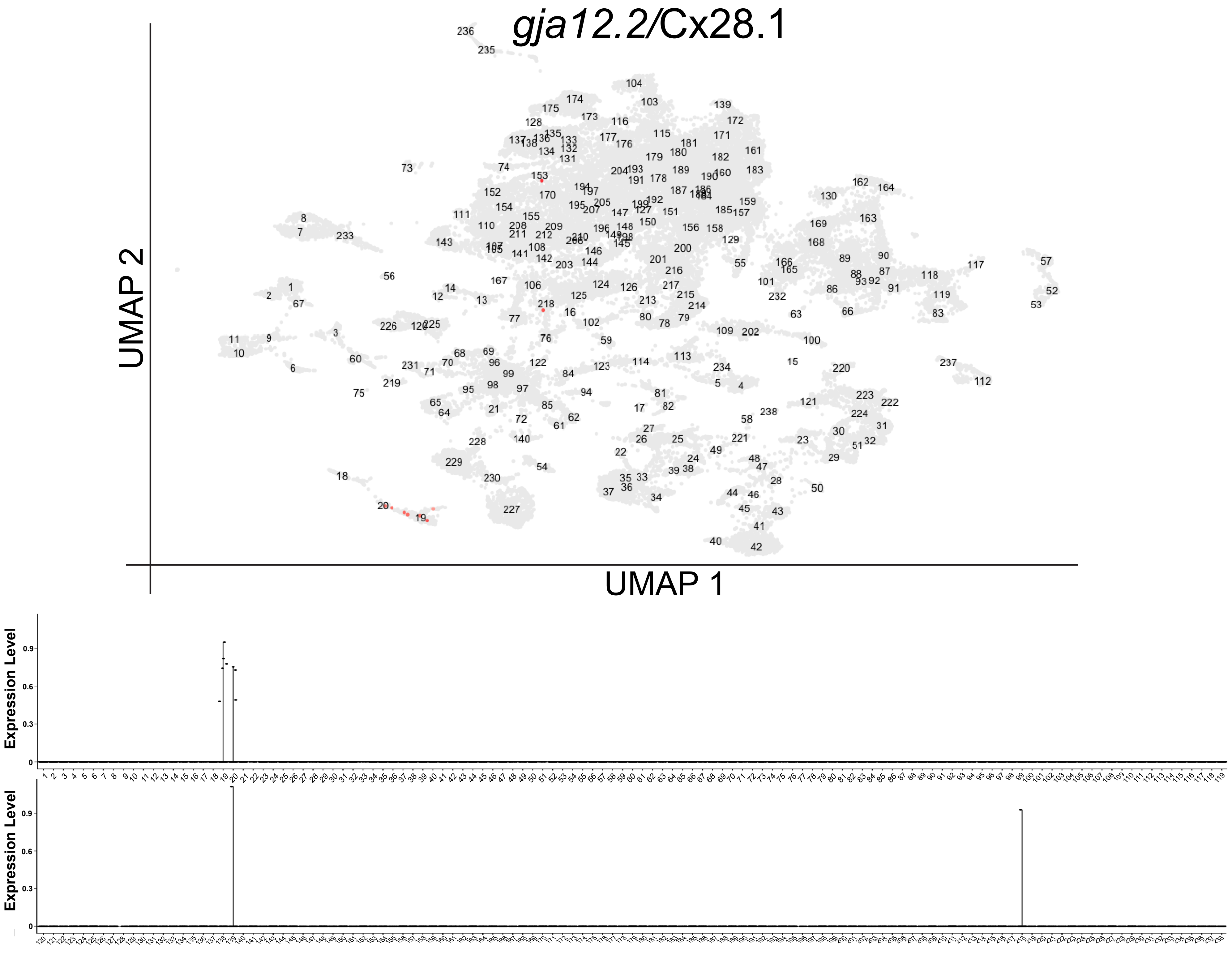
gja12.2/Cx28.1
UMAP 2
UMAP 1
Expression Level
Expression Level

Q

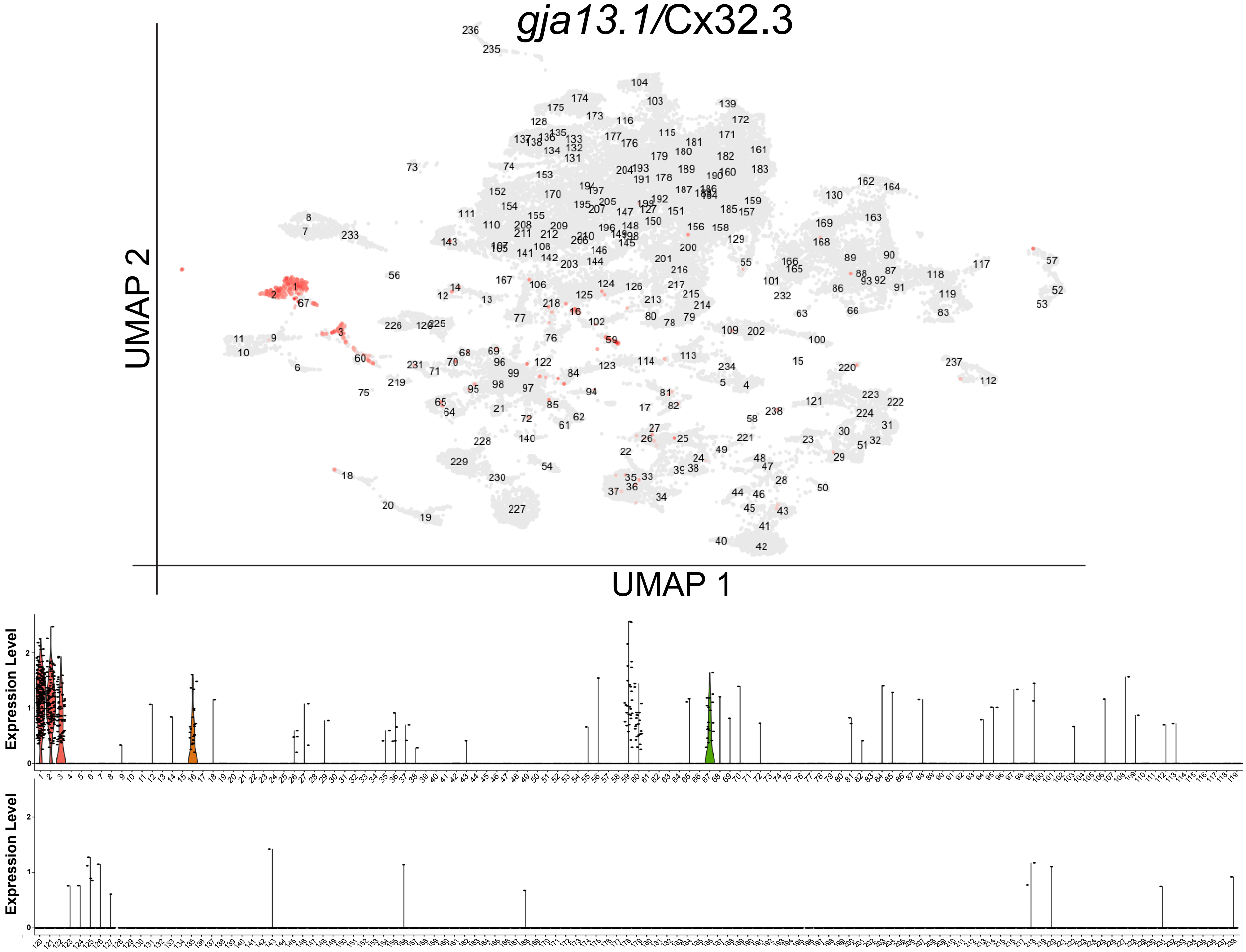
*gja13.1/Cx32.3*
UMAP 2
UMAP 1
Expression Level
Expression Level

R

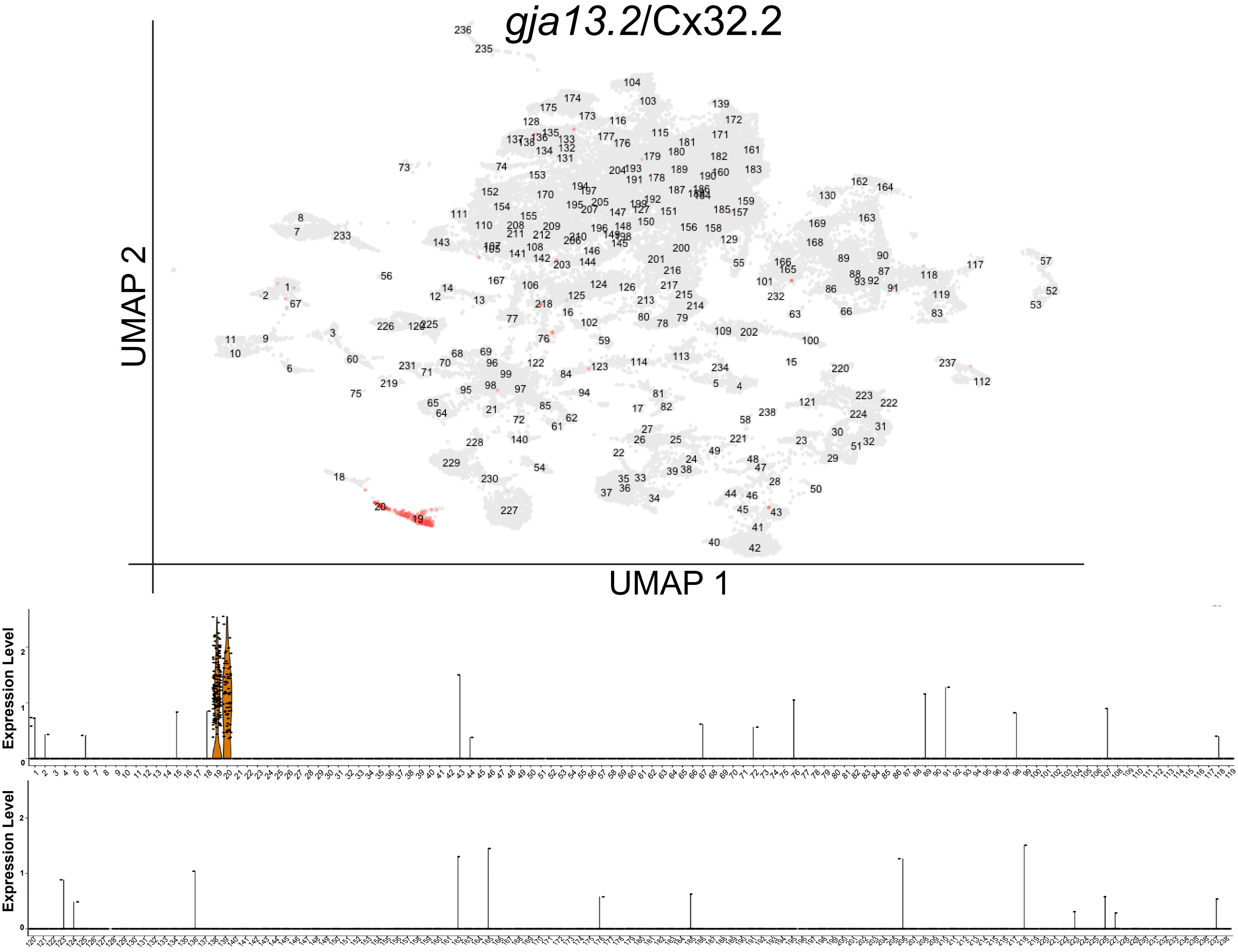
gja13.2/Cx32.2
UMAP 2
UMAP 1
Expression Level
Expression Level

S

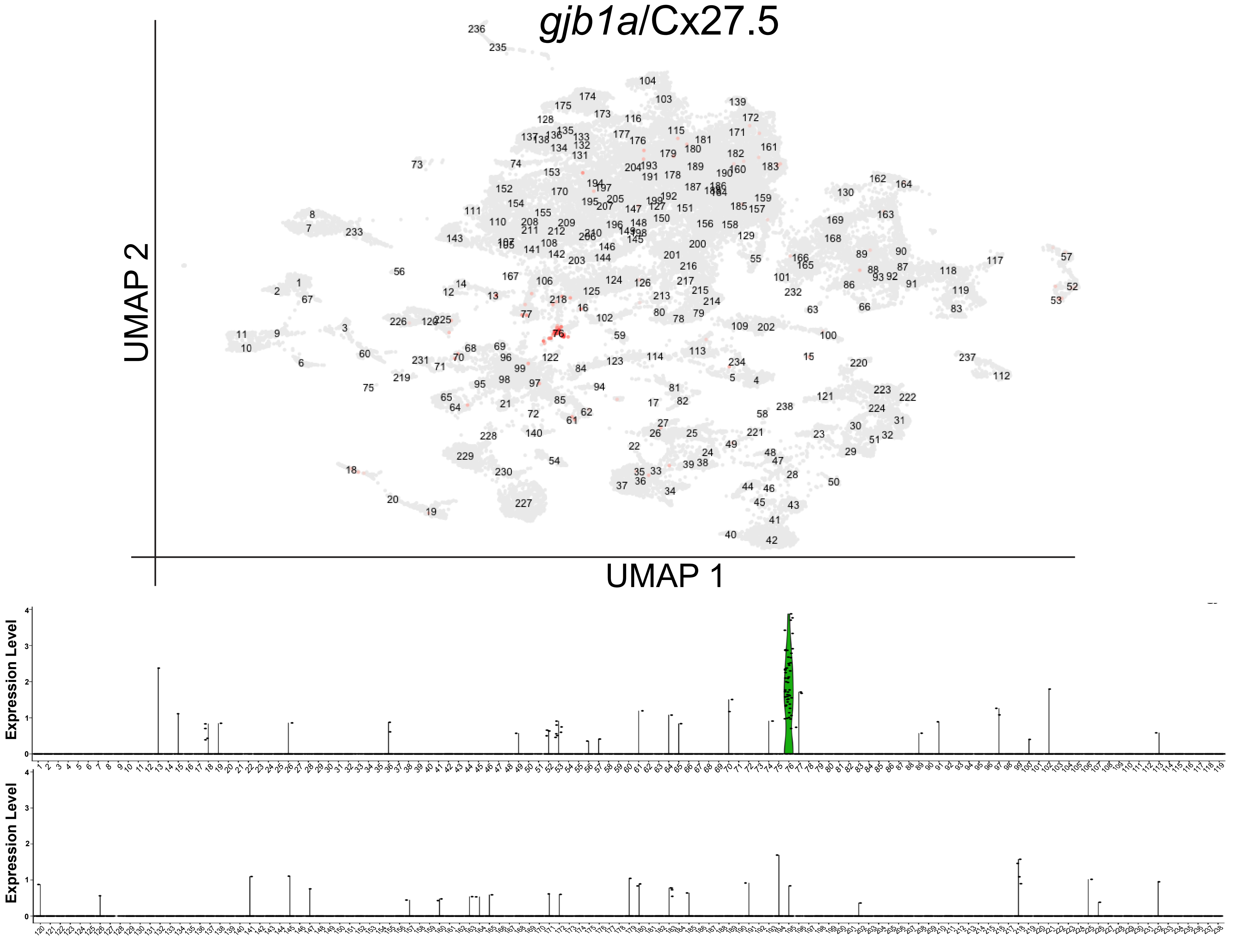

gjb1a/Cx27.5
UMAP 2
UMAP 1
Expression Level
Expression Level

T

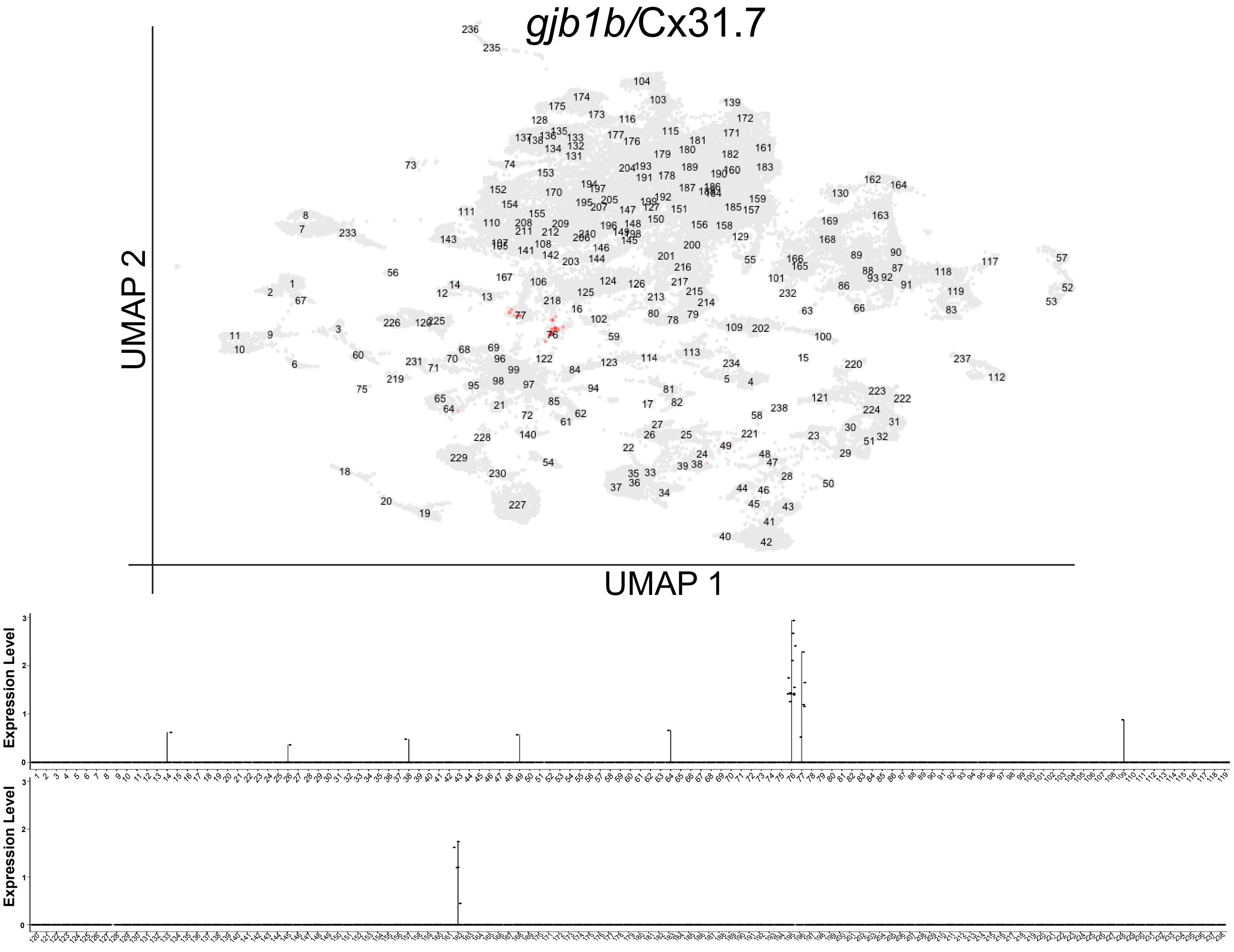

gjb1b/Cx31.7
UMAP 2
UMAP 1
Expression Level
Expression Level

U

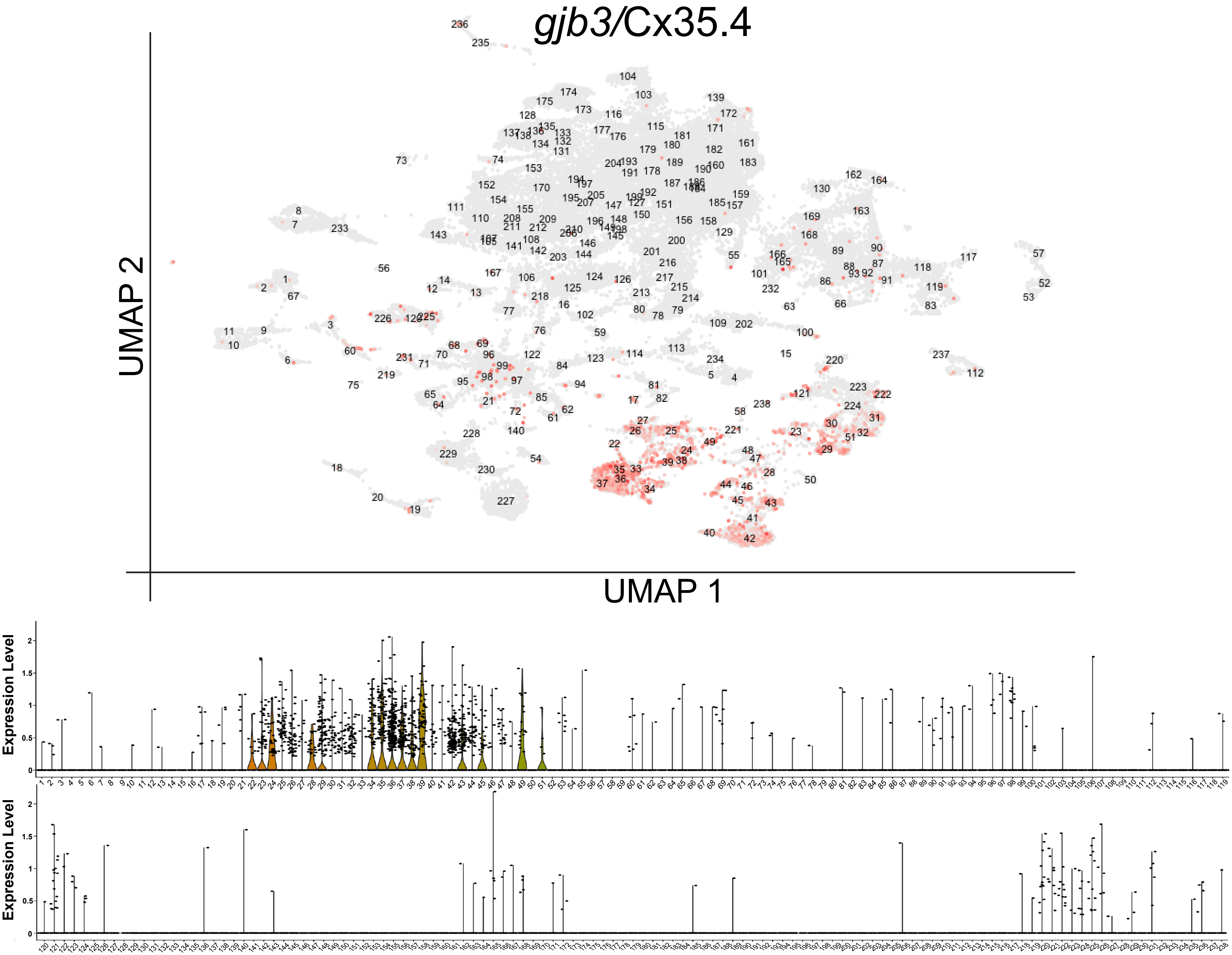

V

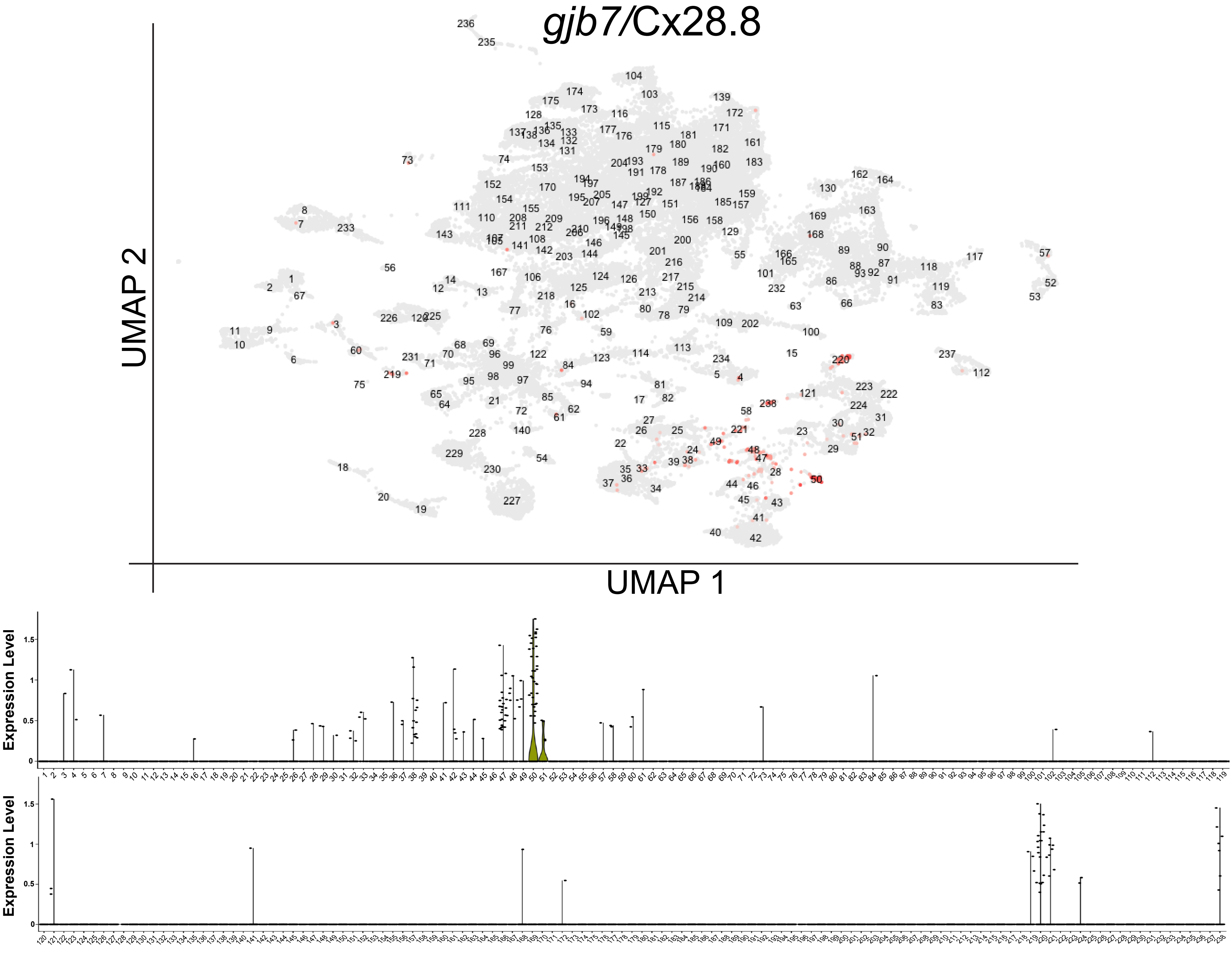

W

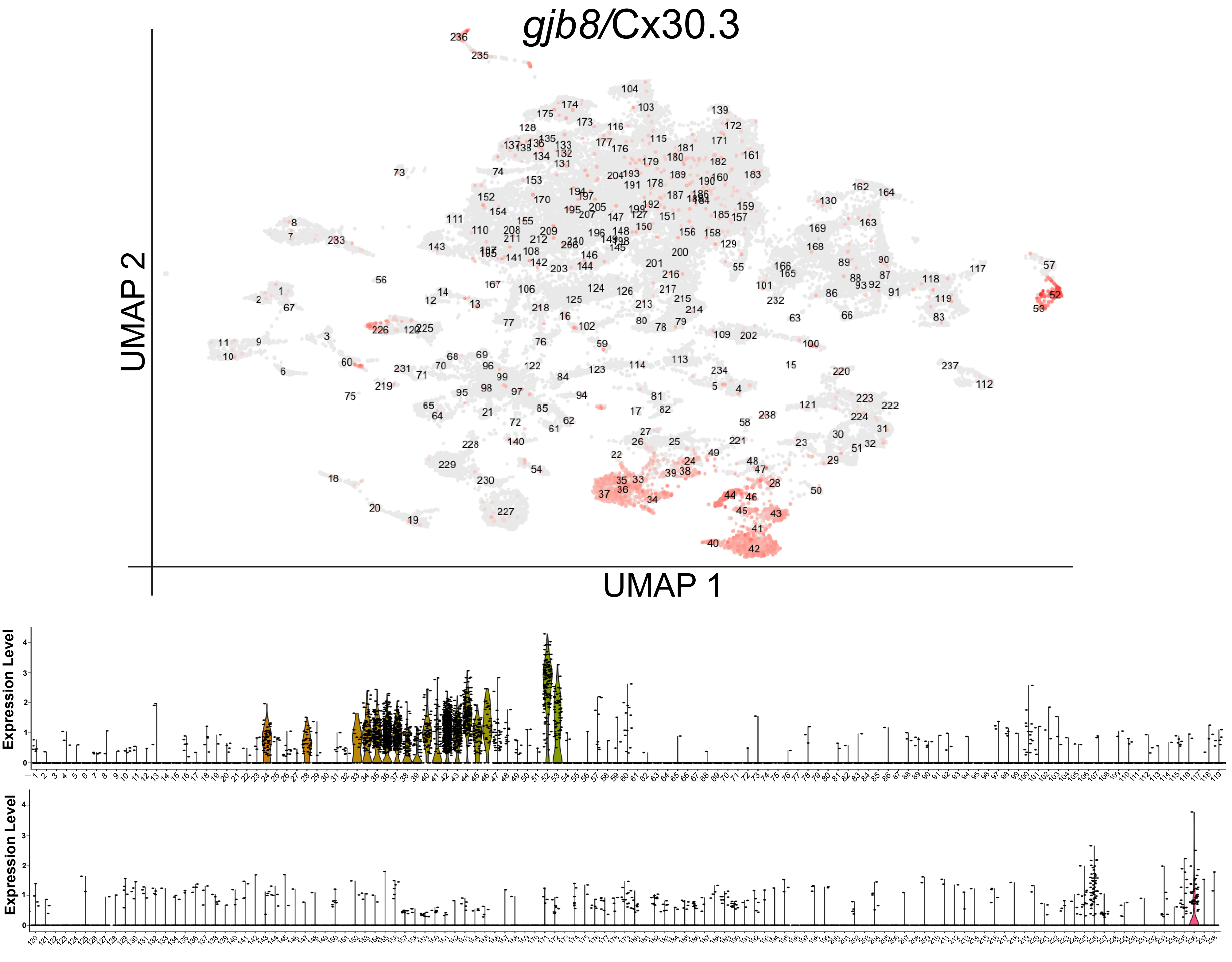

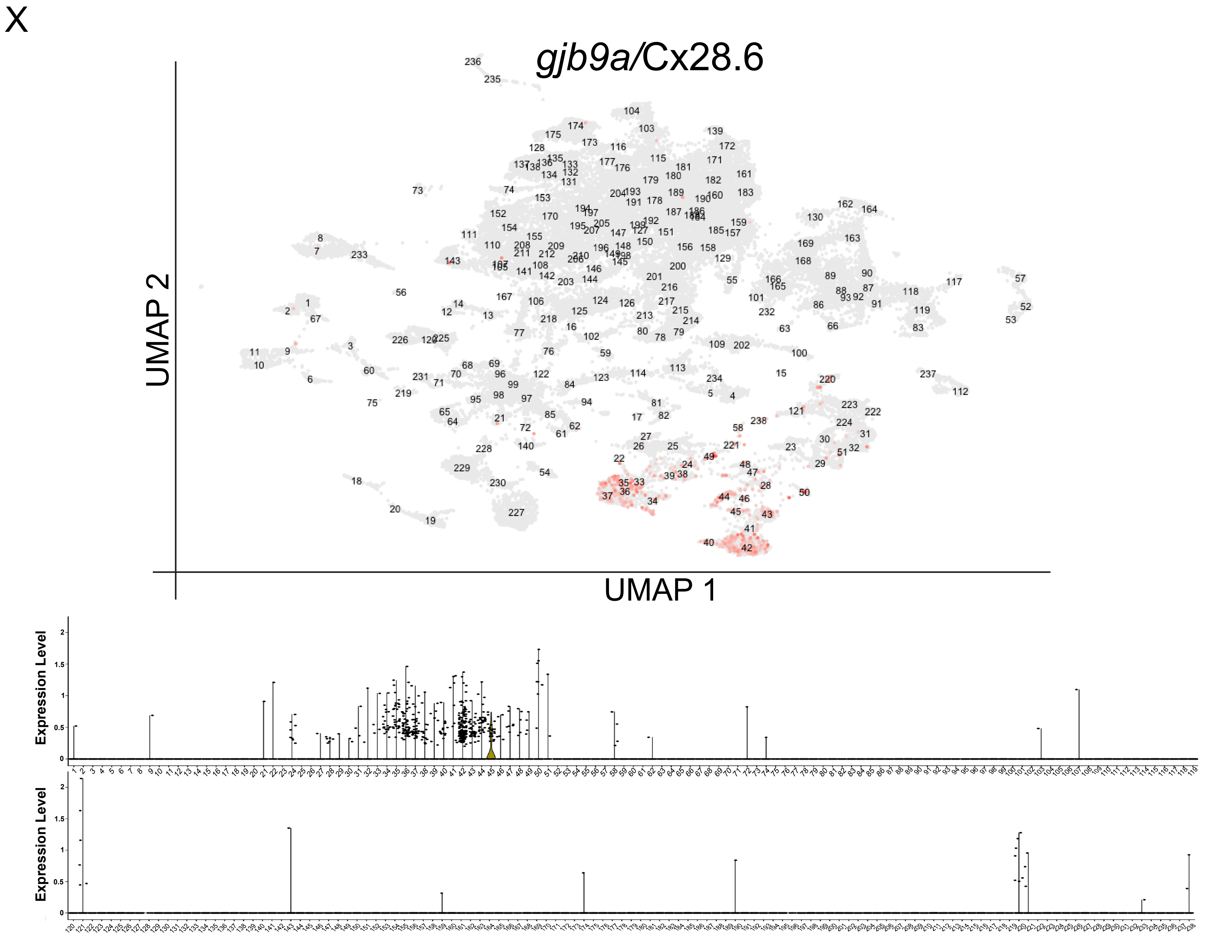
X
gjb9a/Cx28.6
UMAP 2
UMAP 1
Expression Level
Expression Level

Y

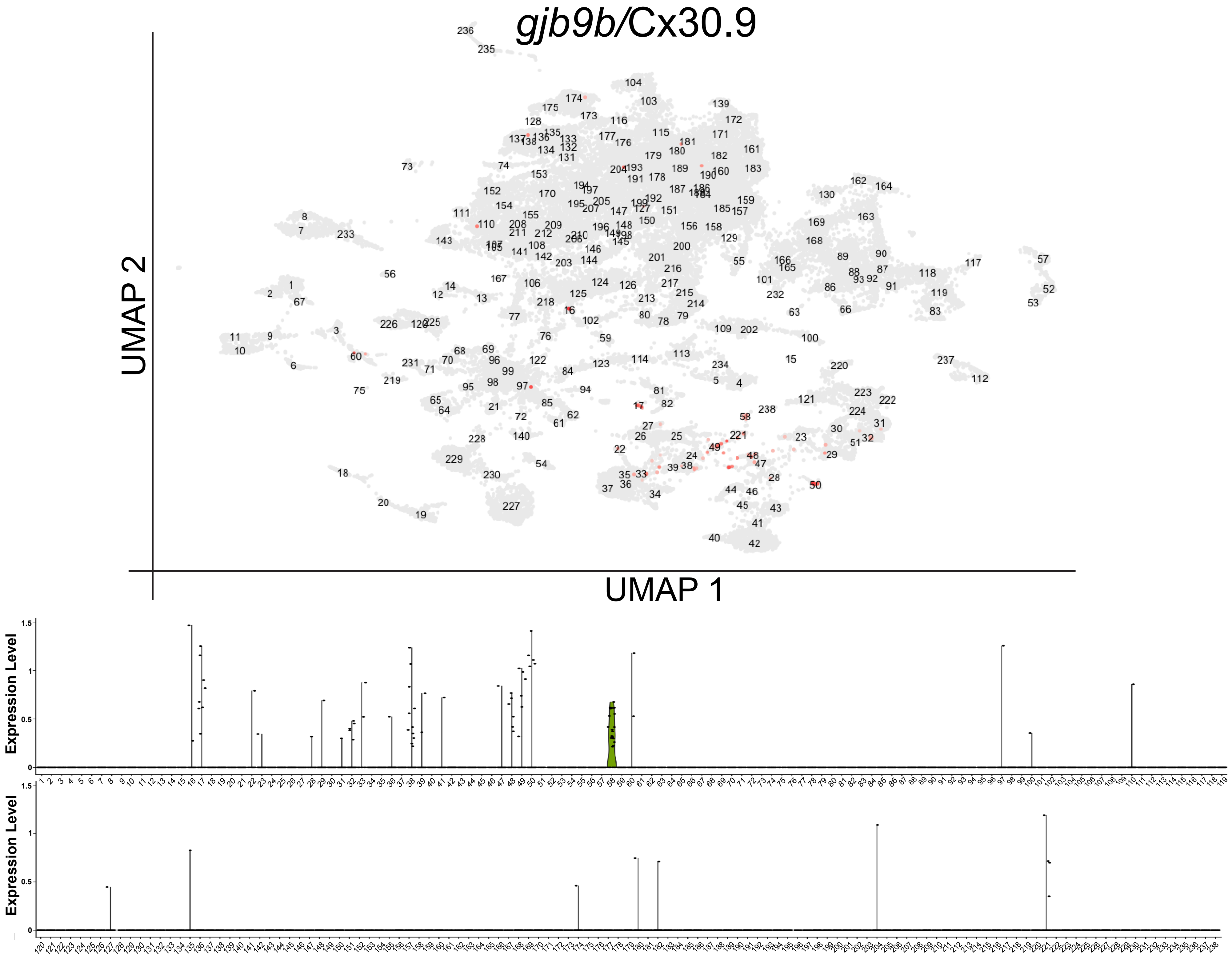

gjb9b/Cx30.9
UMAP 2
UMAP 1
Expression Level
Expression Level

Z

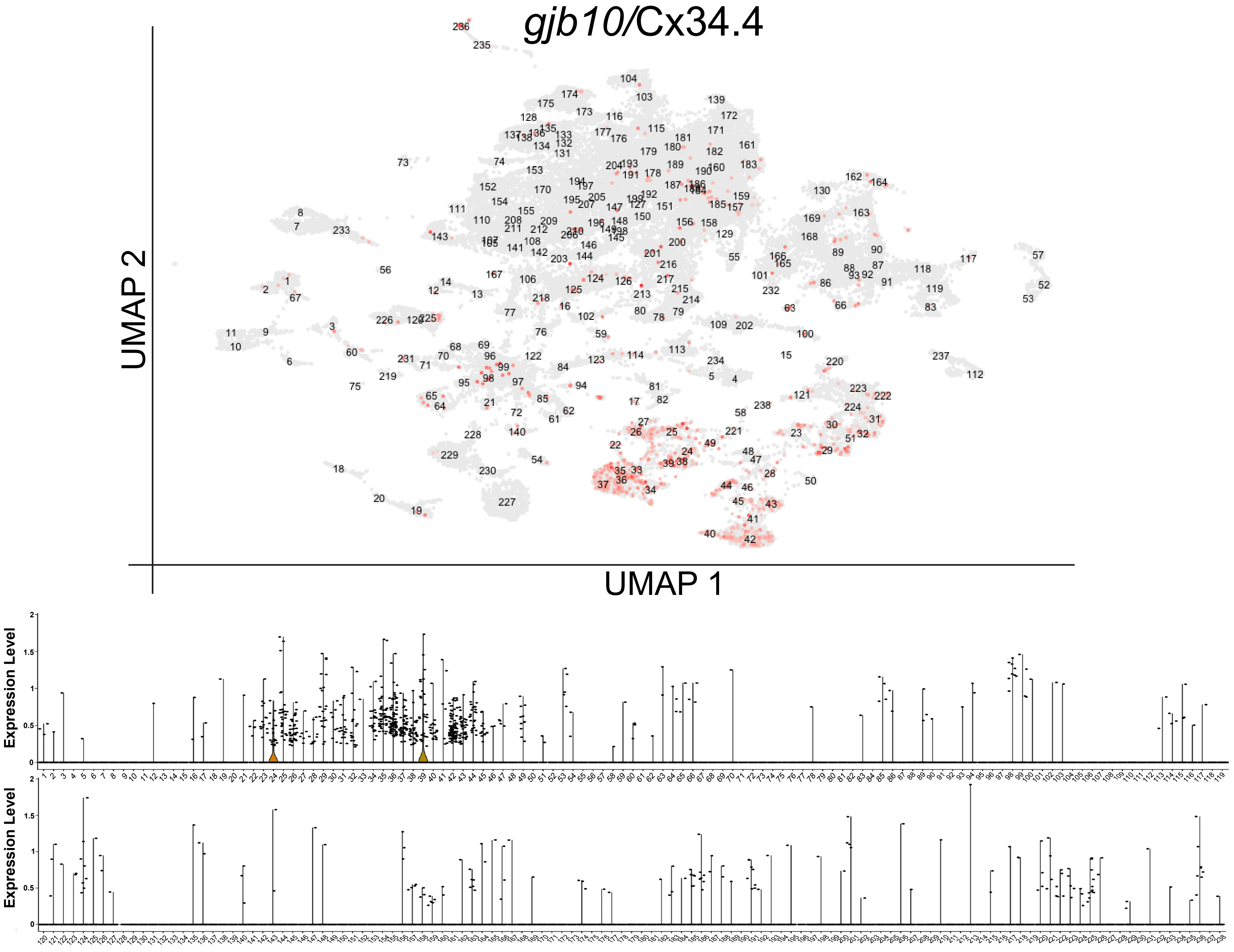

*gjb10*/Cx34.4
UMAP 2
UMAP 1
Expression Level
Expression Level

AA

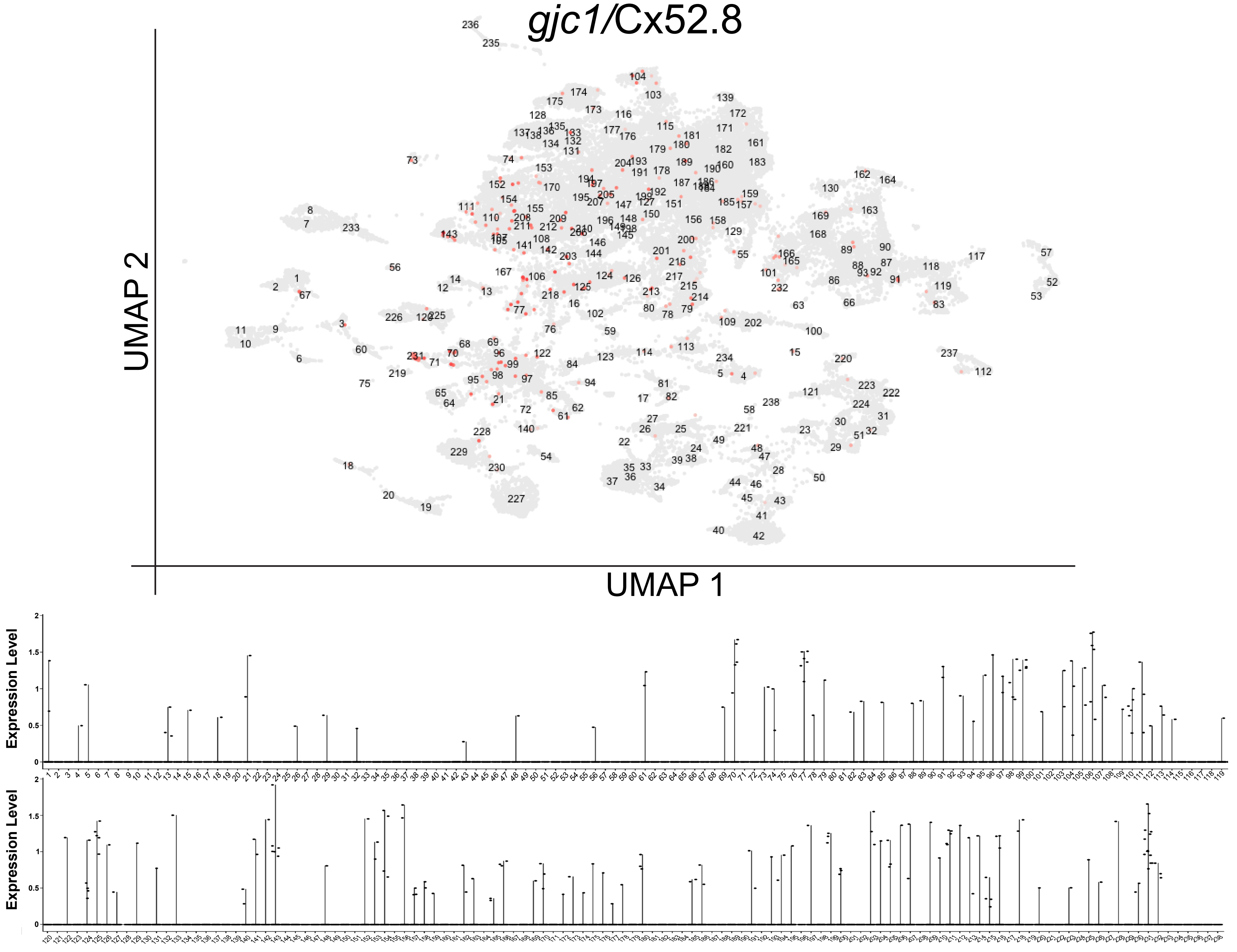

BB

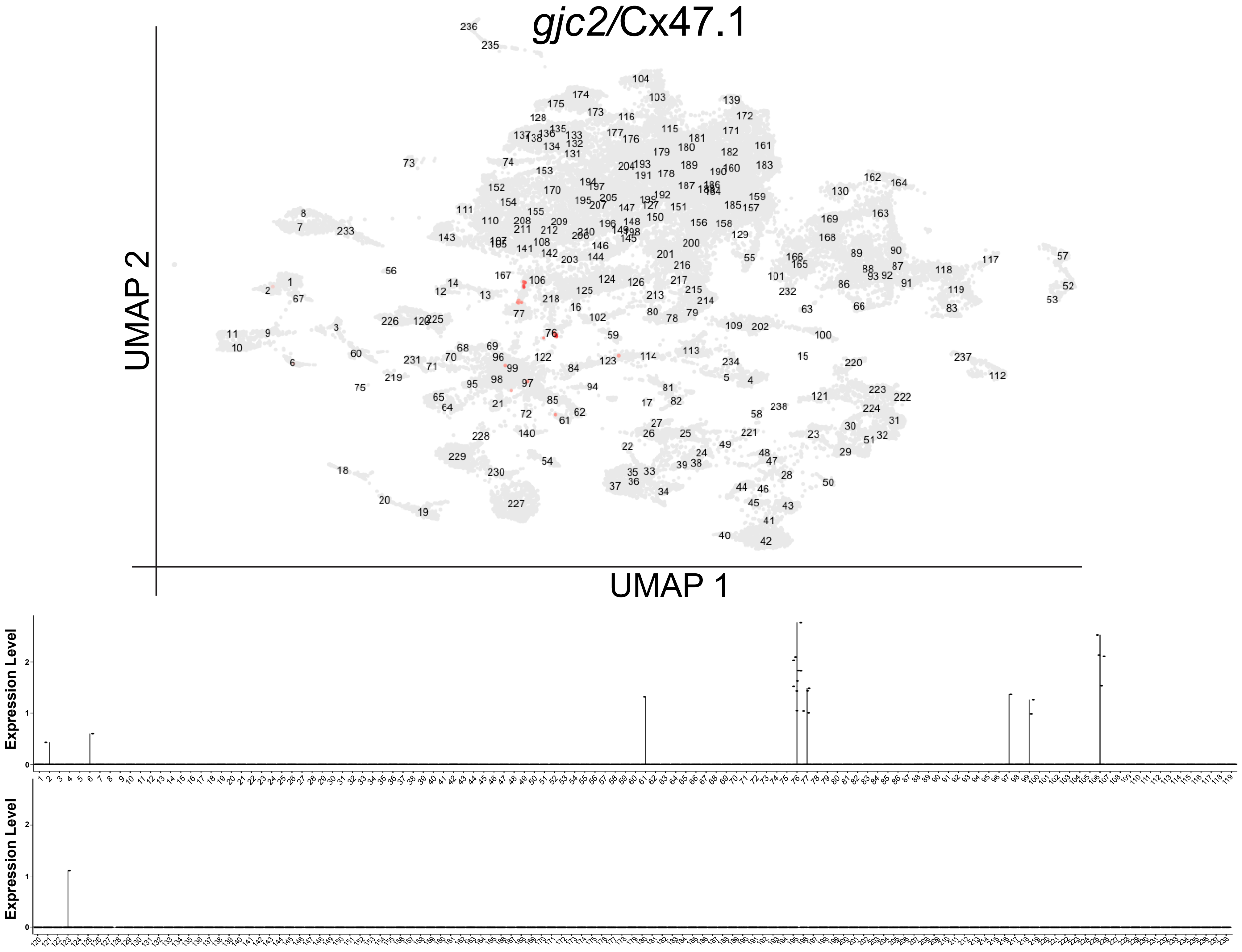

CC

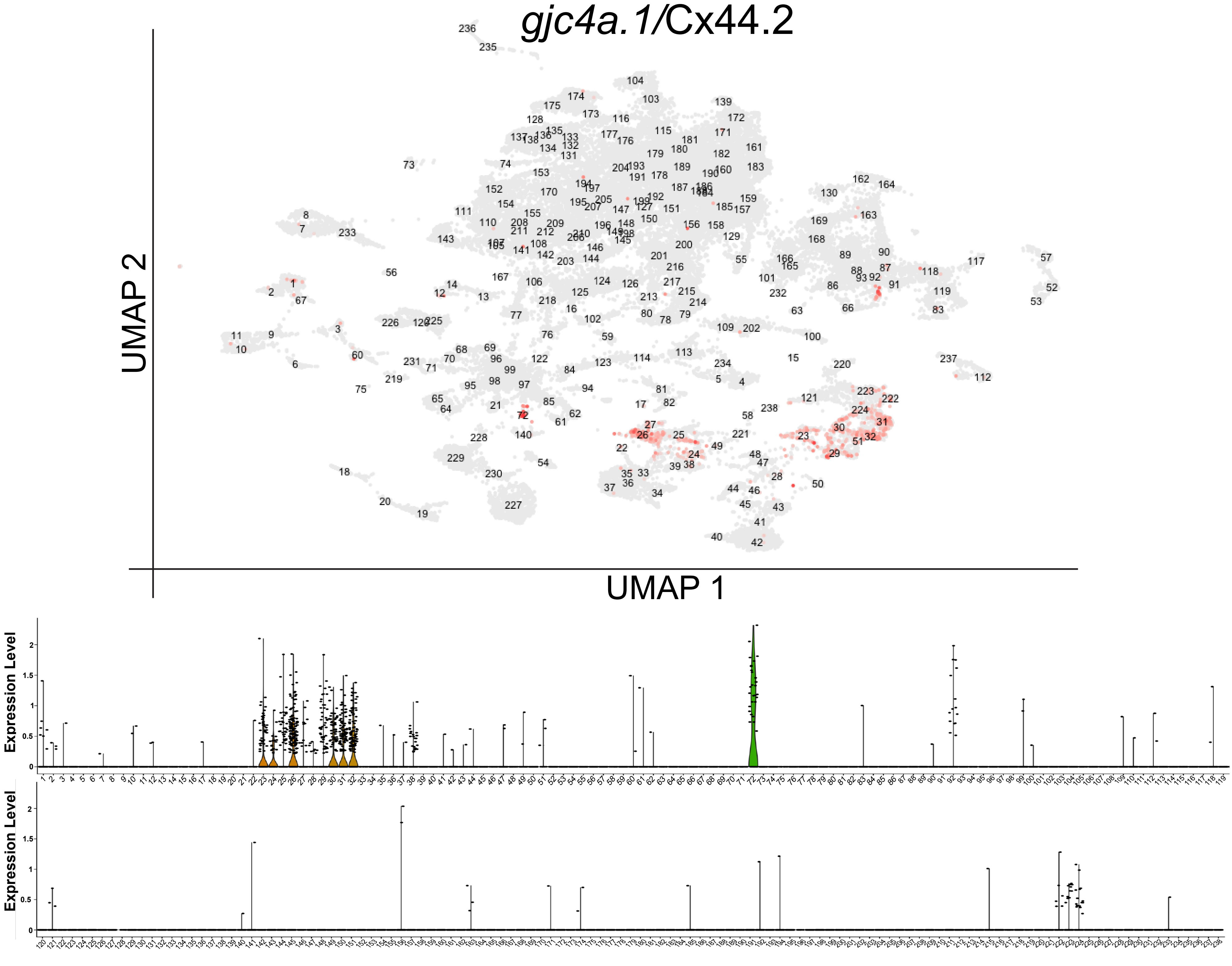
gjc4a.1/Cx44.2
UMAP 2
UMAP 1
Expression Level
Expression Level

DD

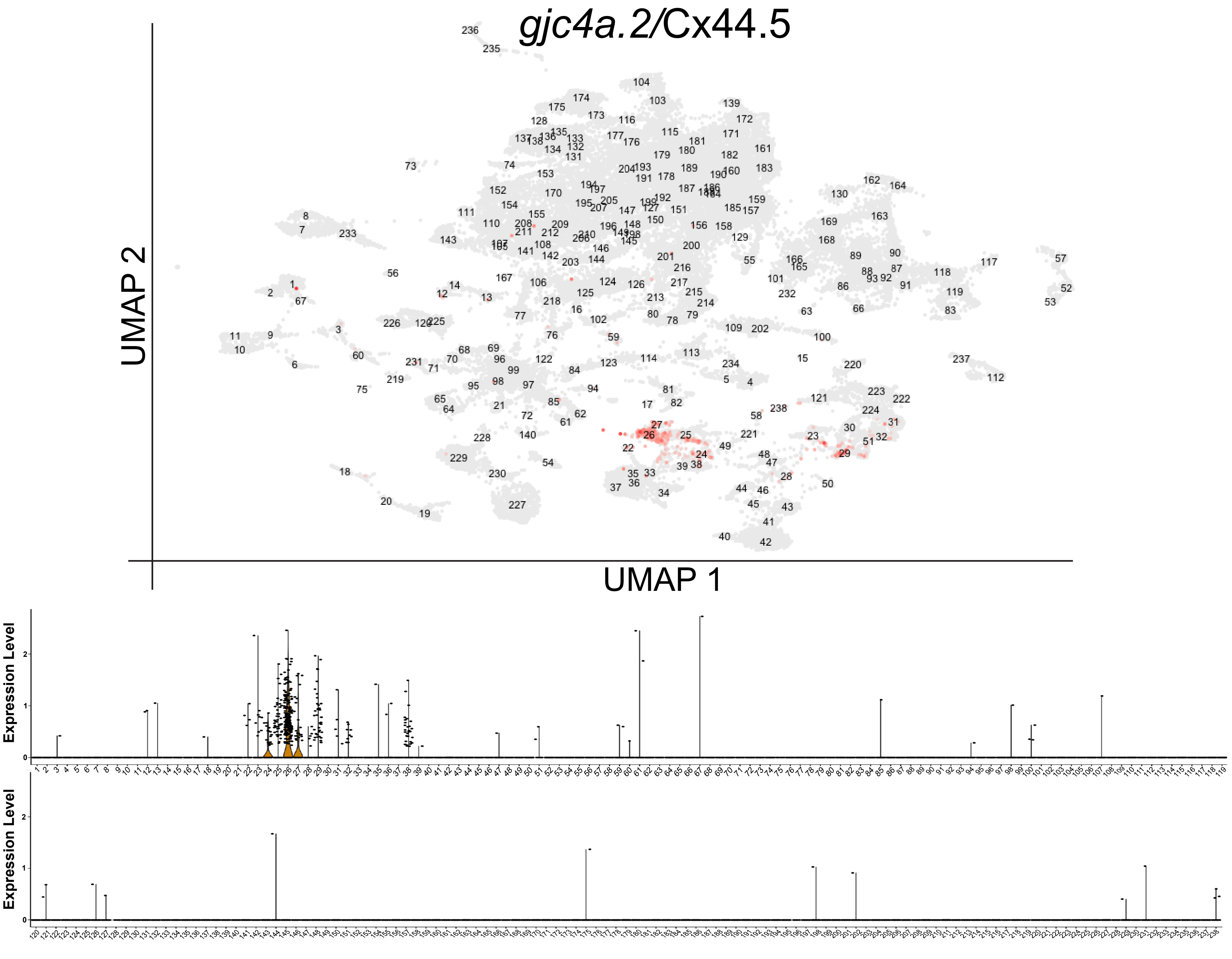

gjc4a.2/Cx44.5
UMAP 2
UMAP 1
Expression Level
Expression Level

EE

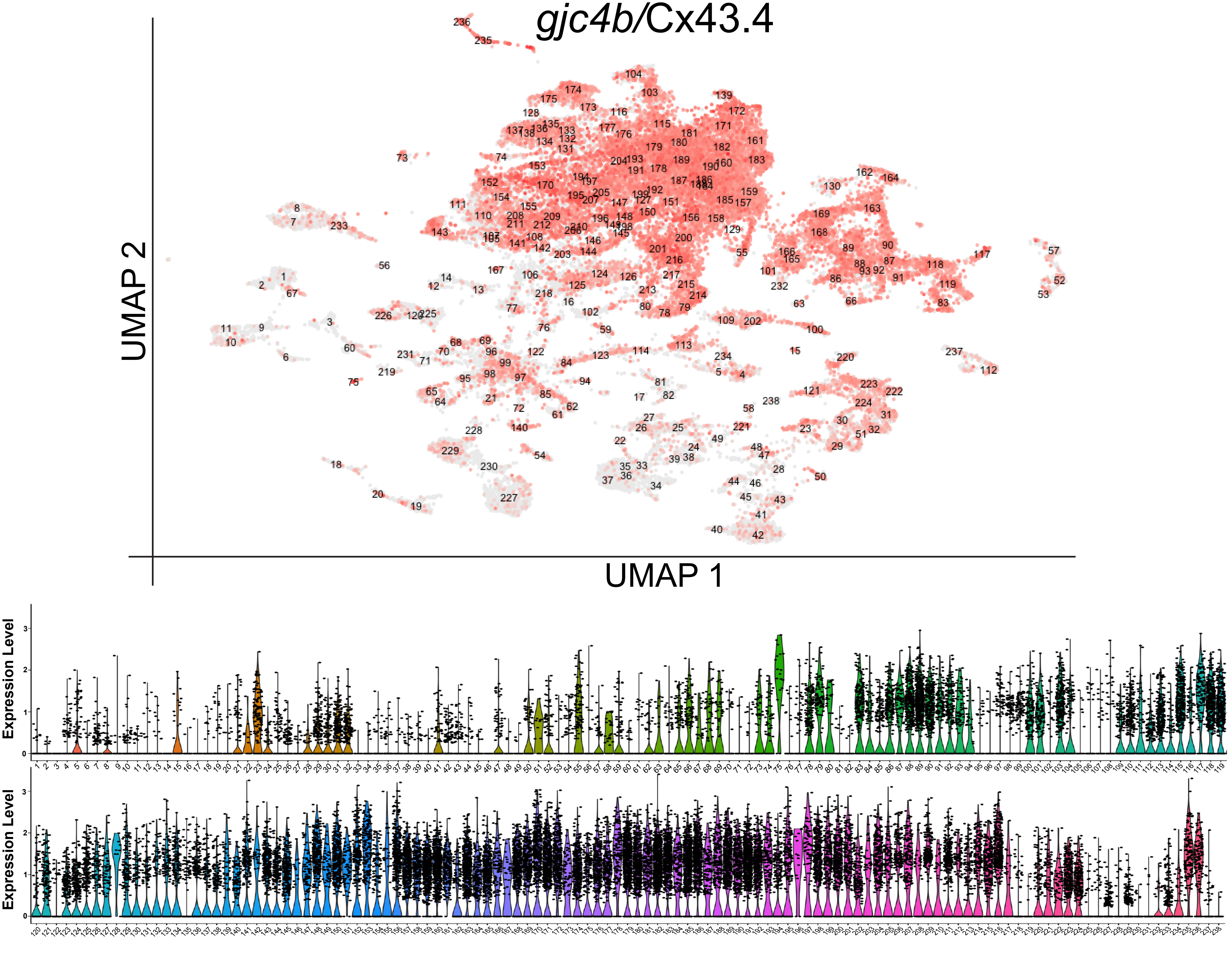

gjc4b/Cx43.4
UMAP 2
UMAP 1
Expression Level
0
1
2
3
Expression Level
0
1
2
3

FF

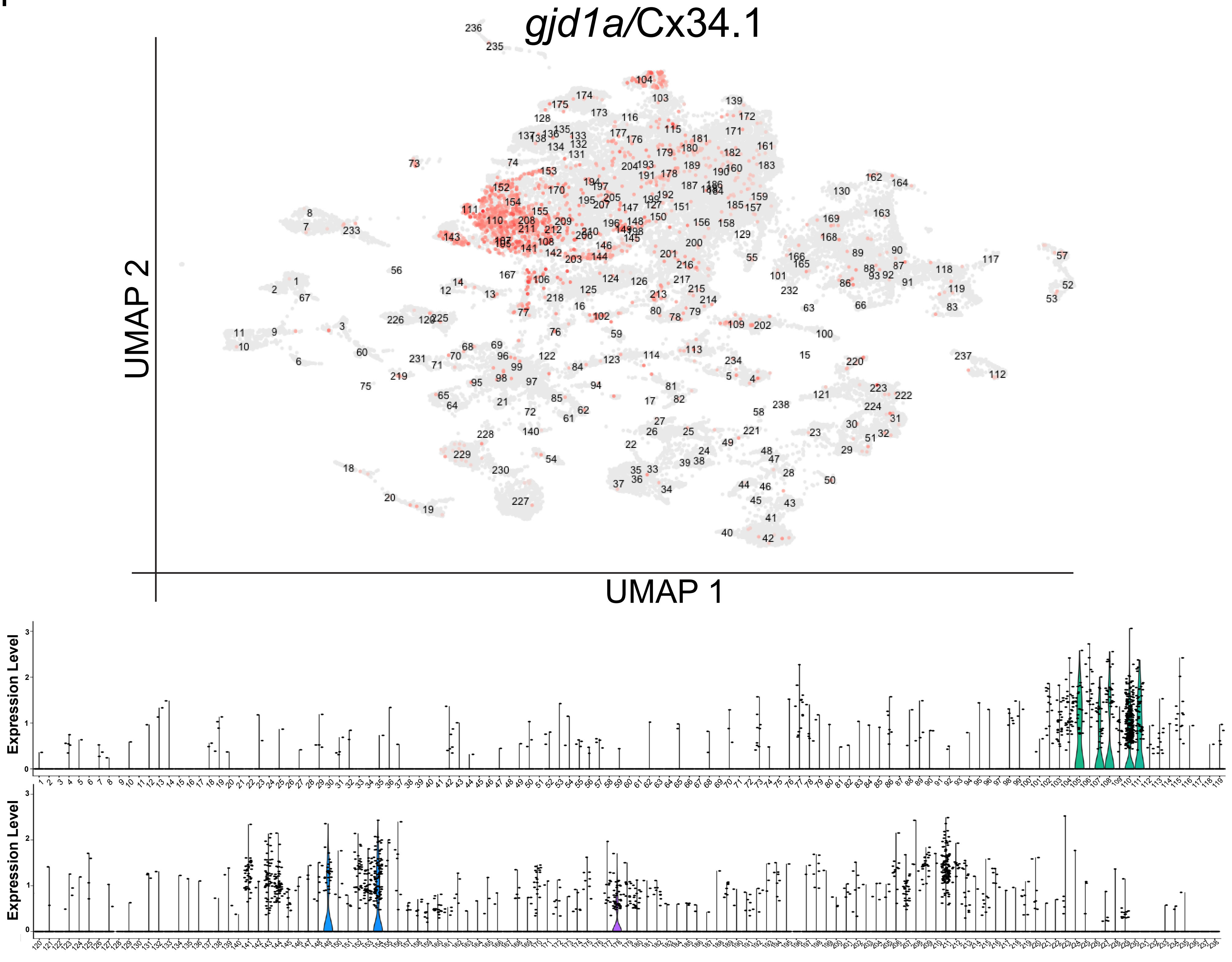

*gjd1a*/Cx34.1
UMAP 2
UMAP 1
Expression Level
Expression Level

GG

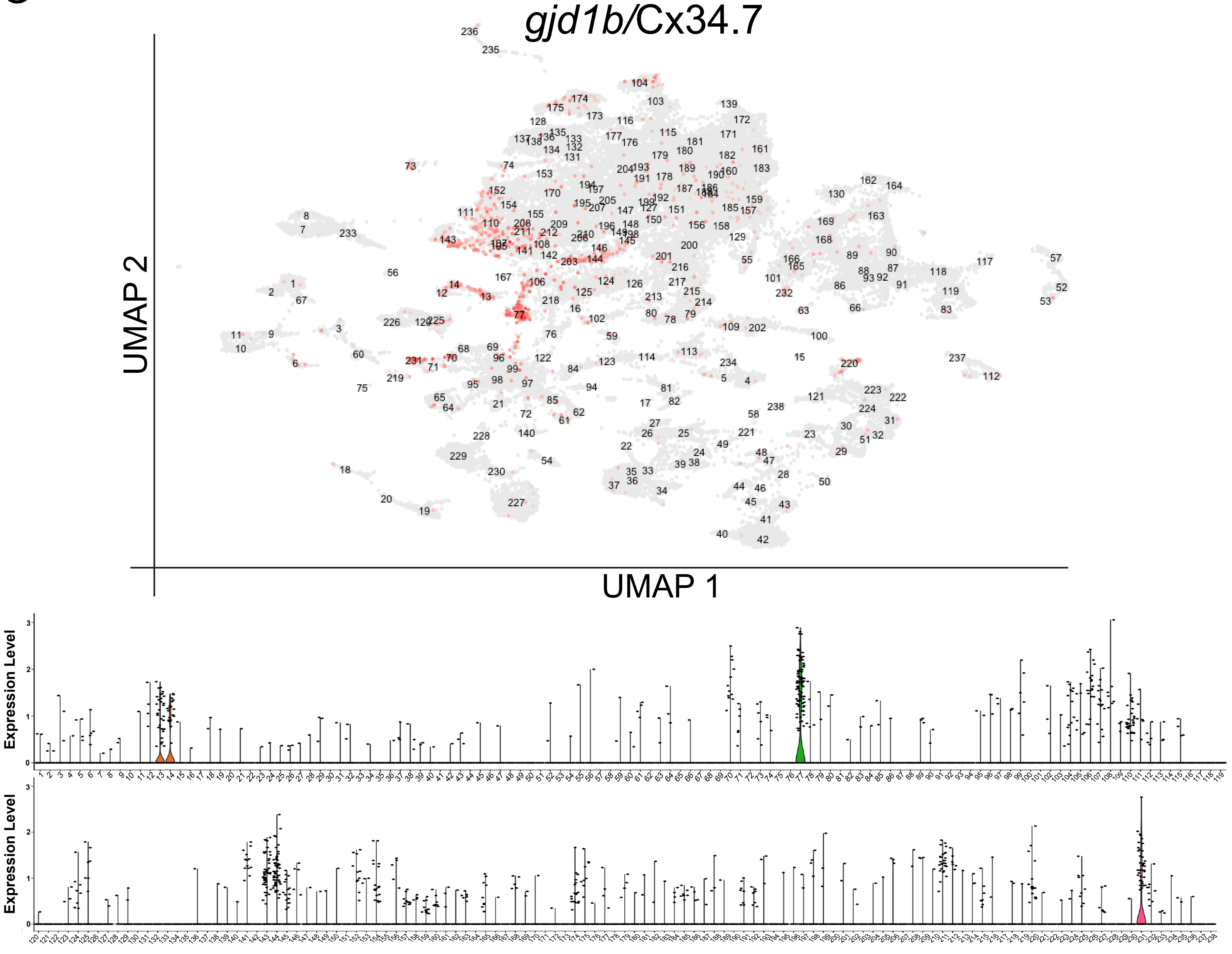

gjd1b/Cx34.7
UMAP 2
UMAP 1
Expression Level
Expression Level

HH

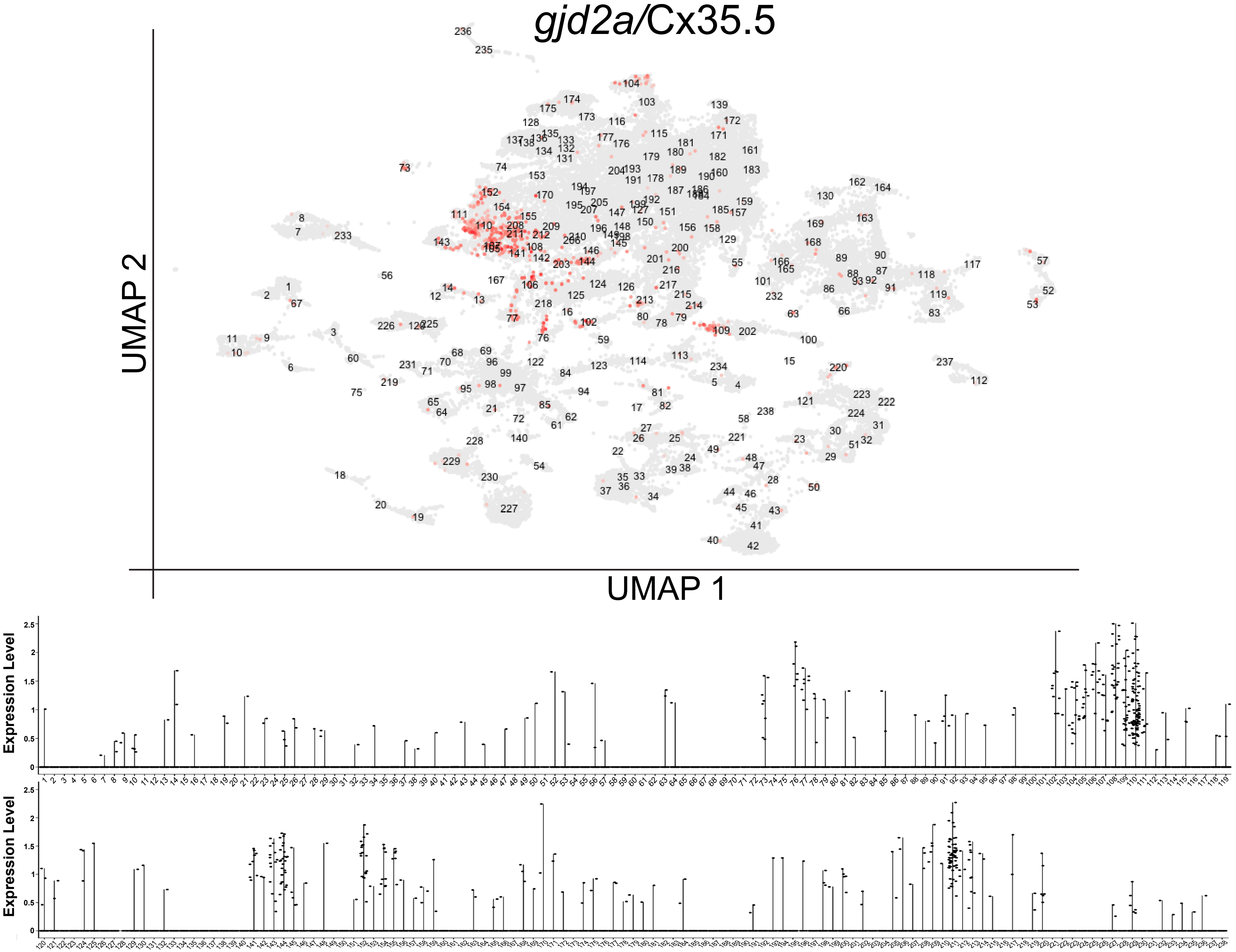
gjd2a/Cx35.5
UMAP 2
UMAP 1
Expression Level
Expression Level

II

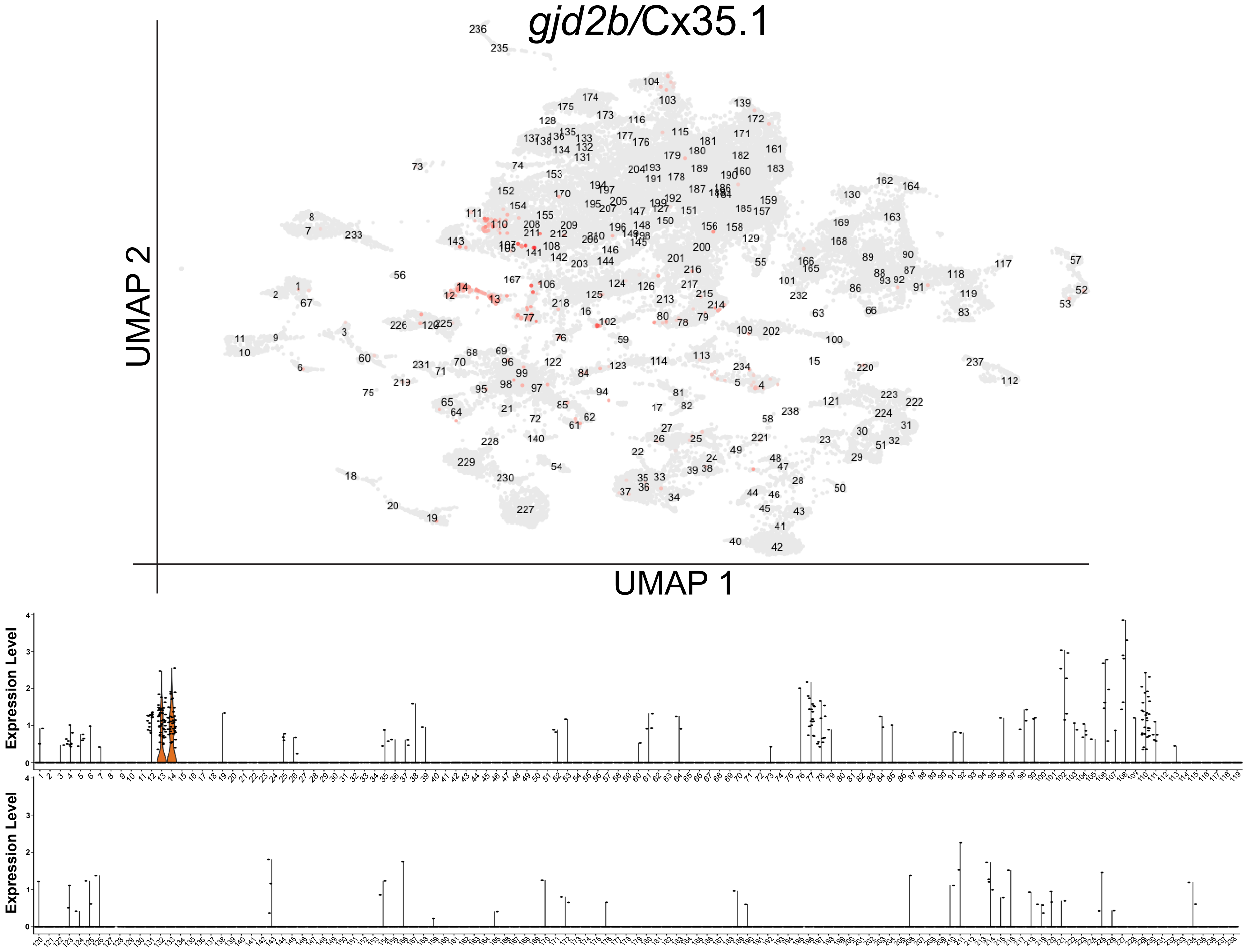
gjd2b/Cx35.1
UMAP 2
UMAP 1
Expression Level
Expression Level

JJ

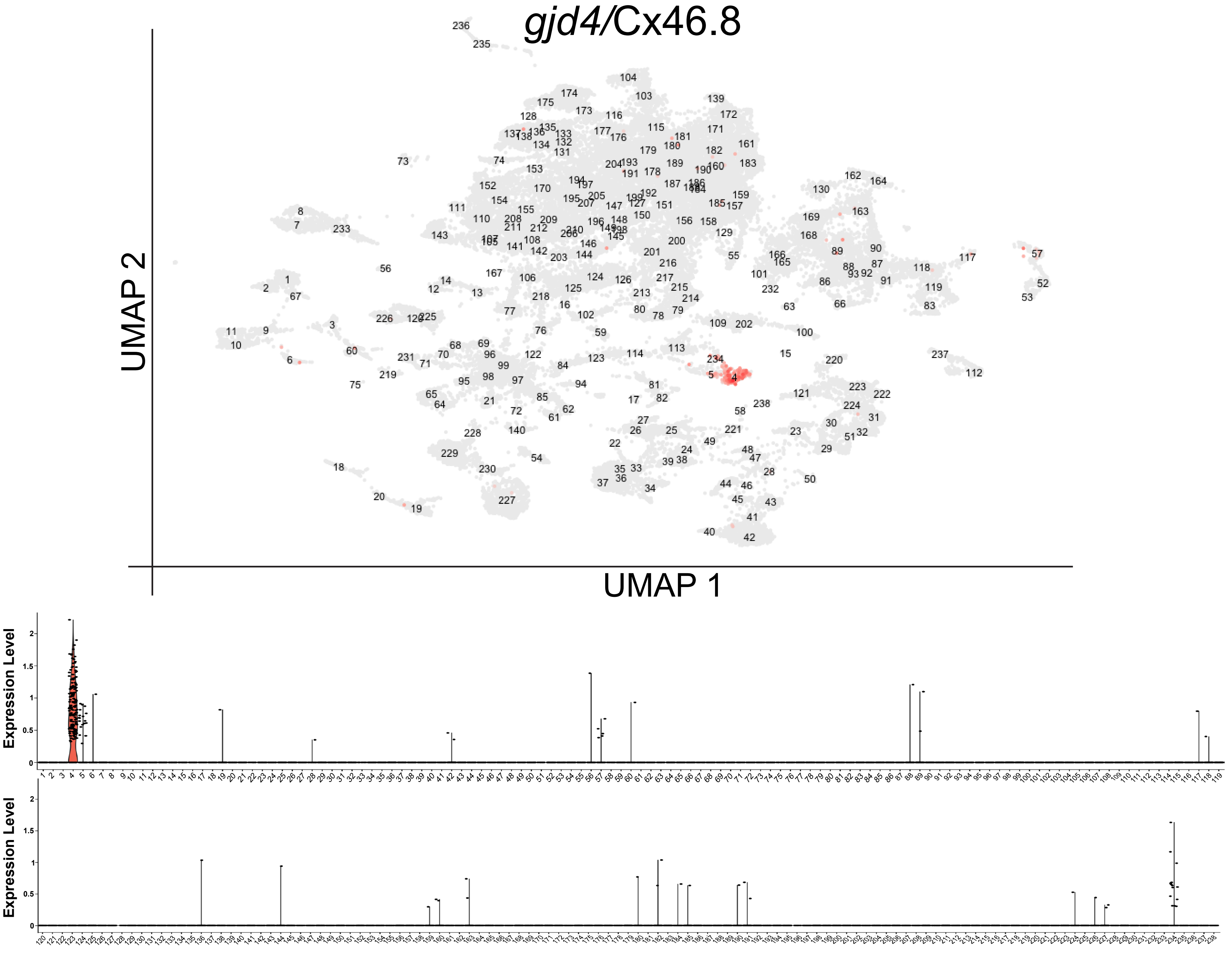

KK

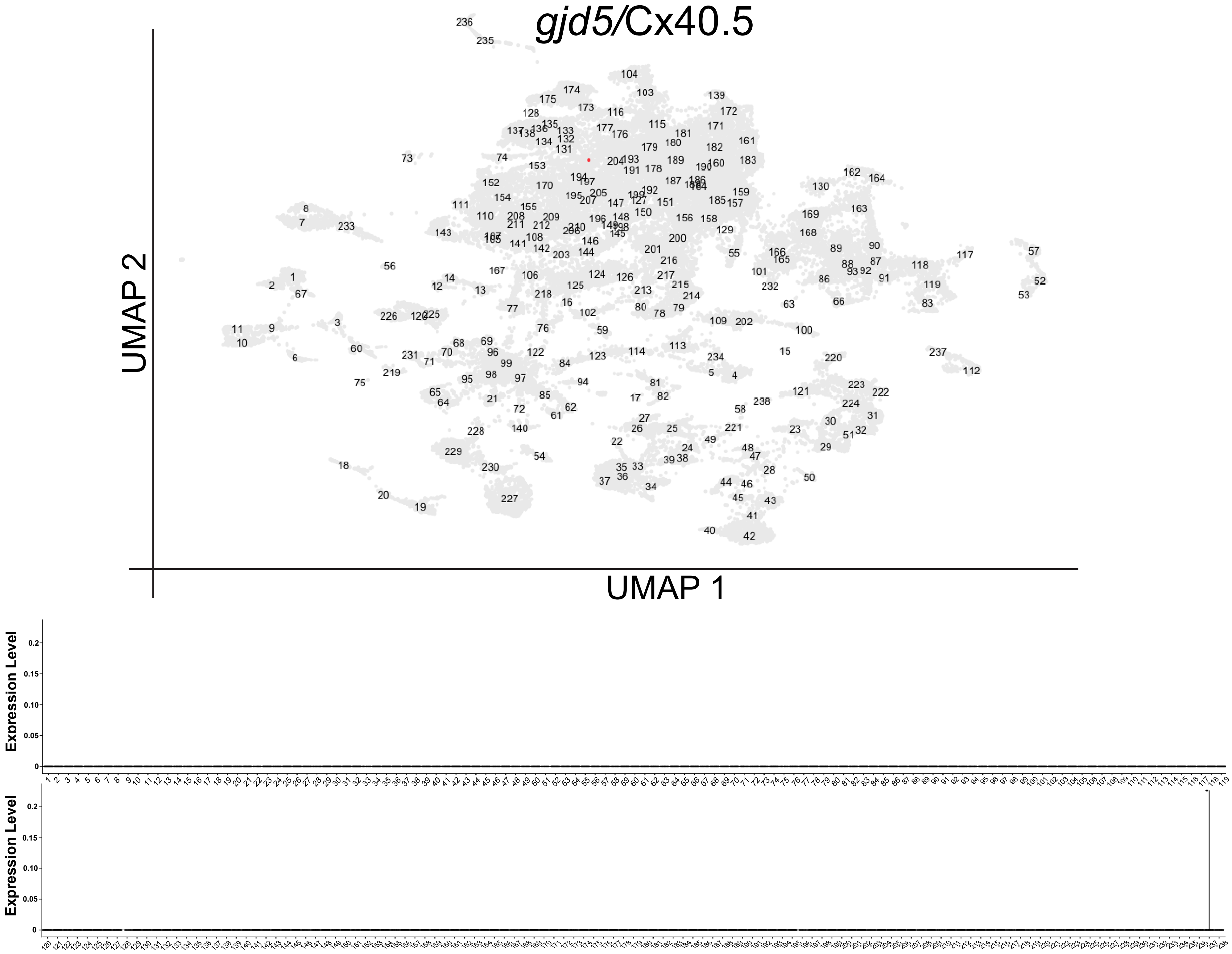

LL

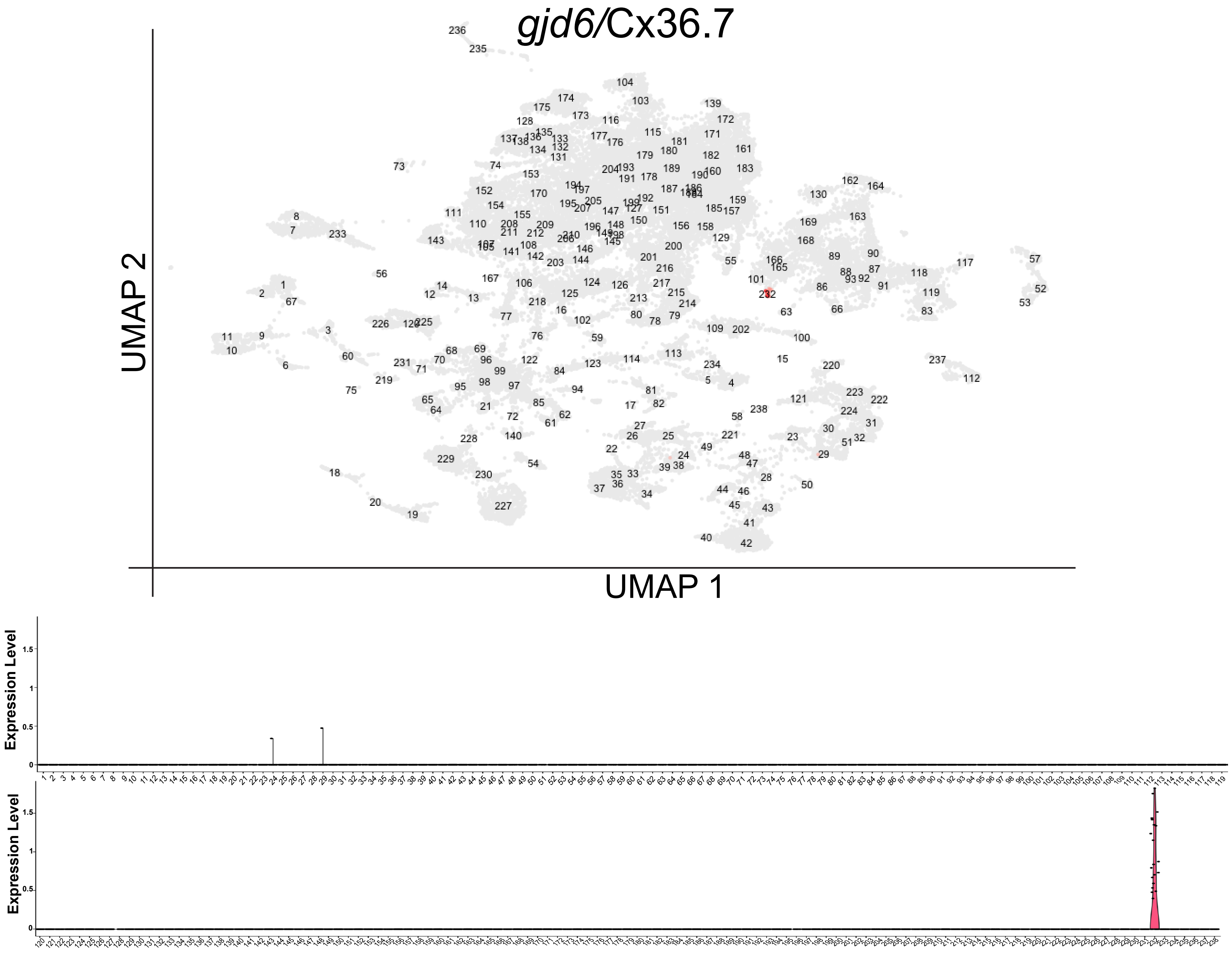

MM

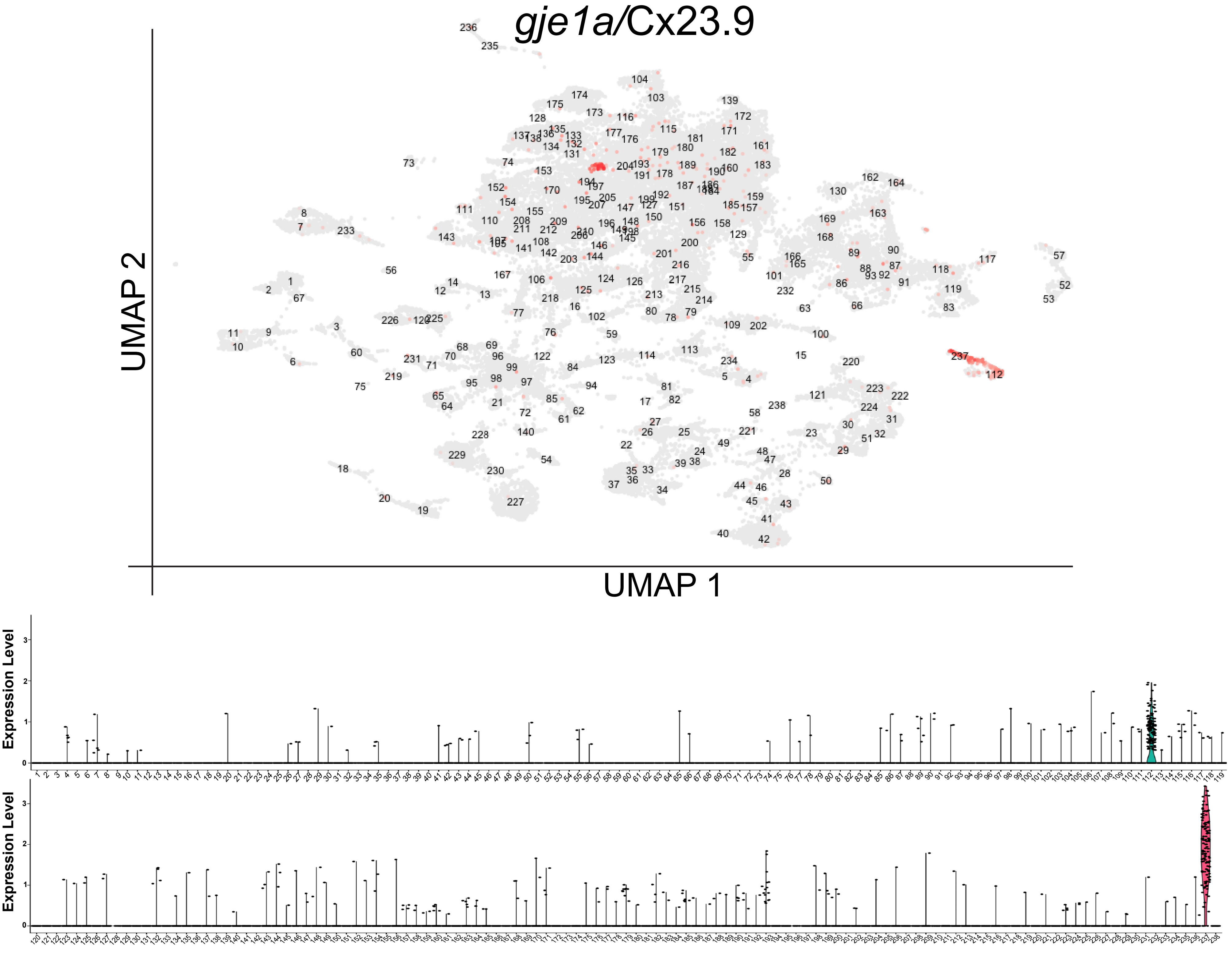

*gje1a*/Cx23.9
UMAP 2
UMAP 1
Expression Level
Expression Level

NN

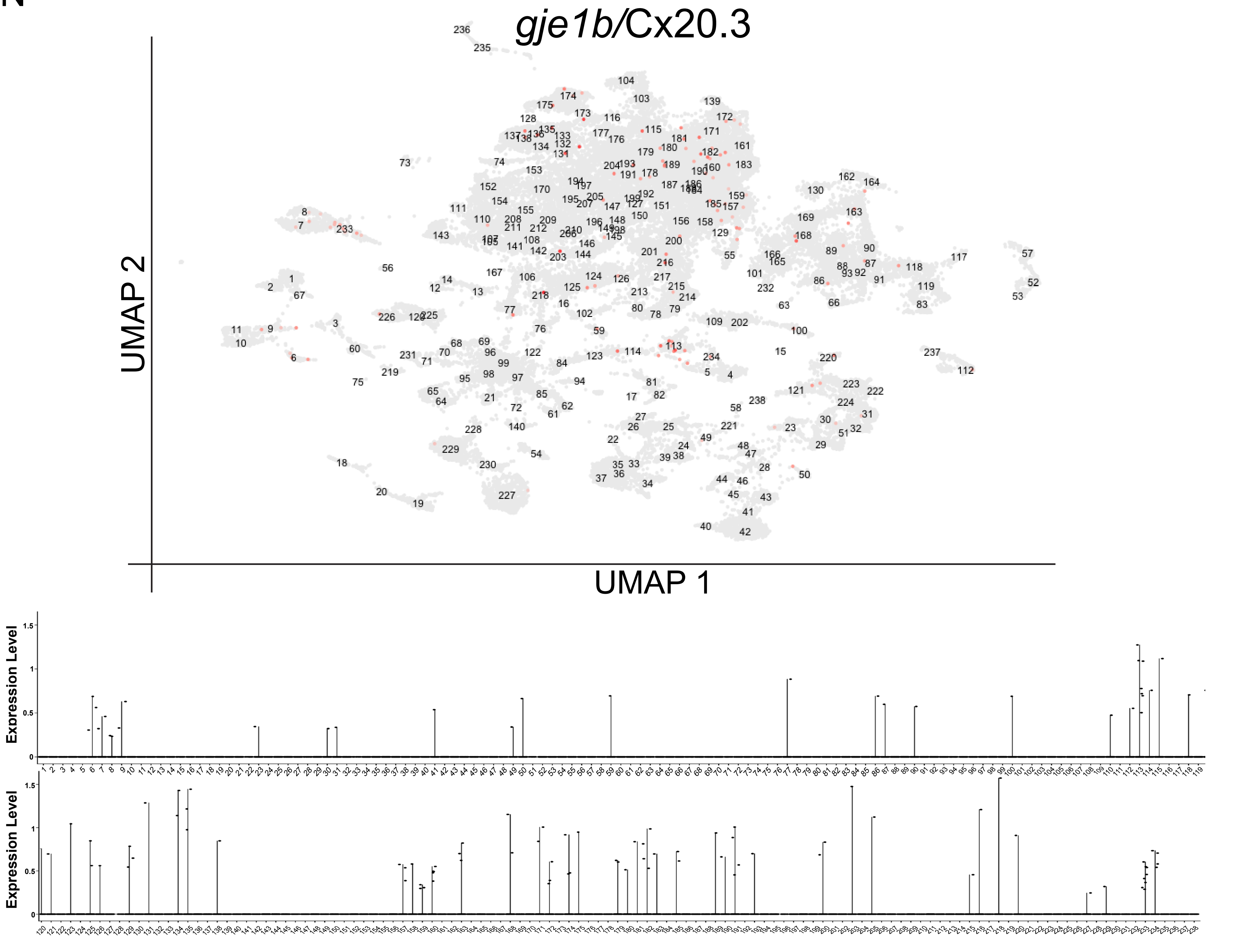
*gje1b*/Cx20.3
UMAP 2
UMAP 1
Expression Level
Expression Level

OO

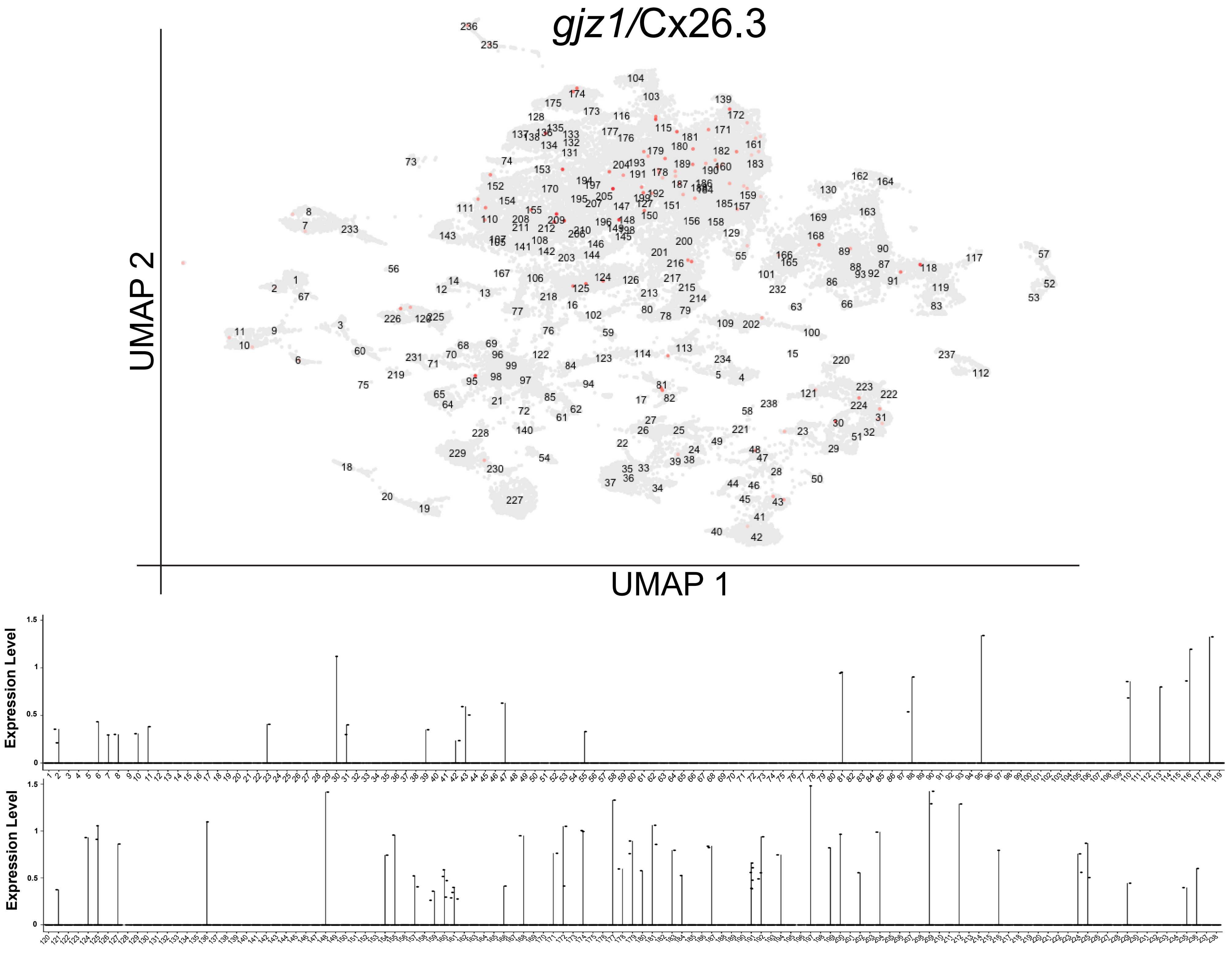

Supplemental Figure 5

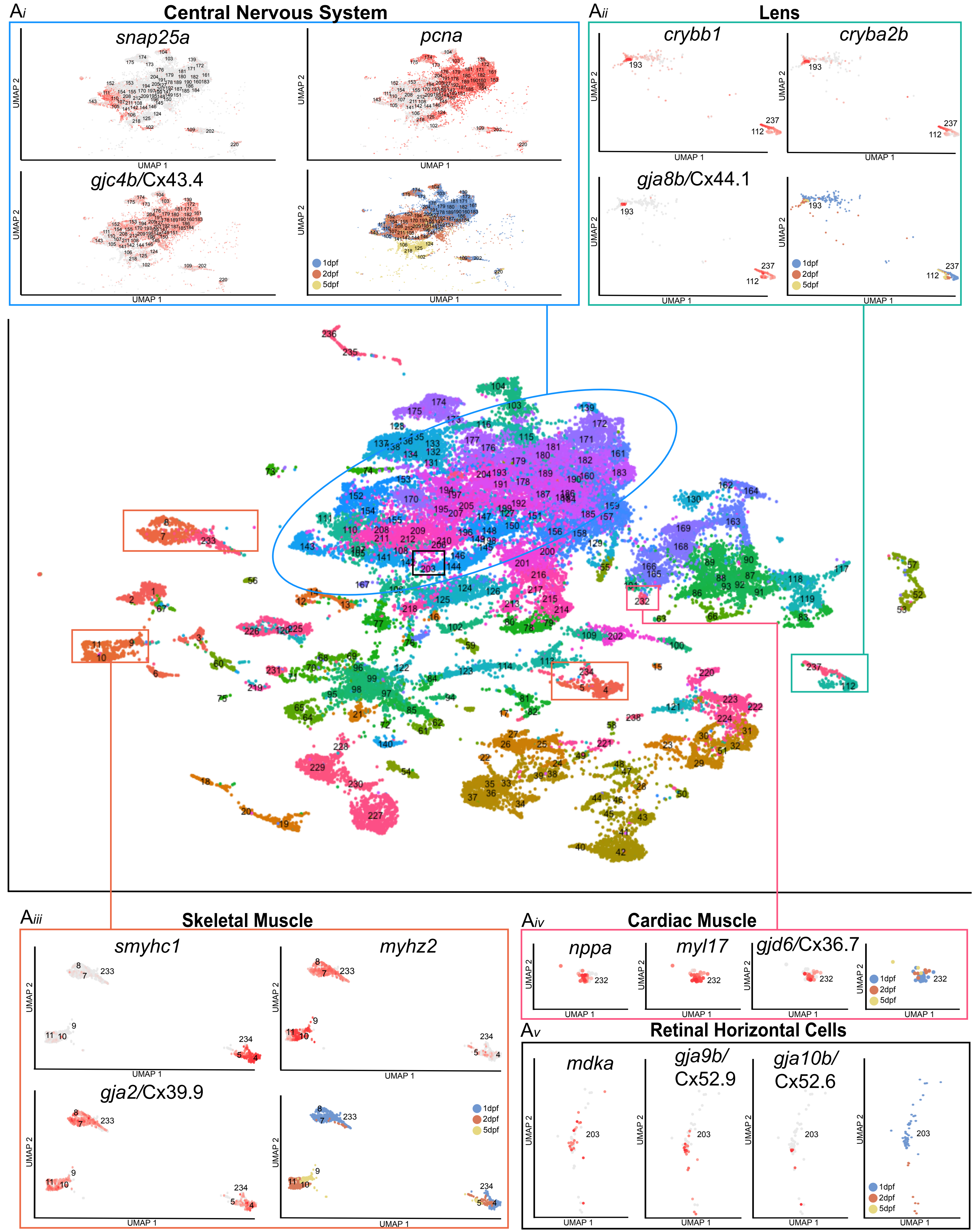

Supplemental Figure 6

A*i*

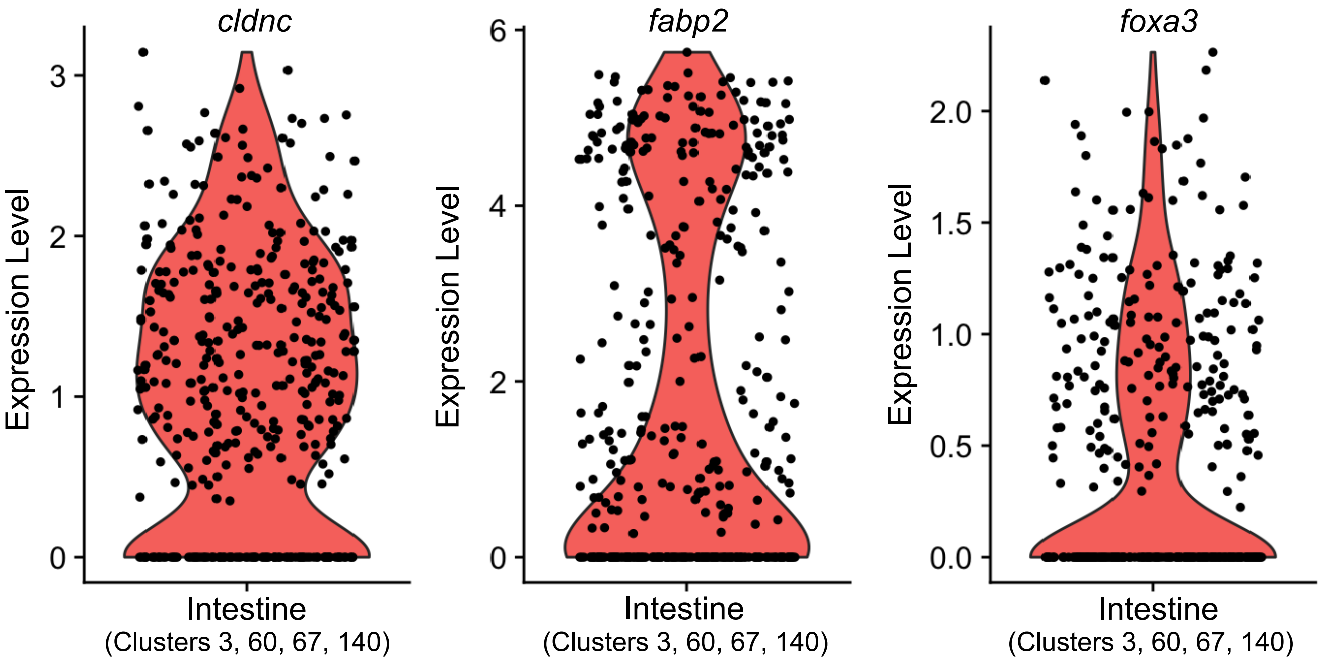


A*ii*

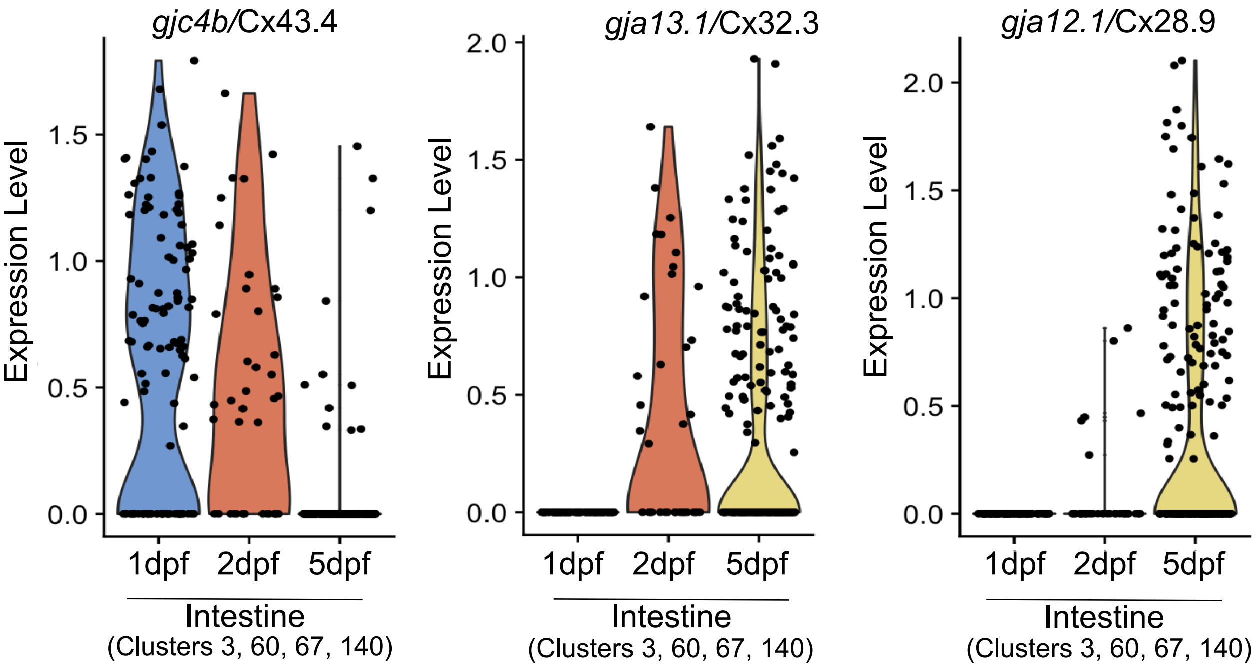

Supplemental Figure 7

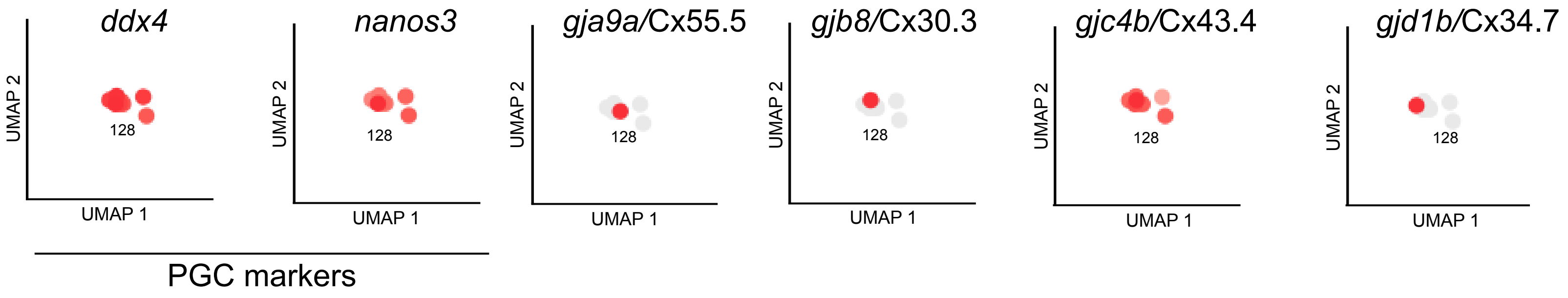

Supplemental Figure 8

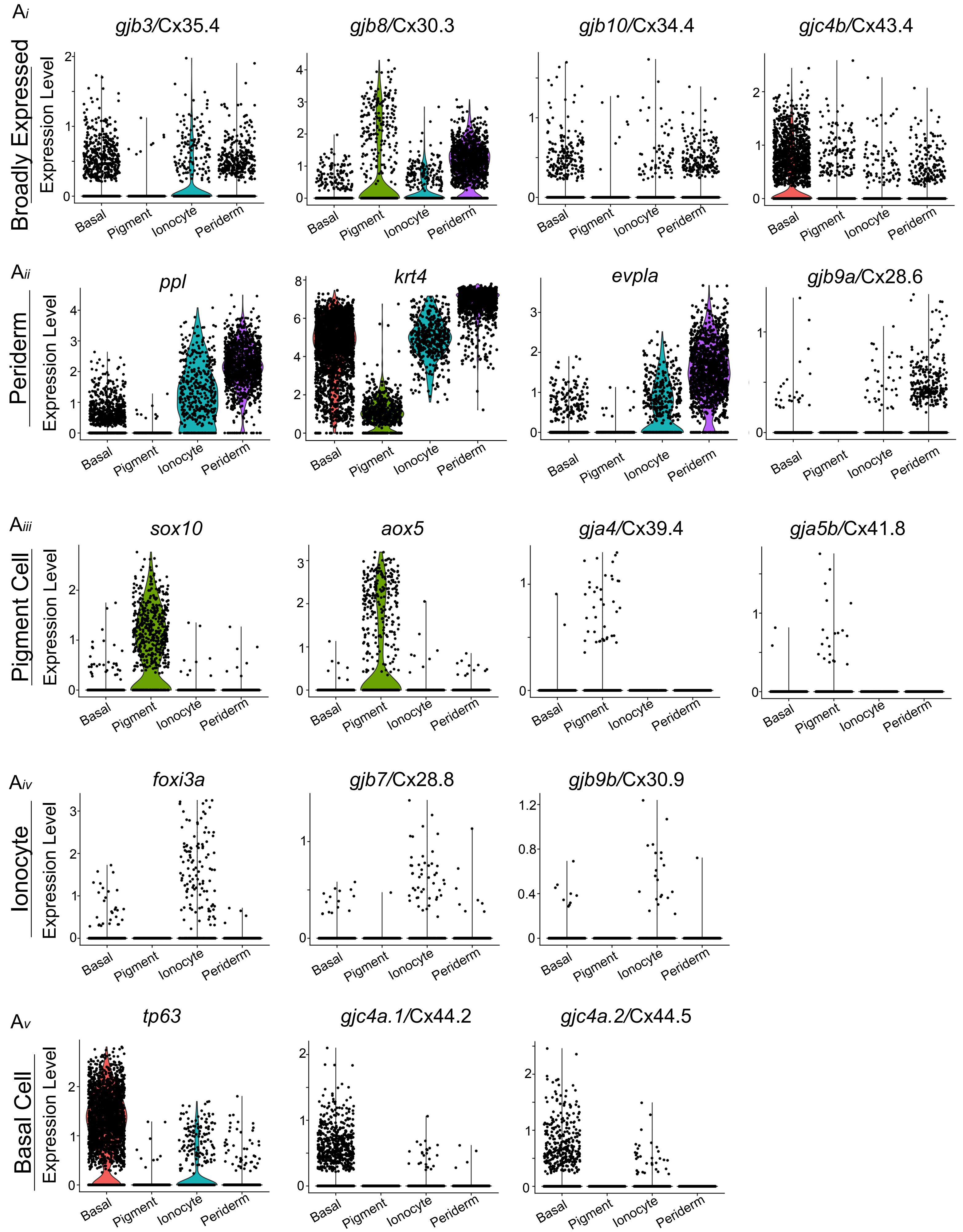
